# Laboratory resistance evolution discovers virulence risks of next-generation antibiotics

**DOI:** 10.64898/2026.07.20.739514

**Authors:** Petra Szili, Márton Simon Czikkely, Zoltán Farkas, Lejla Daruka, Blanka Toldi, Eszter Kurkó, András Zoltán Vonyó, Milán Csernyák, Regina Kámán, Elvin Maharramov, Andreea Daraba, Barbara Benedek, Gergely Imre, Ildikó Ilona Lantos, Léna Mészáros, Áron Somogyi, Dóra Nagy, Zsolt Ferenc Nagy, Gábor Grézal, Eszter Ari, Norbert Kada, Balázs Papp, Zsolt Csenki-Bakos, Edit Kaszab, Balázs Kriszt, István Szabó, Gábor Balogh, Mária Péter, Imre Gombos, Ana-Maria Pilbat, Zsolt Török, Zsófia Varga, Zsolt Czimmerer, Bálint Csörgő, Dóra Adamecz, Csaba Gergő Papp, Zóra Szilovics, Éva Veres, Attila Gácser, Tamara Madácsy, József Maléth, Ferhan Ayaydin, Attila Farkas, Roland Tengölics, Balint Kintses, Valentin Varga, Lajos Haracska, Szilvia Juhász, Csaba Pál

**Author notes:** Corresponding authors. &. These authors contributed equally (Petra Szili, Márton Simon Czikkely, Zoltán Farkas, Lejla Daruka).

## Abstract

Several new antibiotics target multidrug-resistant pathogens, yet resistance is still evaluated mainly by drug-susceptibility, leaving consequences for bacterial pathogenicity poorly understood. Here, we integrate laboratory evolution, genomic surveillance and host–pathogen phenotyping to classify antibiotics by resistance potential and pathogenic consequences. Applying this framework to *Klebsiella pneumoniae* identified functionally distinct antibiotic candidates associated with elevated virulence risk, including host-derived and machine-learning-designed antimicrobial peptides. Laboratory evolution outside the host-environment showed that resistance can rapidly increase virulence, without direct selection for pathogenicity. Despite distinct genetic routes evolving across antibiotic treatments, resistance converged on cell-envelope rewiring. A single, clinically relevant resistance mutation was sufficient to increase epithelial adhesion, intracellular colonization, macrophage immune-evasion, and tissue persistence in murine infection models, transforming *K. pneumoniae* into a highly invasive and cytotoxic pathogen. These findings establish resistance-driven virulence as an underappreciated translational hazard and call for incorporating host–pathogen interactions into resistance surveillance and preclinical antibiotic development.

## Introduction

Antimicrobial resistance is a major threat to global health and a central challenge for antibacterial drug development^1^. Resistance is typically evaluated through its effect on bacterial growth in the presence of antibiotics, most commonly by minimum inhibitory concentration measurements. However, resistance mutations can also alter bacterial physiology in ways that affect host colonization, immune evasion, tissue invasion and disease severity^2–6^. These host-relevant consequences remain poorly integrated into resistance surveillance and preclinical antibiotic evaluation.

A prevailing assumption is that resistance mutations frequently impose fitness costs that attenuate virulence, transmission or persistence when antibiotics are absent^7–10^. Although such trade-offs occur in some systems, resistant pathogens often retain high pathogenic potential, and in some cases resistance evolution has little detectable cost^11,12^. A further possibility is that resistance evolution may actively enhance virulence. If resistance-associated mutations increase infectivity, immune evasion or persistence within host tissues, resistant strains could remain clinically problematic even when antibiotic pressure is reduced^13,14^. Understanding how resistance reshapes host–pathogen interactions is therefore essential for predicting the clinical consequences of resistance evolution.

This question is especially urgent for antibiotics recently introduced into clinical practice or currently in development. Many next-generation antimicrobials target the Gram-negative cell envelope, including membrane-active compounds, host-derived antimicrobial peptides and machine-learning-designed peptide antibiotics. Resistance to these agents may remodel bacterial surface properties in ways that alter host recognition, immune activation and tissue interaction^15–18^. Thus, susceptibility-based resistance profiling alone may miss clinically relevant changes in virulence.

Most evidence linking resistance and virulence remains indirect. Prior studies have largely relied on clinical isolate correlations, individual resistance determinants, or co-occurrence of resistance and virulence genes in genomes and microbiomes^2,19^. These approaches cannot readily determine whether increased virulence is a direct evolutionary consequence of resistance or instead reflects horizontal gene transfer, clonal expansion or shared ecological selection. Experimental systems in which resistance evolves without direct selection for pathogenicity are therefore needed to define whether resistance evolution itself can generate more virulent pathogens.

Here, we combine laboratory resistance evolution, bacterial genomic surveillance and host–pathogen phenotyping to identify antibiotic candidates whose resistance pathways may generate hypervirulent or immune-evasive variants. Laboratory evolution defines the resistance trajectories selected by candidate antibiotics, including shifts in antibiotic susceptibilities and resistance-associated genotypes. Genomic surveillance then establishes whether laboratory-selected mutations, particularly those affecting virulence-associated genes also occur in clinical isolates and high-risk bacterial lineages, allowing to identify mutations with plausible clinical relevance. Initial screening in *G. mellonella* and zebrafish provides scalable preclinical models to prioritize resistant derivatives with elevated host-killing phenotypes. Finally, host–pathogen phenotyping tests whether prioritized resistant mutants alter epithelial adhesion, intracellular colonization, cytotoxicity, immune activation or tissue persistence *in vivo*.

We focus on antimicrobial compounds introduced into clinical practice after 2017 or currently in development, building on previous work that mapped rapid resistance evolution to these drugs in Gram-negative bacteria^20^. We then test whether these resistance trajectories alter host-relevant phenotypes across a multi-scale infection framework, including *Galleria mellonella*, zebrafish, systemic murine infection, *ex vivo* intestinal organoids, and gut and lung epithelial cell models.

We find that laboratory resistance evolution can rapidly increase host-killing capacity in *Klebsiella pneumoniae*. A substantial fraction of adapted lines reached *Galleria mellonella* killing rates comparable to clinical isolates carrying canonical multiple virulence-associated markers, although the evolved lines themselves lacked these markers. These phenotypes are driven by mutations that also occur in clinical isolates and converge on regulatory rewiring of cell-envelope physiology, particularly lipid A remodelling. Functionally, these changes increase epithelial adhesion, intracellular colonization, cytotoxicity, resistance to host antimicrobial peptides, macrophage immune evasion and tissue persistence. A single resistance mutation is sufficient to reproduce many of these phenotypes, demonstrating that virulence can emerge as a pleiotropic by-product of resistance evolution. Together, our results define a practical translational risk for antibiotic development: resistance to candidate antimicrobials can generate bacterial variants that are both less susceptible to treatment and more damaging to the host.

## Results

### Antibiotic adaptation targets virulence genes

We hypothesized that the evolution of antibiotic resistance can influence virulence because it often targets pathways shared between antibiotic stress responses and host–pathogen interactions. Using VirulentHunter^21^, a deep-learning framework trained on curated virulence factor databases, we identified virulence-associated genes in *K. pneumoniae* and *E. coli* and analyzed whole-genome sequencing data from 176 lines evolved under 15 antibiotics (see Methods, Table 1). Mutations in virulence-associated genes were modestly but significantly enriched, with 30.5% of mutated genes in *E. coli*, and 22.4% in *K. pneumoniae* (1.77-fold, permutation test, *P* < 1×10⁻⁴), and with variation across antibiotic classes (Poisson generalized linear model heterogeneity test, *P* < 0.001). In contrast, control lines evolved under identical laboratory conditions without antibiotics showed no enrichment (permutation test, *P* = 0.620), indicating that this pattern is driven by antibiotic selection rather than laboratory adaptation^20^.

**Table 1.** Antibiotics used for laboratory evolution. The table lists antibiotics used for experimental evolution of bacterial strains^20^, together with their abbreviations, antibiotic classes, primary molecular targets, and current clinical status. For approved compounds, the year of first U.S. Food and Drug Administration (FDA) approval is indicated. “Marketed; long-established” refers to clinically approved antibiotics with long-standing use for which the original approval date is less commonly specified. “Phase II-ready” indicates compounds that successfully completed Phase I clinical development and were prepared for Phase II initiation, while “Preclinical” refers to compounds that have not yet entered human clinical trials. For dual-target compounds, both principal molecular targets are listed.

| Antibiotic | Abbreviation | Antibiotic class/target |  | Date of approval/clinical phase |
|---|---|---|---|---|
| Omadacycline | OMA | tetracyclines |  | FDA approved (2018) |
| Eravacycline | ERA |  |  | FDA approved (2018) |
| Doxycycline | DOX |  |  | FDA approved (marketed; long-established) |
| Delafloxacin | DEL | topoisomerase inhibitors |  | FDA approved (2017) |
| Gepotidacin | GEP |  |  | FDA approved (2025) |
| Zoliflodacin | ZOL |  |  | FDA approved (2025) |
| Moxifloxacin | MOX |  |  | FDA approved (1999) |
| Apramycin | APR | aminoglycosides |  | Phase II candidate |
| Gentamicin | GEN |  |  | FDA approved (marketed; long-established) |
| Sulopenem | SUO | carbapenems |  | FDA approved (2024) |
| Meropenem | MER |  |  | FDA approved (1996) |
| Tridecaptin M152-P3 | TRD | membrane targeting antibiotics | Lipid II and lipid A | Preclinical |
| POL-7306 | POL |  | BamA and lipid A | Preclinical |
| SCH79797 | SCH |  | MscL and folate synthesis | Preclinical |
| SPR-206 | SPR |  | Lipid A | Phase II-ready (development discontinued, 2025) |
| Polymyxin B | PMB |  | Lipid A | FDA approved (marketed; long-established) |

Mutated virulence-associated genes were distributed across multiple functional categories, including adhesion, secretion systems, immune modulation, and invasion (Figure S1). Regulatory genes with established links to virulence were recurrently affected in both species (8.66-fold enrichment, permutation test, FDR-corrected *P* < 0.001), particularly the two-component systems *basRS* and *phoPQ*, as well as additional regulators involved in envelope stress and metabolism (e.g., *baeS* in *K. pneumoniae* and *arcA* in *E. coli*). Consistent with these targets, signal transduction genes were the most enriched functional category among mutated loci (3.82-fold enrichment, permutation test, FDR-corrected *P* < 0.001).

To assess clinical relevance, we examined the occurrence of 189 non-synonymous mutations accumulated in virulence-associated genes during laboratory evolution across large genome collections (∼394,900 *E. coli* and ∼72,400 *K. pneumoniae* genomes, Data S1)^22^. After controlling for unequal genome sampling depth (see Note S2), 38–39% of the virulence-associated mutations were detected in natural isolates (Fig. S2), and these mutations were enriched among clinical infection-associated isolates compared with carriage, environmental or other non-infection-associated isolates (Fisher’s exact test, OR = 3.14, *P* < 2.2 × 10⁻¹⁶ for *E. coli* and OR = 1.88, *P* < 2.5 × 10⁻⁷ for *K. pneumoniae*).The mutations were distributed across multiple globally prevalent high-risk clones of *K. pneumoniae* (Figure S3), indicating independent origins^23^. In contrast, mutations arising during adaptation to antibiotic-free medium showed no enrichment in pathogenic isolates (Fisher’s test, OR = 1.04, *P* = 0.798 for *E. coli* and OR = 0.247, *P* = 0.00435 for *K. pneumoniae*). Together, these findings suggest that mutations in virulence-associated genes selected during laboratory resistance evolution also arise in natural populations, including clinically relevant lineages.

### Elevated virulence is frequent and general across antibiotic treatments

To directly investigate how antibiotic resistance shapes virulence, we analyzed 124 adapted lines of *K. pneumoniae* and *E. coli* that had adapted *in vitro* to one of each 16 antibiotics (for information on the lines, see Data S2). We assessed virulence using the invertebrate greater wax moth (*Galleria mellonella*) larval infection model, which enables elevated throughput and reproducible measurements^24–26^. For each adapted line, we quantified relative virulence as the host-killing rate compared to the corresponding ancestor (see Note S3, Fig. SN1).

Remarkably, 24.2% of the 124 adapted lines exhibited increased virulence, whereas 30.6% showed decreased virulence (Fig. 2A). Thus, resistance evolution produced bidirectional effects on pathogenicity, with virulence enhancement being nearly as frequent as virulence impairment in the full dataset. Virulence enhancement was widespread among antibiotic treatments, as 13 out of the 16 antibiotics yielded at least one evolved line with increased virulence (Fig. 2B). Membrane-targeting compounds preferentially selected for increased virulence (Fisher’s exact test with Monte Carlo, simulation, P = 0.02062, Fig 2B), although these antibiotics differ substantially in both structure and mode of action (Table 1, Fig. 1.). Meanwhile, the fraction of lines with increased virulence did not depend on genetic background (Figure S4A-B) and *in vitro* fitness in antibiotic-free medium cannot account for the observed increase in virulence (Figure S5). Overall, the results indicate that resistance-associated changes in virulence depend on the antibiotic class and the physiological route by which resistance evolves.

**Figure 1.**
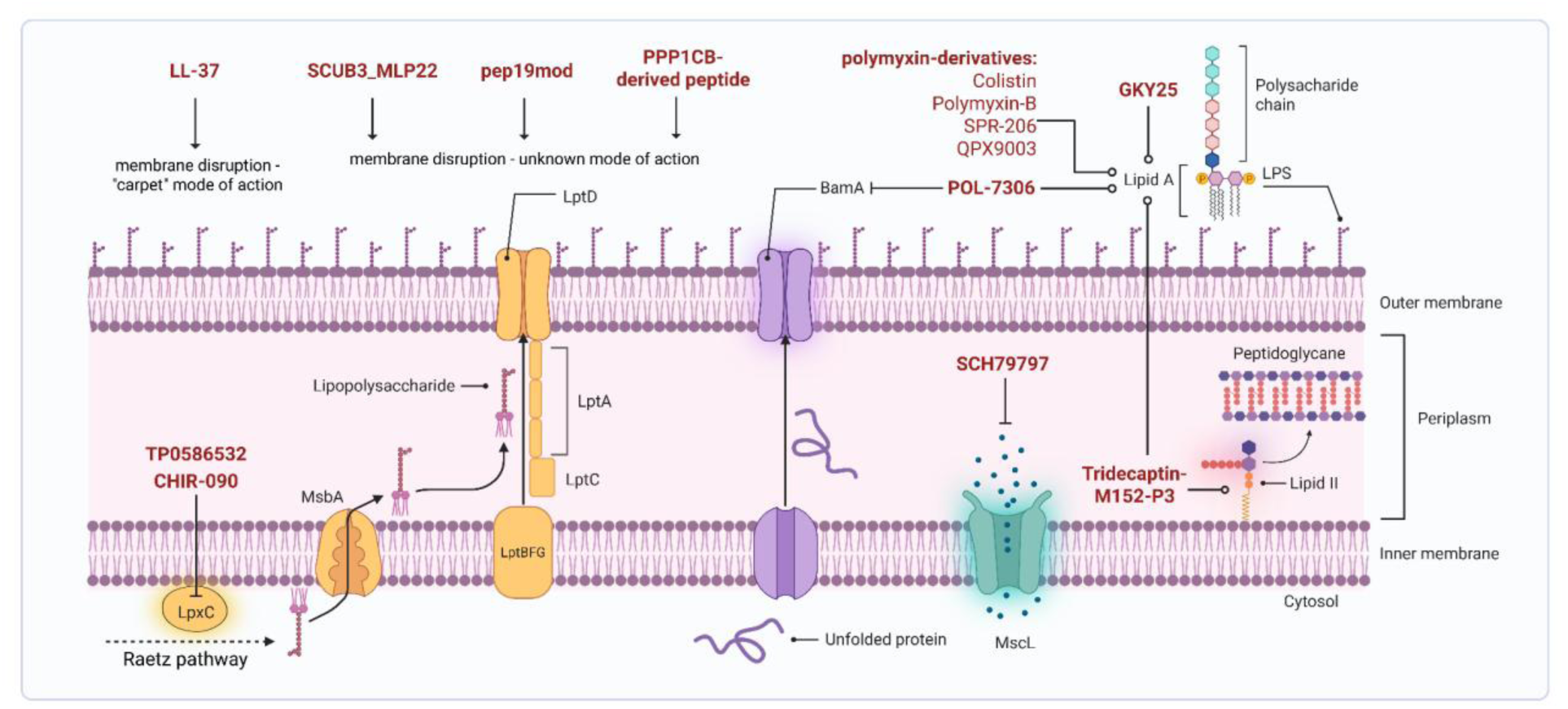
Structurally and functionally distinct antibiotics targeting the Gram-negative cell envelope applied in this study. Schematic representation of the Gram-negative bacterial cell envelope illustrating key pathways involved in lipopolysaccharide (LPS) biogenesis, membrane integrity, and envelope stress, and highlighting the diverse molecular targets of envelope-targeting antibiotics examined in this study. LPS is composed of a membrane-anchored lipid A moiety, a core oligosaccharide, and an O-antigen polysaccharide chain; lipid A serves as a central target for multiple antimicrobial agents due to its essential role in outer membrane stability. LPS is synthesized via the Raetz pathway (including LpxC) at the inner membrane (IM), flipped by MsbA, and transported to the outer membrane (OM) through the Lpt machinery (LptBFGC–LptA–LptD). β-barrel assembly is mediated by BamA. Lipid II, a precursor of peptidoglycan synthesis, resides in the outer leaflet of the IM. The mechanosensitive channel MscL functions as an osmotic safety valve, opening under membrane tension to prevent cell lysis. Blunt-ended inhibitory connectors denote direct suppression of protein function or enzymatic activity, whereas circle-headed connectors indicate molecular binding interactions that initiate downstream effects on membrane structure or envelope-associated processes. The figure was created in BioRender®. (For a more detailed description see, Note S1.)

**Figure 2.**
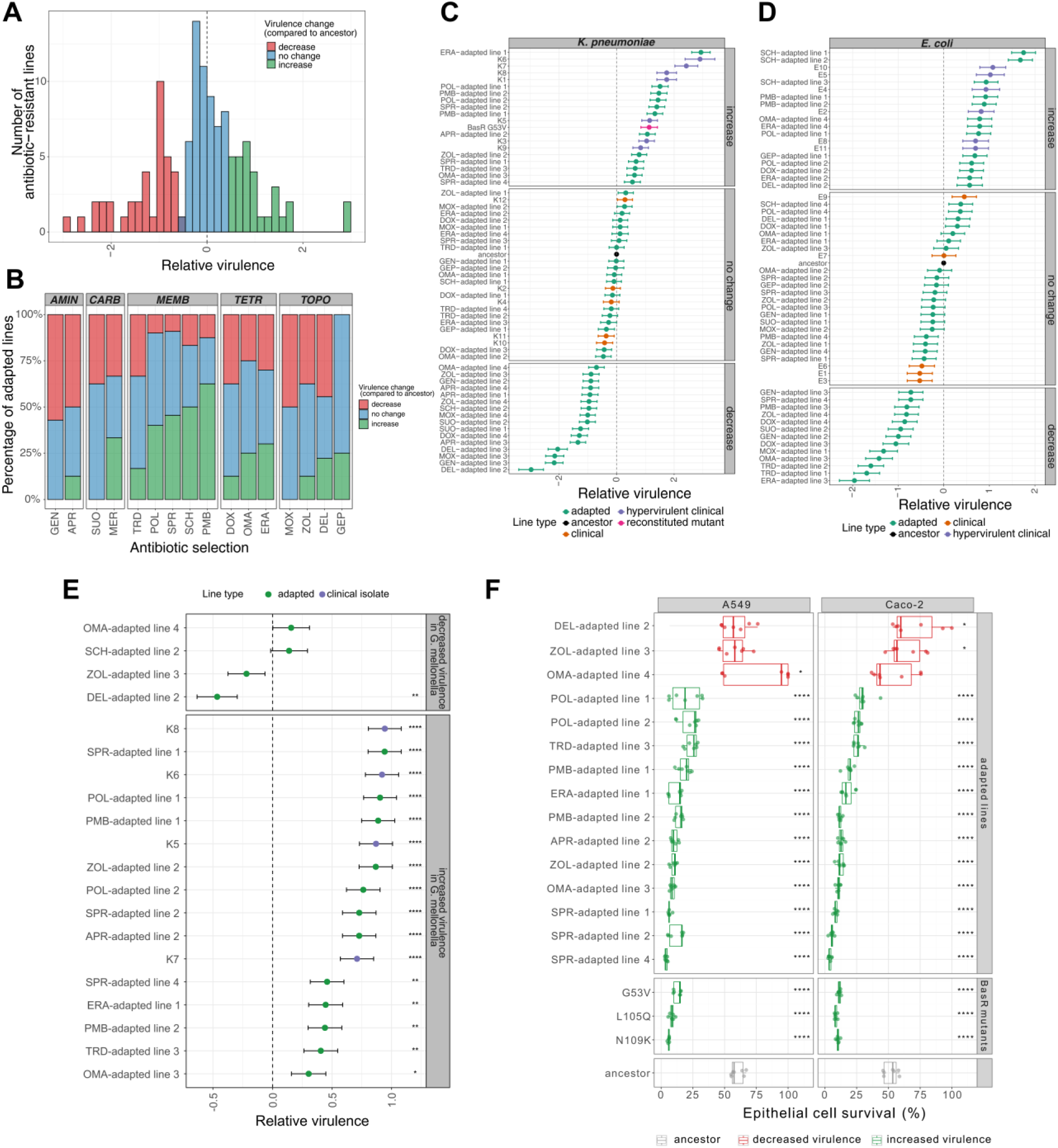
In vivo virulence and in vitro epithelial cytotoxicity of antibiotic-adapted lines. **(A)** Relative virulence distributions for 124 adapted *E. coli* and *K. pneumoniae* lines. Relative virulence was defined as host-killing rate (Cox hazard model estimate) in *G. mellonella* (as in panels B/C/D) or zebrafish larvae (panel E) relative to the corresponding ancestor. Zero indicates ancestral virulence; negative/positive values indicate decreased/increased virulence. Lines are grouped by virulence outcome, based on model estimates and statistical significance (likelihood-ratio test at 10% FDR correction, adjusted *P* < 0.1, see Methods). Strains in panels A-B: *E. coli* ATCC 25922; *K. pneumoniae* ATCC 10031; *K. pneumoniae* ATCC 700603. **(B)** Virulence outcome proportions across antibiotic adaptations. TOPO, topoisomerase inhibitors; TETR, tetracyclines; AMIN, aminoglycosides; CARB, carbapenems; MEMB, membrane-targeting antibiotics (Table 1). Continuous virulence differed between treatments (Kruskal-Wallis, *P* = 0.002823); categorical outcomes differed among classes (Monte-Carlo Fisher’s exact test, *P* = 0.02062; Cramér’s *V* = 0.21), reflecting excess increases among MEMB-adapted and decreases among AMIN-adapted lines. **(C–D)** Relative virulence for adapted lines (green), reconstituted BasR-G53V mutant (pink) and ancestors (black) of *K. pneumoniae* ATCC 10031 and *E. coli* ATCC 25922 along with clinical isolates (orange/blue). Clinical isolates represent common sequence-types identified through AMRWatch^131^ (Figure S5; Data S3). Hypervirulence was defined using Kleborate-defined^23^ virulence-factor genes or literature^27,90,91^ (Methods). Increased-virulence adapted lines did not differ significantly from hypervirulent isolates (*P* = 0.14 and 0.36). **(E)** Relative virulence of adapted lines (green) and clinical isolates (purple) of *K. pneumoniae* ATCC 10031 lines in zebrafish larvae (*n* = 30/group), grouped by Galleria-phenotype. Asterisks denote significant difference in virulence compared with the ancestor (Likelihood ratio test, ***/**/* indicates P ≤ 0.0001/0.01/0.05). Adapted lines with increased virulence exhibit virulence levels comparable to common hypervirulent clinical isolates (P = 0.13, Wilcoxon rank-sum test). Error bars in C-E indicate standard errors. **(F)** Epithelial cell (A549/Caco-2) survival after 3h infection with ancestor, BasR mutants or adapted lines of *K. pneumoniae* ATCC 10031 (≥6 biological replicates) was compared with the ancestor using cell-line-specific Gamma generalized linear models (log-link), two-sided Wald-tests and Benjamini-Hochberg correction: ****/* indicate adjusted P ≤ 0.0001/0.05. Green/red indicate increased/decreased virulence in *Galleria*.

Several adapted lines exhibited host-killing rates comparable to those of clinical isolates classified as hypervirulent based on established genetic markers (Fig. 2C-D, Figure S5, Data S3)^23,26,27^. Notably, neither the ancestral strain nor the adapted lines carry these canonical virulence markers (Figure S6), indicating that resistance-associated mutations alone can drive substantial increases in host-killing rate.

*K. pneumoniae* is uniquely suited for dissecting resistance-virulence interactions because it is one of the few pathogens in which multidrug resistance and hypervirulence increasingly co-occur in globally disseminated lineages^28,29^. Therefore, all subsequent analyses focus on deciphering the molecular mechanisms that drive increased virulence in *K. pneumoniae* and exploring their possible clinical consequences.

To confirm the robustness of the above findings, we next evaluated a representative subset of antibiotic-adapted *K. pneumoniae* strains using the zebrafish (*Danio rerio*) larval infection model^30,31^. Larvae were infected immediately after fertilization, and host-killing rates were determined daily for 72 hours after infection^32^. The bacterial strains exhibiting increased virulence in *G. mellonella* had significantly higher host-killing rates in zebrafish larvae compared to the ancestor (Fig. 2E, Data S2, see Methods), demonstrating that the increased virulence is reproducible across phylogenetically distinct hosts.

As epithelial surfaces represent the primary interface between *K. pneumoniae* and the host during infection, we next asked whether resistance-associated virulence phenotypes identified *in vivo* are reflected at the level of human epithelial cell damage. To address this, we quantified human epithelial cell survival following infection with the same panel of antibiotic-adapted *K. pneumoniae* lines. Lung (A549) and intestinal (Caco-2) epithelial cells were infected for 3 h with the antibiotic-sensitive ancestor or adapted lines, followed by quantification of surviving host cells. Reassuringly, strains with increased virulence as identified in *G. mellonella* and *D. rerio* consistently resulted in reduced epithelial cell survival in both A549 and Caco-2 cells (Fig 2F, Data S2). These effects were highly concordant across the two epithelial cell types, indicating that resistance-associated virulence alterations translate into reproducible epithelial cytotoxicity phenotypes across distinct cellular contexts (Figure S7, Data S2).

### Convergent up-regulation of virulence genes

Having identified antibiotic-adapted lines with increased virulence across multiple infection models, we next asked whether shared physiological mechanisms underpinned these phenotypic changes. We therefore compared transcriptomic profiles of the ancestral strain with 29 antibiotic-adapted lines under antibiotic-free conditions using RNA-sequencing (Figure S8, Data S4).

To identify transcriptional features associated with virulence, we compared expression profiles between lines with (n=11) and without (n=17) increased virulence, using a regression-based exploratory outlier framework (Methods). The selected increased-virulence lines arose across diverse antibiotic treatments, indicating that the observed transcriptional patterns are not restricted to a single mode of action or resistance pathway. This analysis identified 55 genes with disproportionately elevated expression in virulence-enhanced lines (Fig. S9). Predicted virulence factors were significantly enriched (Fisher’s test, OR ≈ 3.26, *P* = 1.69 × 10⁻⁴) in this set, indicating functionally relevant changes.

These genes clustered into a small number of recurrent functional modules (Figure S10). The clearest signal involved lipid A and LPS remodelling pathways, including the *arnBCADTEF* operon, *eptA*, and their regulators *basRS/pmrAB* and *phoPQ*^33,34^. A custom-curated set of LPS-related genes (Data S5) confirmed strong enrichment among the outlier genes (Fisher’s test, OR ≈ 48.8, *P* = 2.58 × 10⁻¹⁸). Additional enriched categories included cell wall and membrane biogenesis (COG category M: OR ≈ 5.3, FDR-adjusted *P* = 3.7 × 10⁻⁴) and inorganic ion transport and nutrient acquisition (COG category P: OR ≈ 5.0, FDR-adjusted *P* = 3.7 × 10⁻⁴)^35,36^. Of note, genes associated with fimbriae and capsule production showed no systematic upregulation in the evolved lines (Figure S11)^37^. Together, these data indicate that increased virulence is associated with recurrent transcriptional changes in lipid A/LPS remodelling and broader envelope-related pathways.

### Lipid A modifications link antibiotic resistance, cell surface characteristics, and virulence

To directly investigate how antibiotic adaptation reshapes lipid A structure, we performed high-performance mass spectrometry-based lipid A profiling on 35 adapted lines and the corresponding ancestor spanning diverse antibiotic selection regimes and virulence phenotypes. Across these lines, we detected 68 distinct lipid A species, defined by variation in phosphate headgroup modifications - most prominently 4-amino-4-deoxy-L-arabinose (L-Ara4N) and phosphoethanolamine (PEtn) - and by differences in acyl-chain number, length, saturation, and oxygenation (Figure S12).

Because most inter-line variation was driven by headgroup chemistry and acylation state, individual lipid A species were merged into feature-level groups capturing these properties (Fig. 3A, Data S2). Hierarchical clustering of changes relative to the wild type revealed three reproducible lipid A remodelling profiles strongly associated with virulence phenotypes (Fig. 3B, Figure S13).

**Figure 3.**
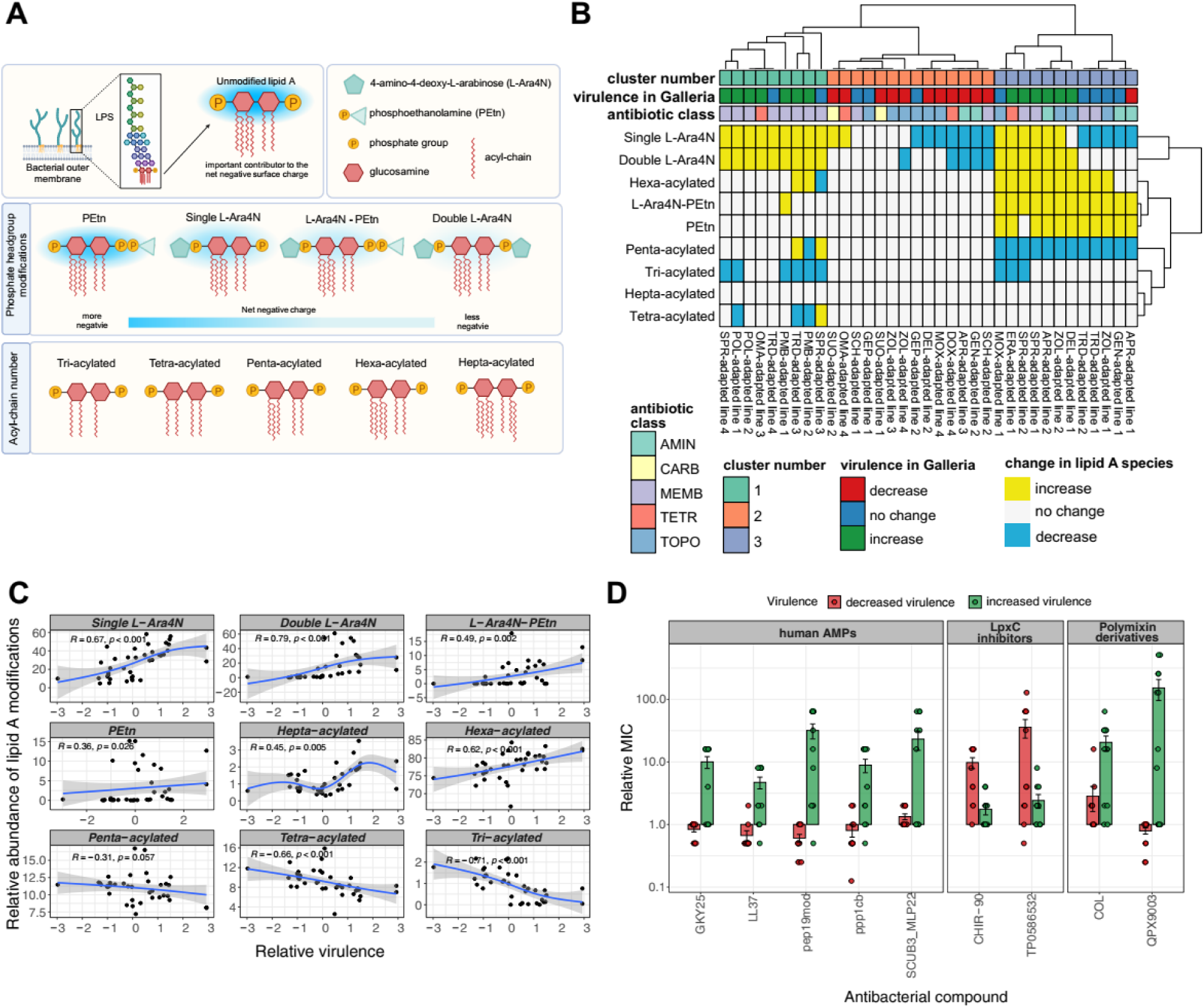
Association of lipid A structural modifications, virulence and cross-resistance in antibiotic-adapted *K. pneumoniae* lines. **(A)** Schematic figure of lipid A in the Gram-negative outer membrane, showing its glucosamine disaccharide backbone, phosphate groups and acyl chains. Phosphate-group modifications include phosphoethanolamine (PEtn), single or double 4-amino-4-deoxy-L-arabinose (L-Ara4N), and combined L-Ara4N–PEtn substitutions. The gradient indicates changes in net negative charge. Acylation states range from tri- to hepta-acylated forms. LPS, lipopolysaccharide. Created in BioRender®. **(B) Categorical heatmap of lipid A modifications across 35 antibiotic-adapted lines.** Columns represent lines and rows represent modifications. Colours indicate increased (yellow), decreased (blue) or unchanged (white) mean relative abundance compared with the ancestor, using thresholds of ±2 standard deviations around the ancestral mean. Relative abundance is expressed as a percentage of the total identified lipid A signal. Lines were hierarchically clustered using Gower distance and average linkage on the categorical change matrix. Annotations indicate cluster assignment, virulence phenotype in *G. mellonella* and selecting antibiotic class. (TOPO, topoisomerase inhibitors; TETR, tetracyclines; AMIN, aminoglycosides; CARB, carbapenems; MEMB, membrane-targeting antibiotics). **(C) Associations between lipid A modification abundance and relative virulence in *G. mellonella* across 35 adapted lines.** Each point represents one line, with modification abundance averaged across three biological replicates and expressed as a percentage of the total identified lipid A signal. Relative virulence is the Cox proportional hazards coefficient relative to the ancestor; zero indicates ancestral host-killing rate, while negative and positive values indicate decreased and increased virulence, respectively. Blue lines show generalized additive model fits, with shaded 95% confidence intervals. Associations were tested using Spearman’s rank correlation tests. **(D) Relative minimum inhibitory concentrations (MICs) of adapted lines against human-proteome- or oral- microbiome-derived antimicrobial peptides (AMPs), LpxC inhibitors and polymyxin derivatives.** Points represent individual lines; bars and error bars indicate means and standard errors, respectively. Red and green indicate decreased and increased virulence, respectively. MICs are normalized to the ancestor and plotted on a logarithmic scale. Cross-resistance denotes reduced susceptibility to antimicrobial compounds that were not used during experimental adaptation.

Cluster 1, characteristic of lines with increased virulence, was defined by a pronounced enrichment of single and double L-Ara4N modifications, a near absence of PEtn, a moderately elevated level of hexa-acylated lipid A species, and depletion of tetra- and tri-acylated variants. In contrast, Cluster 2, dominated by lines with decreased virulence, showed minimal L-Ara4N incorporation, little to no PEtn modification, and the lowest levels of total acylation, consistent with largely unmodified lipid A phosphates. Cluster 3 comprised lines with increased or unchanged virulence and was distinguished by elevated PEtn-containing lipid A species. Despite this distinct headgroup profile, lines showing increased virulence in Cluster 3 also showed high levels of L-Ara4N modification, suggesting that L-Ara4N rather than PEtn modification *per se* could be the dominant determinant of virulence (Figure S13).

Across all lines, increased virulence was consistently associated with enrichment of L-Ara4N modifications and elevated levels of total acylation relative to the wild type, whereas decreased virulence correlated with their depletion. These relationships were quantitatively supported by strong positive correlations between relative virulence and the abundance of single L-Ara4N, double L-Ara4N, and hexa-acylated lipid A species (Fig. 3C, Data S2). Notably, these lipid A remodelling patterns arose across adaptation to functionally distinct antibiotics. Consistent with these observations, the extent of L-Ara4N modification in the evolved lines correlated with transcriptional upregulation of the *arnBCADTEF* operon and its regulatory two-component system BasS/BasR, as determined by transcriptomic analysis (Figure S14).

Ordinal regression analysis controlling for antibiotic class and lipid A–associated mutations showed that the interaction between L-Ara4N modification and increased acylation accounted for nearly 85% of the observed variation in virulence (Note S4). Although PEtn modification has been implicated in virulence in some contexts^34,38,39^, our data indicate that its contribution is secondary to the dominant effects of L-Ara4N–driven charge modulation coupled with acylation.

Lipid A remodelling also linked resistance evolution to altered membrane biophysical properties. As expected, enrichment of L-Ara4N correlated with reduced outer-membrane surface charge (Figure S15, Data S2). Using DPH fluorescence anisotropy, we further showed that a representative virulence-enhanced mutant with enriched L-Ara4N modifications showed reduced membrane fluidity compared to the wild type at both 37 °C and 42 °C (Figure S16). Because membrane fluidity typically decreases with lowering temperature, we next quantified the magnitude of this effect. The fluidity shift observed in the mutant corresponded to an approximate temperature decrease of 6– 8 °C in model membranes (Figure S17), indicating a substantial alteration in membrane biophysical properties. These findings suggest that resistance-associated lipid A remodeling, and the resulting decrease in membrane fluidity, enhance membrane stability and host-killing potential.

### Association between virulence and cross-resistance to LPS-targeting antibiotics

As lipopolysaccharide (LPS) is a major target for antibiotic development^40,41^, we asked whether lipid A remodeling (and the associated shifts in virulence) also alter susceptibility to LPS-targeting compounds. Elevated virulence was strongly associated with resistance to colistin, a last-resort membrane-disrupting antibiotic, and to QPX9003 (F365), a synthetic lipopeptide in Phase I clinical trials designed to overcome colistin toxicity and resistance (Fig. 3D)^42^. Thus, selection for resistance to these compounds may inadvertently favor strains with altered virulence.

We next evaluated three recently discovered peptides: SCUB3_MLP22^15^, derived from the human proteome; PPP1CB^43^, from proteasomal cleavage of human proteins; and pep19mod^16^, originating from the proteomes of human oral bacteria (Fig. 3D, Data S2). All three peptides demonstrate robust antimicrobial activity in murine infection models and represent promising candidates for antibiotic development. Pep19mod was discovered through deep-learning–guided peptide design and optimization, SCUB3_MLP22 was identified through biophysical property-based mining of human proteins, while PPP1CB was discovered via *in silico* prediction of proteasome-derived defense peptides. Although these peptides share a membrane-targeting mode of action, their mechanisms are considered distinct from those of colistin and polymyxin B (Fig. 1, Note S1)^15,16,43–45^. Adapted lines with increased virulence showed markedly reduced susceptibility to these peptides, despite their distinct origins and discovery pipelines. Consistently, increased-virulence lines were also less susceptible to endogenous host defense peptides LL-37 and GKY25^44^, whereas lines with decreased virulence showed weak sensitization (Fig. 3D, Data S2).

We hypothesized that specific lipid A modifications may impose physiological constraints on cell envelope homeostasis, thereby creating vulnerabilities to compounds targeting lipid A biosynthesis. We examined the susceptibility of the antibiotic-adapted lines to the LpxC inhibitors CHIR-090 and TP0586532, which target an early, committed step in lipid A biosynthesis and exhibit potent activity against Gram-negative bacteria (Fig 1., Note S1)^44,45^. In contrast to membrane-disrupting and LPS-binding compounds, LpxC inhibitors remained effective against virulence-enhanced lines, with resistance observed in lines with reduced virulence only (Fig. 3D, Data S2). Notably, the extent of L-Ara4N modification correlated negatively with susceptibility to LpxC inhibitors, particularly with CHIR-090. In contrast, L-Ara4N levels correlated positively with resistance to membrane-disrupting compounds (Figure S18, Data S2). These opposing trends suggest that lipid A remodeling, while conferring resistance to several membrane-targeting compounds, may simultaneously increase dependence on intact lipid A biosynthetic flux.

In sum, distinct classes of LPS-targeting compounds differ fundamentally in their evolutionary consequences: membrane-disrupting compounds and host peptides may select for hypervirulent, lipid A–modified bacteria, whereas inhibition of the lipid A biosynthesis pathway may exploit these same adaptations. Future studies should study this possibility further.

### Pleiotropic side-effects of antibiotic resistance on host colonization and immune evasion

BasS/BasR, also referred to as PmrAB is an envelope-responsive two-component system that regulates lipid A remodelling through incorporation of L-Ara4N, reducing outer-membrane susceptibility to polymyxins^33,46–49^. Multiple BasR mutations arose independently in antibiotic-adapted lines, several of which are also found in natural isolates (Figure S1-2). We isolated three single BasR mutants (G53V, L105Q, and N109K), with all substitutions mapping to the N-terminal receiver domain and therefore potentially affecting response-regulator activation. G53V has previously been characterised as a gain-of-function substitution that drives constitutive expression of BasR/PmrA-regulated lipid A remodelling genes, whereas the specific signalling effects of L105Q and N109K remain to be established^50^. All three mutants displayed broad resistance to membrane-targeting antibiotics (Fig. S19), including host-derived antimicrobial peptides. In addition, they showed increased host-killing rate in *G. mellonella* (Fig. S19) and elevated epithelial cell toxicity (Fig. 2F).

We focused on BasR G53V, the most prevalent BasR variant detected in natural isolates, including isolates belonging to clinically important high-risk K. pneumoniae lineages with multiple independent phylogenetic origins (Figure S21). Transcriptomic profiling of BasR G53V relative to its isogenic K. pneumoniae ATCC 10031 ancestor revealed strong induction of lipid A metabolic pathways (Figure S22, Data S4), accompanied by increased L-Ara4N modification and acylation, closely recapitulating the lipid A profiles of laboratory-evolved lines with enhanced virulence (Fig. S23). Together, the transcriptional and lipid A profiling data indicate that BasR G53V increases activity of the BasRS/PmrAB-dependent lipid A remodelling program, consistent with an activating effect on this regulatory pathway.

Elevated host-killing rates of the G53V mutant may reflect either increased bacterial toxicity per se or an enhanced capacity for host colonization, which, in turn, drives host damage. Consistent with the latter scenario, the mutant rapidly outcompeted the ancestor in *G. mellonella* within 54 hours, despite a marked fitness disadvantage *in vitro* (Fig. 4A). This pattern indicates an evolutionary trade-off: although lipid A modification could incur a metabolic cost under laboratory conditions, it confers a strong selective advantage in the host environment. As might be expected, the mutation led to antibiotic treatment failure *in vivo* (Figure S24).

**Figure 4.**
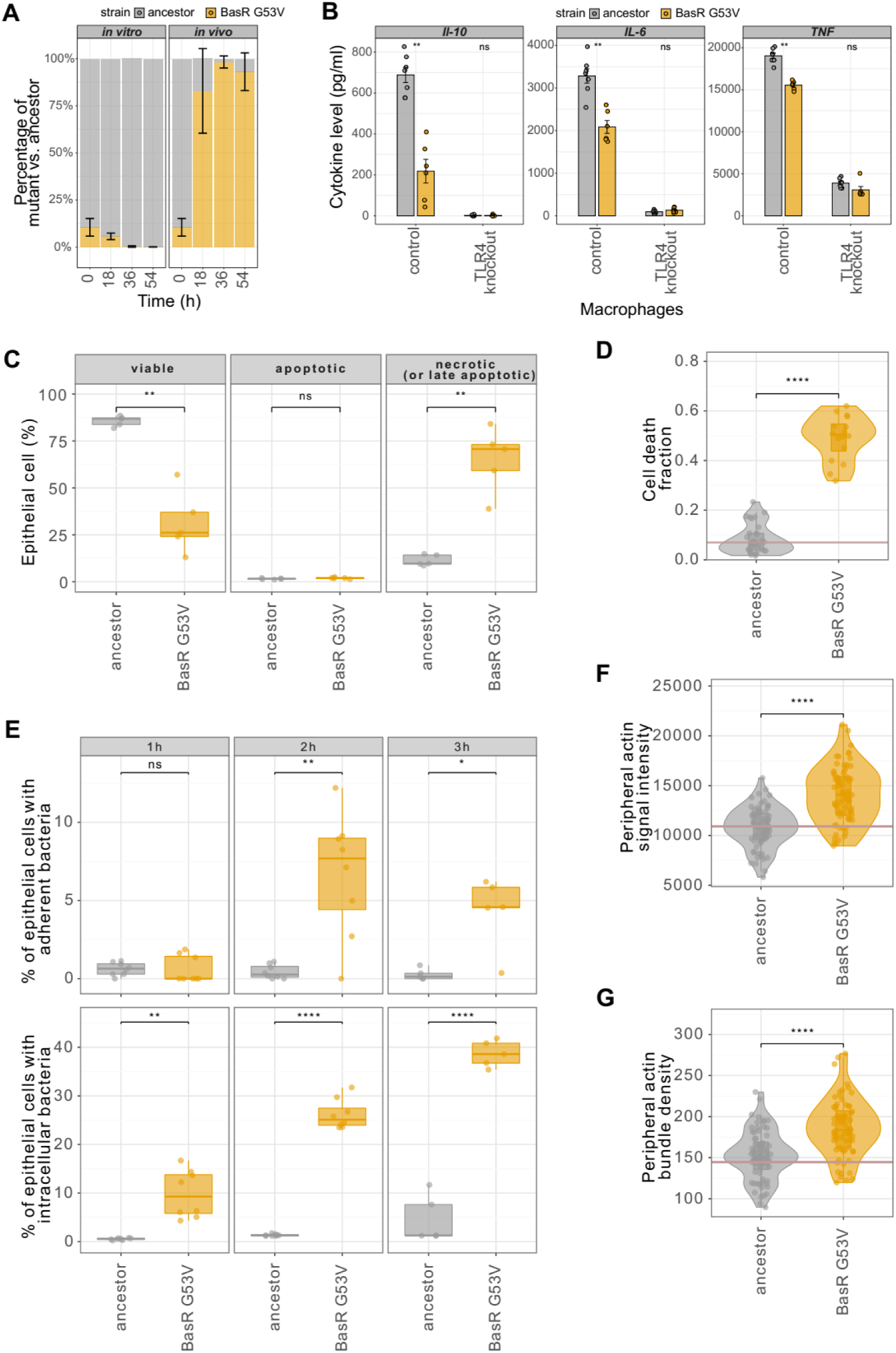
Competition and host-cell responses of the BasR G53V mutant *K. pneumoniae*. **(A)** *In vitro* and *in vivo* competition between *K. pneumoniae* BasR G53V and ancestor. Relative strain proportions (%) are shown across three growth cycles, from a 1:9 mutant-to-ancestor ratio. Six biological replicates; error bars indicate standard deviation. All panels use *K. pneumoniae* ATCC 10031 lines. **(B)** Cytokine production in control and TLR4-deficient macrophages following infection with the *K. pneumoniae* BasR G53V and ancestor. IL-6, IL-10, and TNF-α levels in immortalized bone marrow–derived macrophages (iBMDMs) and TLR4-knockout iBMDMs are normalized to ancestor control. Bars represent means ± SEM from six biological replicates pooled across two experiments. Wilcoxon rank-sum test: **, *P* < 0.01; ns, not significant. **(C)** Epithelial cell death following infection with the *K. pneumoniae* BasR G53V and ancestor. Proportions of viable (Caspase⁻/SYTOX⁻), apoptotic (Caspase⁺/SYTOX⁻), and necrotic or late-apoptotic (Caspase⁺/SYTOX⁺) A549 cells are shown after 1h infection. Three biological replicates. Two-sided Welch’s *t*-tests: **, *P* ≤ 0.01; ns, not significant. **(D)** Cytotoxicity in apical-out human-colon organoids. Violin plots show dead-cell-positive area relative to total organoid area after 1h infection with *K. pneumoniae* BasR G53V (*n*=16 organoids) or the ancestor (*n*=31). The horizontal line indicates the uninfected-control mean (*n*=38). Organoids originated from a single human biopsy. Two-sided Mann–Whitney *U*-test: ****, *P* < 0.0001. **(E)** Bacterial adhesion to and invasion of epithelial cells. Adhesion and invasion are quantified from percentages of YFP-positive A549 cells at 1h, 2h, and 3h following infection with *K. pneumoniae* BasR G53V and ancestor. Two-sided Welch’s *t*-tests at each time point: ****, *P* ≤ 0.0001; **, *P* ≤ 0.01; *, *P* ≤ 0.05. **(F–G)** Peripheral actin organization in epithelial cells following infection with the *K. pneumoniae* BasR G53V and ancestor. Violin plots show per-well medians of peripheral F-actin fluorescence intensity (F) and intensity-weighted filament density (G), reflecting filament length, thickness, and organization (Note S6). Each point represents one well; horizontal lines indicate uninfected-control medians. Linear mixed-effects models accounted for between-experiment variability, with Benjamini–Hochberg correction: ****, adjusted *P* < 0.0001. Boxplots in panels C-G show medians, interquartile ranges and 1.5× interquartile whiskers.

Modifications of lipid A could also influence the innate immune response. These modifications are expected to weaken lipid A recognition by the TLR4/MD-2 receptor complex on macrophages, dampening activation of the NF-κB pathway and the subsequent production of inflammatory cytokines^51^. Consistent with altered lipid A recognition due to L-Ara4N modifications, the G53V mutant elicited reduced TNF-α, IL-6, and IL-10 production from murine primary bone marrow-derived macrophages compared with the wild type (Figure S25). This difference was abolished in TLR4-deficient macrophages, demonstrating TLR4 dependence (Fig. 4B).

Together, these results show that a single resistance mutation can simultaneously remodel lipid A, confer broad antimicrobial resistance, enhance the host-killing rate, and dampen host immune response *in vitro*, highlighting how pathogenic traits can rapidly change as pleiotropic by-products of resistance evolution.

### BasR mutant increases epithelial association, invasion and cytotoxicity

Translocation across mucosal epithelia is a key step in *K. pneumoniae* pathogenesis, requiring coordinated adhesion, invasion, and manipulation of host cytoskeletal and signaling pathways^52,53^. The BasR G53V mutant elicited particularly strong epithelial cytotoxicity (Fig. 2E, Data S2). Flow cytometric analysis of infected A549 cells revealed a selective expansion of necrotic or late-apoptotic populations, defined by concurrent caspase-3/7 activation and SYTOX uptake, without a corresponding increase in early apoptosis (Fig. 4C, Data S2). This pattern indicates preferential induction of terminal epithelial damage^54^.

To assess whether this phenotype persists in a 3D *in vitro* infection model, we studied apical-out human colon organoids^55^. Infection with the BasR G53V mutant resulted in markedly elevated epithelial cell death relative to the ancestral strain, demonstrating that BasR-dependent cytotoxicity is preserved in an *ex vivo* model (Fig. 4D, Figure S26, Data S2).

Although *K. pneumoniae* is classically considered an extracellular pathogen, recent reports indicate strain-specific differences in its capacity to invade epithelial cells^52,56^. Therefore, we next quantified bacterial adhesion and invasion to identify determinants of epithelial damage. The BasR G53V mutant exhibited substantially increased epithelial association over time, reflecting both enhanced surface attachment and progressive intracellular accumulation (Fig. 4E, Data S2). Confocal microscopy of infected Caco-2 cells confirmed dense intracellular bacterial clusters following removal of extracellular bacteria, with tight-junction staining delineating epithelial boundaries and verifying intracellular localization (Fig. S27). These effects were independent of fluorescent reporter expression or antibiotic susceptibility, supporting increased epithelial association and intracellular accumulation by the BasR G53V mutant (Figure S28-29).

Ultrastructural analyses further linked the BasR G53V mutation to epithelial injury. Field-emission scanning electron microscopy of infected A549 and Caco-2 epithelial cells revealed extensive plasma membrane rupture, collapse of apical microvilli, and severe cortical surface disruption in cells infected with the mutant compared with the ancestor (Figure S30; Video S1).

Reorganization of cortical actin is a common strategy used by bacterial pathogens to facilitate attachment and trigger host membrane remodeling during entry^57^. Such actin dynamics generate membrane protrusions and localized cytoskeletal scaffolds that promote bacterial uptake and intracellular persistence^52,58,59^. Quantitative high-content imaging revealed pronounced reorganization of cortical actin in cells infected with the mutant. Radial zoning analysis showed a reproducible redistribution of F-actin from perinuclear regions toward the cell periphery, accompanied by increased peripheral actin intensity and elevated actin bundle density (Fig. 4F-G, Figure S31, Data S6, Note S6, Fig. SN2-3). This combination indicates enhanced membrane-proximal actin accumulation and stabilization of higher-order filament structures.

Together, these assays show that BasR G53V increases epithelial association, intracellular accumulation, cortical actin redistribution and terminal epithelial damage. These cellular phenotypes provide a direct explanation for the increased epithelial cytotoxicity observed in this resistance mutant.

### Rewired epithelial host–pathogen interactions alter infection dynamics

Finally, we tested whether the BasR G53V mutation alters infection dynamics in vivo using two murine infection models. An intravenous systemic infection model was used to assess bacterial persistence and dissemination across host tissues, whereas an intratracheal infection model was used to examine pulmonary infection.

In the systemic infection model, immunocompetent 6–8-week-old female BALB/c mice were infected by tail-vein injection with the ancestor or the BasR G53V mutant^52^. Bacterial clearance was defined as the net removal of viable bacteria from four organs (intestines, lungs, liver, and stomach) and was quantified over 72 hours. Based on CFU measurements, bacterial clearance estimates the net balance between two opposing forces: bacterial replication and host pathogen removal^60,61^. Across all four organs, the BasR G53V mutant exhibited a significant reduction in clearance relative to the ancestor, indicating substantially enhanced survival capacity in multiple organs, revealing a shift in pathogenic potential (Fig. 5A). Decreased bacterial clearance and increased colonization capacity are characteristic features of invasive liver abscess syndrome, a life-threatening condition in which hypervirulent K. pneumoniae spreads to multiple vital organs simultaneously^62^. To investigate this issue further, mouse livers were cryosectioned and processed for immunofluorescence microscopy. Consistent with cortical actin remodeling observed in epithelial cell models (Fig. 4F-G, Figure S31, Data S6), infection with the BasR G53V mutant was associated with substantial changes in host actin structure in the liver, suggesting altered interactions between the pathogen and liver cells during infection. As anticipated, intracellular accumulation of BasR G53V mutant bacteria within liver epithelial cells was observed after 24 hours (Fig. 5B, Video S2).

**Figure 5.**
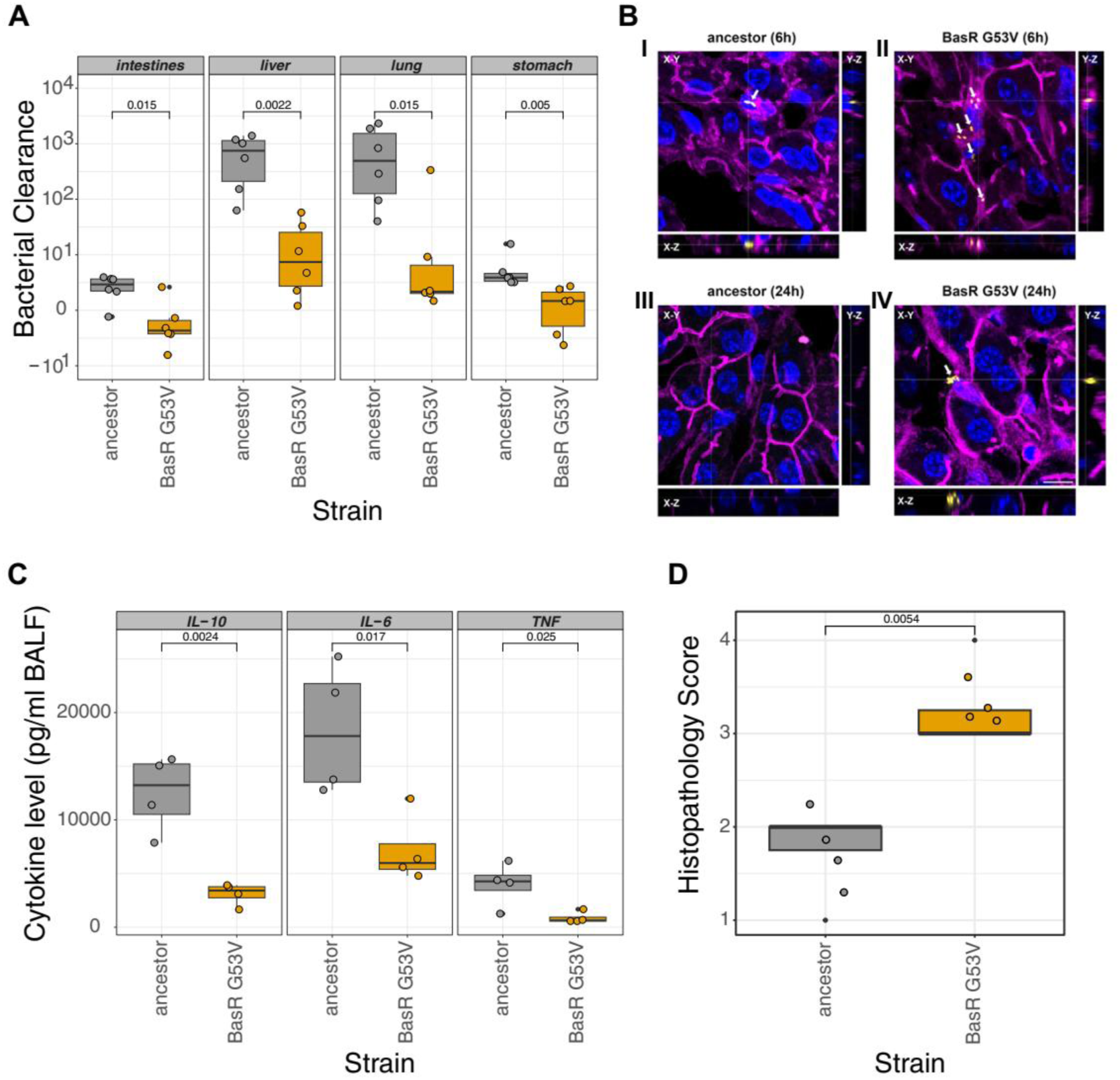
Systemic and pulmonary infection phenotypes of the BasR G53V mutant *K. pneumoniae* in immunocompetent female BALB/c mice aged 6–8 weeks. **(A)** Boxplots show clearance of *K. pneumoniae* ATCC 10031 BasR G53V and corresponding ancestor in the intestines, liver, lung, and stomach at 72h after intravenous infection. Clearance is expressed as Δlog₁₀(CFU) between the initial inoculum and 72h; negative values indicate net bacterial expansion. Each point represents the difference in bacterial burden between two animals sampled at 0h and 72h. Two-sided Wilcoxon rank-sum tests were used; exact *P* values are shown. **(B)** Hepatic persistence and cytoskeletal architecture following infection with the *K. pneumoniae* ATCC 10031 BasR G53V and ancestor *in vivo*. Representative confocal images show liver sections at 6h and 24h after intravenous infection with mScarlet-expressing ancestor (I, III) or BasR G53V mutant (II, IV). Maximum-intensity projections and corresponding X–Z and Y–Z orthogonal views are displayed. Bacteria (mScarlet), nuclei (Hoechst), and the actin cytoskeleton (CellMask counterstain) appear in yellow, blue, and magenta, respectively. Scale bar, 10 µm. Corresponding videos for the ancestor (Video S2A) and mutant (Video S2B) are available at [Zenodo](https://tinyurl.com/5xs5pywc) (Methods). **(C)** Boxplots show IL-10, IL-6, and TNF-α levels in bronchoalveolar lavage fluid collected 48 h after intratracheal infection with the ancestor or BasR G53V mutant *K. pneumoniae* ATCC 10031. Each point represents one animal; boxes indicate medians and interquartile ranges. Each group contained four animals. Strains were compared using Student’s *t*-tests; exact *P* values are shown. **(D)** Intracellular bacterial clustering during pulmonary infection with the *K. pneumoniae* ATCC 10031 BasR G53V mutant and ancestor. The boxplot shows bacterial localization assessed in hematoxylin–eosin-stained tissue at 48h after intratracheal infection (Methods). Observations were classified using an established semiquantitative scoring system^129^: 0, absent; 1, minimal (<10%); 2, mild (10–39%); 3, moderate (40–79%); and 4, marked (80–100%). Each group contained four animals. Strains were compared using Student’s *t*-tests; exact *P* values are shown. Boxplots in panels A, C-D show medians, interquartile ranges and 1.5× interquartile whiskers.

The pulmonary infection route was tested using a standard model (intratracheal injection) with immunocompetent 6-8 week old female BALB/c mice^63^. While inflammatory response and lung injury was detectable both in the case of the ancestor and the BasR G53V mutant, two important differences were identified. First, in line with previous *in vitro* results, the BasR G53V mutant elicited reduced TNF-α, IL-6, and IL-10 production, signifying a dampened host immune response (Fig. 5C). Additionally, histopathological analyses showed increased intracellular bacterial clustering in the BasR G53V mutant infected lungs, indicating that the mutant also displays enhanced epithelial invasion *in vivo* (Fig. 5D).

Overall, these findings indicate that mutational rewiring of host-pathogen interactions does not simply increase bacterial burden or tissue destruction *in vivo*. Instead, the BasR G53V mutation appears to alter the cellular niche occupied by the pathogen, promoting closer engagement with host cells and enhanced intracellular accumulation in multiple organs.

## Discussion

This study addresses a translationally important question for antimicrobial development: whether resistance to newly introduced or investigational antibiotics can create pathogens that are not only harder to treat but also more virulent. Resistance evolution did not uniformly increase virulence. Across the full dataset, resistance was associated with both increased and decreased host-killing capacity, indicating that pathogenicity can shift in either direction depending on the antibiotic class and the physiological route to resistance.

Across structurally and functionally diverse antibiotics, laboratory evolution repeatedly targeted an interconnected cell-envelope program, particularly lipid A remodelling (Fig. 6). Several adapted *K. pneumoniae* lines reached host-killing rates in *Galleria mellonella* comparable to those of clinical isolates carrying canonical hypervirulence-associated markers, despite lacking these markers themselves. This indicates that resistance-associated mutations alone can substantially increase host-killing capacity. The same classes of mutations were detectable in high-risk clinical isolates, and resistant bacteria showed increased persistence across host tissues in a systemic murine infection model, supporting their relevance to disease outcomes.

**Figure 6.**
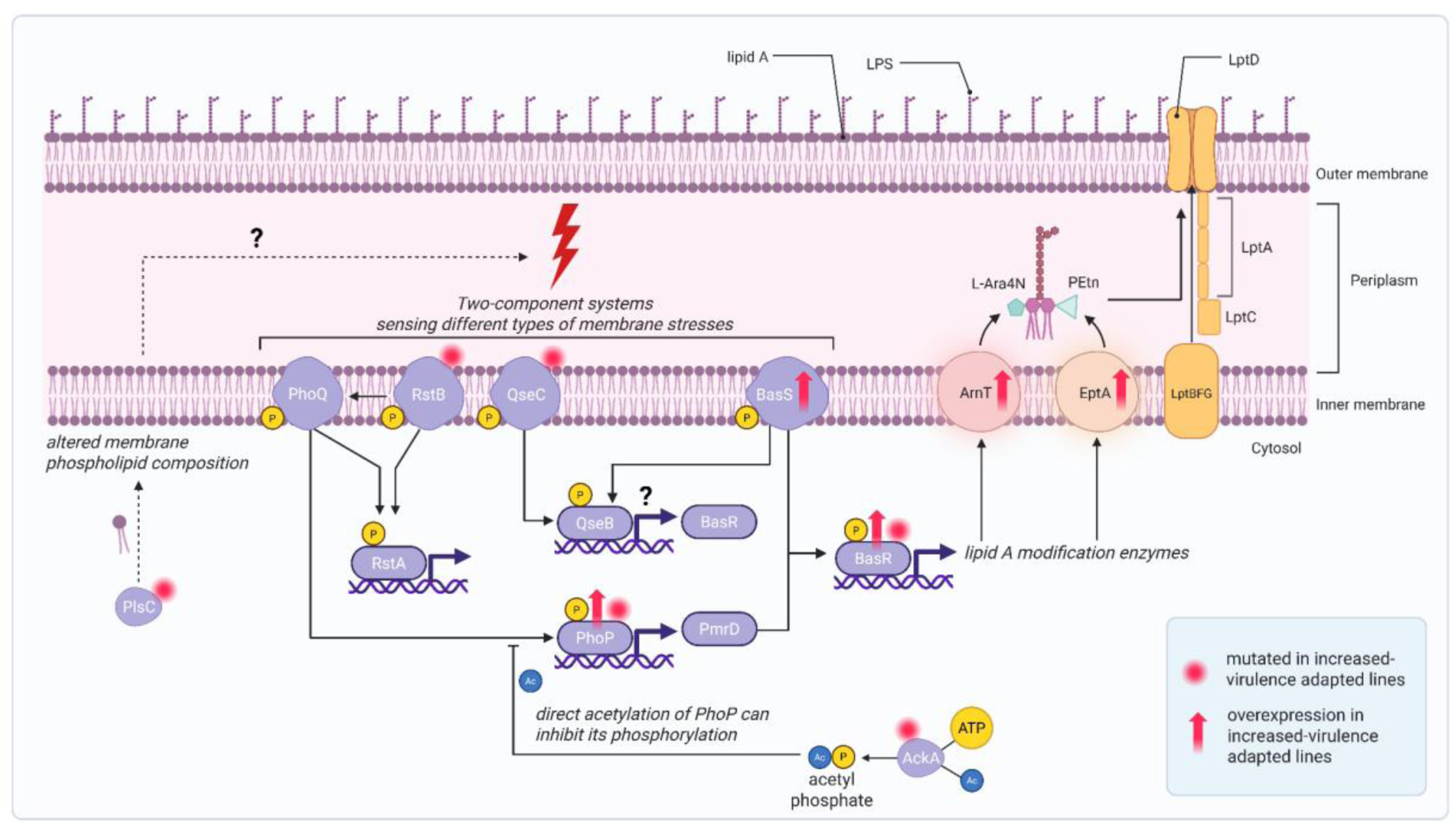
Interconnected regulatory routes converging on lipid A modification in virulence-increased *Klebsiella pneumoniae* adapted lines. Schematic model of the Gram-negative bacterial membrane-stress signaling network linking diverse resistance-associated mutations to increased lipid A remodeling. PhoPQ, RstAB, QseBC, and BasSR form an interconnected regulatory module that converges on BasR-dependent induction of *arnT* and *eptA*, promoting L-Ara4N and phosphoethanolamine (PEtn) modification of lipid A. PmrD links PhoP activity to BasR signaling, whereas altered phospholipid composition (PlsC) and acetyl-phosphate-dependent regulation (AckA) may further tune signal flow through this network. Red dots indicate proteins encoded by genes mutated in our adapted *K. pneumoniae* lines displaying increased virulence in multiple models; red arrows indicate genes with increased expression in increased-virulence lines relative to other adapted lines. The figure was created with BioRender^®^. (For a more detailed description, see Note S5.)

Mechanistically, our data identify cell-envelope regulatory rewiring as a central link between resistance and virulence. Distinct genetic routes to resistance converged on altered expression of pathways controlling lipid A modification, outer-membrane properties and host interaction. These changes were associated with epithelial invasion, resistance to host antimicrobial peptides and reduced macrophage inflammatory responses. A single resistance mutation was sufficient to transform *K. pneumoniae* from a weakly invasive epithelial colonizer into a highly adherent and cytotoxic pathogen, highlighting how single-step resistance events can have immediate effects on pathogenicity.

The implications are particularly important for membrane-active and peptide-based antimicrobials. *K. pneumoniae* is a major target for next-generation agents that disrupt or remodel the Gram-negative cell envelope, including host-derived and machine-learning-designed antimicrobial peptides^15,16,64^. Our results suggest that resistance to such compounds can select physiological states that couple reduced drug susceptibility with enhanced virulence and immune evasion. QPX9003/F365, a synthetic lipopeptide in clinical development, illustrates this risk: although designed to improve upon colistin-related limitations, resistance to this class of activity may favour lipid A-remodelled states associated with increased virulence^65,66^.

At the same time, not all LPS-directed strategies appear to carry equivalent evolutionary risks. Resistance-associated lipid A remodelling may create vulnerabilities to compounds that inhibit lipid A biosynthesis or transport rather than directly disrupting the outer membrane^41,45^. This distinction suggests that host–pathogen phenotyping could help prioritize antibiotic mechanisms less likely to select for virulence-enhancing resistance states.

Several limitations should be acknowledged. Mechanistic dissection focused primarily on *K. pneumoniae*, and the generality of these pathways in other pathogens remains to be established. Specifically, because detailed host–pathogen experiments were centered on one *K. pneumoniae* strain and a limited set of infection models, the extent to which lipid A remodelling, L-Ara4N modification and BasR-dependent virulence effects generalize across strain backgrounds, species and infection routes requires further analysis. Laboratory evolution enabled controlled testing of antibiotic selection but does not fully capture clinical pharmacokinetics, host immunity or microbiome-mediated selection. Our analysis also focused on chromosomal mutations and did not assess mobile resistance genes, whose contribution could be investigated using functional metagenomic approaches^20^. Although the identified mutations in BasR have been observed in high-risk clinical isolates and showed similar phenotypic effects across the experimental models examined, including murine, organoid and epithelial systems, their clinical relevance remains uncertain. Patient-level outcome data were not available, and whether these mutations are associated with disease severity, treatment failure or transmission in humans remains to be established.

In sum, by integrating laboratory resistance evolution, bacterial genomic surveillance and host– pathogen phenotyping, our work distinguishes resistance trajectories that merely reduce drug susceptibility from those that also alter virulence through clinically relevant mutations. A key value of our work is that it captures both directions of the resistance–virulence relationship: trajectories in which resistance is coupled to virulence attenuation, and trajectories in which resistance increases pathogenic potential. The results highlight compounds whose resistance pathways are coupled to epithelial invasion, immune evasion, systemic dissemination or treatment failure. Prospective validation across broader pathogen species, antibiotic classes and clinical isolate collections will be required, but applying similar approaches during preclinical development could help prioritize antimicrobial candidates that remain effective without inadvertently selecting for more invasive, immune-evasive or treatment-refractory pathogens.

## Materials and Methods

### Strains and growth conditions

Antibiotic-adapted lines analysed in this study were previously generated by frequency-of-resistance assays or adaptive laboratory evolution under antibiotic selection, as described in detail in our prior work ^20^. In addition, medium-adapted control lines were generated in the same adaptive laboratory evolution framework without antibiotic selection. *Escherichia coli* ATCC 25922, *Klebsiella pneumoniae* ATCC 10031, and *Klebsiella pneumoniae* ATCC 700603 were used as ancestors. ATCC 10031 is a laboratory/reference strain deposited as *K. pneumoniae subsp. pneumoniae* strain PCI 602 and commonly used in antibiotic assay, inhibition-testing, and quality-control contexts. ATCC 700603 was historically described as *K. pneumoniae* K6 and used as a multidrug-resistant reference strain^67^. Escherichia coli ATCC 25922 is a whole-genome-sequenced CLSI control strain widely used for antimicrobial susceptibility testing and quality control (ATCC 25922; CLSI AST Subcommittee, 2013). Whole-genome sequencing of adapted lines was performed in the original study. Detailed information on resistance levels, the antibiotic to which each antibiotic-adapted line was adapted, and the corresponding genomic alterations of all evolved lines is provided in Data S2 and described in detail in our prior work^20^.

Clinical isolates of *K. pneumoniae* and *E. coli* representing diverse sequence types and geographical origins are listed in Data S4. Clinical isolates were selected from a strain repository available in-house. The collection of clinical isolates complies with all relevant ethical regulations and was approved by the Scientific and Research Ethics Committee of the Hungarian Health Science Council (permit Reference Numbers BMEÜ/271-3/2022/EKU and BM/22994-1/2024). Three independent *K. pneumoniae* isolates, each carrying a single amino acid substitution in BasR (G53V, L105Q, or N109K), were derived from a previously described SPR-206–adapted population; whole-genome sequencing confirmed that no additional mutations were present.

Unless otherwise indicated, microbial cultivation was performed in cation-adjusted Mueller–Hinton Broth II. (MHB) (Merck, Millipore: 90922-500G) at 37 °C with shaking.

### Antibiotics

LL-37 and GKY25 were synthesized by ProteoGenix Company (custom peptide synthesis service). Colistin was purchased from Sigma-Aldrich. All other compounds (SCUB3_MLP22, PPP1CB-derived peptide, pep19mod, CHIR-090, TP0586532, and QPX9003) were synthesized by WuXi AppTec (custom synthesis service). According to the manufacturers’ HPLC analyses, the purity of all synthesized compounds was greater than 95%. Upon receipt, all compounds were reconstituted in sterile solvent according to solubility specifications. Following dissolution, solutions were sterile filtered using 0.22 µm syringe filters to remove potential particulate contaminants. Aliquots were prepared to avoid repeated freeze-thaw cycles and stored at −20 °C until use. Before each experiment, aliquots were thawed immediately prior to use and handled under sterile conditions. Additional information on the compounds used for cross-resistance measurements, including their targets and mechanisms of action, is provided in Supplementary Text 1. Figure 1 also summarizes these antibiotics together with their mechanisms of action.

### MIC measurement

A standard serial broth microdilution technique was used to determine MICs, as suggested by the Clinical and Laboratory Standards Institute guidelines^68^. A robotic liquid-handling system was used to automatically prepare 11-step serial dilutions in 384-well microtiter plates. A total of 5 × 10^5^ bacterial cells per ml were inoculated into each well containing 60 µl medium. Bacterial cultures were incubated at 37 °C with continuous shaking (300 rpm) for 18 h. All experiments were performed in two independent biological replicates, each measured in two technical replicates. Cell growth was monitored by measuring the optical density at 600 nm (OD600) values using a Biotek Synergy microplate reader. MIC was defined as the lowest antibiotic concentration at which OD_600_ remained below 0.05. Relative MIC was calculated as log_2_(MIC_evolved / MIC_ancestor).

### In vitro growth measurements

Growth phenotypes of strains with available whole-genome sequencing data were assessed in antibiotic-free cation-adjusted Mueller–Hinton Broth at 37 °C^69,70^. Approximately 5×10^5^ cells from early stationary-phase cultures were inoculated into 60 µL medium in 384-well microtiter plates. Optical density (OD600) was recorded every 5 min for ∼24 h using a BioTek Synergy HL1 microplate reader with continuous shaking. Each variant and its corresponding ancestor were measured in at least nine biological replicates. Relative fitness was approximated by the area under the growth curve (AUC), using established protocols^71^.

### Identification and classification of virulence-associated genes

To assess whether antibiotic adaptation targets genes linked to pathogenicity, we identified virulence-associated genes in *E. coli* and *K. pneumoniae* using VirulentHunter^21^. This deep-learning framework applies transformer-based protein language models derived from Evolutionary Scale Modeling 2 (ESM2)^72^. The base model (esm2_t30_150M_UR50D) was fine-tuned on curated virulence factor datasets compiled from the Virulence Factor Database (VFDB 2022)^73,74^, Victors^75^, and the Bacterial and Viral Bioinformatics Resource Centre (BV-BRC; formerly PATRIC)^76^. In contrast to homology-based approaches (e.g., BLAST against virulence databases), which depend on sequence similarity thresholds and can miss divergent determinants, protein language models learn statistical patterns from large protein corpora that correlate with structure and function^72,77^, enabling prediction beyond strict sequence conservation.

Complete reference genomes for *E. coli* ATCC 25922 and *K. pneumoniae* ATCC 10031 were obtained from NCBI RefSeq together with GFF3 annotations^78^. Protein-coding sequences were extracted by translating annotated CDS features; RefSeq protein FASTA files were used when available, otherwise CDS were translated in-frame from coordinates. The resulting proteomes contained 4852 predicted proteins for *E. coli* ATCC 25922 and 4872 for *K. pneumoniae* ATCC 10031.

Virulence prediction was performed on the full proteomes in batches of 250 proteins to improve stability and reproducibility. For each protein, VirulentHunter returned (i) a virulence probability score (vf_prob, 0–1) and (ii) multi-label predictions across virulence functional classes (aligned to VFDB categories), including adherence, invasion, biofilm formation, effector delivery systems, immune modulation, stress survival, exotoxin, exoenzyme, motility, nutritional/metabolic factor, antimicrobial activity/competitive advantage, post-translational modification, regulation, and others. Predictions were generated directly from the primary amino acid sequence, without incorporating genomic context or homology constraints.

Proteins with vf_prob ≥ 0.75 were classified as virulence associated. This cutoff was chosen to balance sensitivity and specificity based on model validation. Virulence-associated proteins were assigned to functional categories based on the highest-probability label; proteins exceeding thresholds for multiple categories were retained as multifunctional determinants. Category distributions were summarized genome-wide and compared between species. Functional enrichment analyses were performed relative to the full proteome background using Fisher’s exact test^79^, with Benjamini– Hochberg correction applied for multiple testing across functional categories^80^.

### Functional annotation of E. coli and K. pneumoniae proteomes

Predicted proteomes of *E. coli* and *K. pneumoniae* were analysed for functional annotation using the Clusters of Orthologous Groups (COG) database (2024 release)^81,82^. Protein sequences were searched against the COG reference protein set (COGorg24.faa) using DIAMOND (v2.1.x) in BLASTp mode^83,84^. A DIAMOND binary database was constructed with diamond makedb, and similarity searches were performed using tabular output format (outfmt 6). For each query protein, only the top-scoring hit was retained (--max-target-seqs 1).

DIAMOND outputs were processed in R (v4.4.3). Query sequence lengths were extracted from the corresponding FASTA files to calculate alignment coverage. Hits were filtered using the following thresholds: ≥90% amino acid identity, ≥90% query coverage, and E-value ≤1×10⁻⁵. For each protein, the highest bit score hit passing these criteria was retained as the representative COG assignment. Functional category information and COG definitions were obtained from the 2024 COG metadata tables, and proteins were assigned to functional categories according to the annotated COG entries.

### Enrichment analysis of functionally defined gene sets

To test whether antibiotic resistance evolution preferentially targets specific functional gene groups, we performed enrichment analyses for different gene sets: virulence-associated genes predicted by the VirulentHunter framework, functional categories defined by the Clusters of Orthologous Groups (COG) database^81^, and a manually curated set of genes involved in lipopolysaccharide (LPS) biosynthesis and modification (table S3).

Virulence-associated genes were identified using VirulentHunter^21^, a deep-learning–based prediction framework trained on curated virulence factor databases. Gene annotations for the *E. coli* and *K. pneumoniae* reference genomes were screened with VirulentHunter, resulting in 799 and 651 predicted virulence-associated genes, respectively. Functional annotations for broader cellular processes were obtained from the COG database, which assigns genes to conserved functional categories based on orthology. In addition, a manually curated set of genes involved in lipopolysaccharide (LPS) biosynthesis and modification was compiled based on genome annotations and previous studies of Gram-negative cell envelope structure. These annotations defined the genome-wide functional gene sets used in subsequent enrichment analyses.

Whole-genome sequencing data from experimentally evolved antibiotic-resistant populations were previously used to identify genes carrying mutations acquired during adaptation. For each evolved population, mutated loci were defined as genes containing at least one nonsynonymous mutation relative to the ancestor genome. To avoid counting multiple mutations within the same locus repeatedly, each gene was counted only once per evolved line. Because adaptive evolution often produces recurrent mutations in the same genes across independently evolved populations, we quantified recurrence-weighted mutational burden, defined as the number of evolved lines in which each gene was mutated. For each permutation, mutated loci were randomly reassigned while preserving the number of mutated genes per evolved line, and the total number of line-level mutation events falling within each gene set was recorded. Enrichment was calculated as the ratio between the observed recurrence-weighted mutational burden and the mean value obtained from the permutation distribution. Empirical p-values were calculated as the fraction of permutations producing values equal to or greater than the observed statistics.

To determine whether enrichment differed among antibiotic classes, recurrence-weighted mutation counts were calculated separately for each antibiotic class. Differences in enrichment across antibiotic classes were evaluated using a Poisson generalized linear model in which the observed mutational burden was modeled as a function of antibiotic class, with the expected mutational burden (derived from the permutation model) included as an offset. This approach tests whether enrichment ratios differ significantly among antibiotic classes while accounting for differences in mutation counts among evolved populations.

Multiple hypothesis testing across gene sets and antibiotic classes was controlled using the Benjamini–Hochberg false-discovery-rate correction. All statistical analyses were performed in R (version 4.4.3).

### Assessing the prevalence of laboratory-detected mutations in natural bacterial isolates

We assessed whether amino acid substitutions described in our previous study^20^ are present in natural populations of *E. coli* and *K. pneumoniae* using massive-scale population genomics as follows. We downloaded a publicly available comprehensive genomic dataset with consistent assembly and quality controls, maintained by the collaborative project “AllTheBacteria” (release: Augustus, 2024)^22^. Genome sequences and corresponding metadata are available in the Open Science Framework storage space^85^.

Gene prediction was performed using Prodigal v.2.6.3 ^86^, excluding ORFs <100 amino acids. Genomes were filtered for quality (≥95% completeness) using BUSCO v.5.4.6 with automatic lineage detection^87^. After exclusion of lines generated in our previous study, this yielded 393,816 *E. coli* and 72,080 *K. pneumoniae* high-quality genomes (table S1)^20^.

Next, we searched for the presence of each amino-acid-changing SNP across the *E. coli* and *K. pneumoniae* genome collections as follows. First, we performed a sequence similarity search of each gene carrying a given variant using DIAMOND BLAST (v.2.0.2)^83^ using an *e*-value (expect-value) of 10^-5^ with 90% coverage and 90% identity to identify orthologues among the genomes. Next, orthologs were aligned using MAFFT (v.7.475)^88^ with the –retree 2 option. The resulting multiple sequence alignments were parsed to determine the amino acid frequency at each mutated position using the custom R script from^89^.

To characterize the genomic and clinical context of genomes carrying specific amino acid variants, we leveraged precomputed genome annotations and metadata provided by the AllTheBacteria project^22^. To determine whether laboratory detected substitutions were preferentially associated with pathogenic strains, we classified isolates by isolation source, using metadata when available (table S1). Metadata was downloaded from the NCBI BioSample database (https://ftp.ncbi.nlm.nih.gov/biosample/) and standardized using an in-house script that applied a large language model, Mistral-small3.2:24b, run on the HUN-REN Cloud GenAI4Science service. Isolates recovered from blood, nervous system, reproductive, respiratory, urinary, wound, or other infection sites were classified as pathogenic, whereas the remainder were considered non-pathogenic. Additionally, multi-locus sequence types (STs), antimicrobial resistance genes, and virulence factors for *K. pneumoniae* isolates were obtained from Kleborate output files generated by the AllTheBacteria consortium and made publicly available via the Open Science Framework (https://osf.io/8r5kd/overview)^23^.

### Galleria bacterial infection model

*Galleria mellonella* 3rd instar larvae were obtained in bulk from local supplier company TerraPlaza (Vál, Hungary) and stored at 15°C without food. In sum, K. *pneumoniae* ATCC 110031, K. *pneumoniae* ATCC 700699, and *E. coli* ATCC 25922 ancestor strains, a set of antibiotic-resistant adapted lines, and a set of *K. pneumoniae* and *E. coli* clinical isolates were selected for the measurements. Among highly resistant lines within each ancestral-strain and antibiotic-treatment group, we selected lines with divergent fitness states spanning a range of *in vitro* growth-rate changes and carrying diverse adaptive mutations. Adapted lines were grouped by ancestral strain background and antibiotic treatment per evolution method utilized; lines with the highest resistance within each group were prioritized; where multiple highly resistant lines were available, lines with divergent growth phenotypes were selected; and, where growth phenotypes did not distinguish candidates, lines with distinct mutational profiles were chosen.

Clinical isolates were selected from a strain repository available in-house based on the following criteria: 1) they should represent sequence types (MLST) that are prevalent globally, and 2) multiple virulence factor genes should be present in them. Hypervirulent clinical strains were identified based on the presence of virulence factor genes, utilizing the scoring system defined in the Kleborate database or other literature sources^23,27,90,91^. Clinical isolates with a score of 2 or higher were considered hypervirulent clinical isolates. Prior to injection, all bacterial cultures were grown overnight in MHB broth and washed twice in 1x PBS. Bacterial suspensions were injected into the hemocoels of the larvae via the first left proleg using 10 μL Hamilton syringes (Reno, Nevada, U.S.A.). Larvae were incubated in petri dishes lined with filter paper (Roth, Rotilabo type 111A) at 37 °C for 48 h, and survival was documented every 6 hours. Insects were considered dead if they failed to respond to touch. The experiments were conducted on 30 animals per treatment group, with 10 animals per group exposed to a single biological replicate of each tested bacterial line. This sample size aligns with established protocols from previous studies using this animal model^17,92^.

To establish the size of the bacterial inoculum required to kill *G. mellonella* over 48h for the ancestor, 10 larvae were inoculated with 10 μL of bacterial suspensions in PBS containing 10^4^ to 10^8^ bacterial cells per animal. The number of viable cells in the bacterial suspension was estimated by counting the colony-forming units (CFU) on MHB agar. Based on this preliminary experiment, the 10^7^ inoculum size per animal was determined as ideal for killing *G. mellonella* larvae for *K. pneumoniae* ATCC 10031 and *E. coli* ATCC 25922, and 10^6^ for *K. pneumoniae* ATCC 700699.

All experimental data were visualized with Kaplan-Meier survival curves. All visualization and statistical analysis were performed with R version 4.2.2 and compatible packages survival, survminer, and ggsurvplot. Comparison of survival time between the evolved lines and the wild-type ancestor was evaluated using the Cox proportional-hazards model with the coxph() function built in the survival package (for further details, see Supplementary Note 3).

Antibiotic treatments were injected in the second right proleg using 10 μl Hamilton syringes (Reno, Nevada, U.S.A.). Antibiotic toxicity was determined by injecting 10–10 larvae with 5, 10, 20, 50, and 100 mg per body weight kg of SPR206, and no toxicity was observed at these dosages. SPR206 monotherapy was tested by first injecting the larvae with the appropriate *K. pneumoniae* ATCC 10031 strain or antibiotic-adapted line, then the SPR206 treatment dosage. Monotherapies were tested with 5, 10, 20, 50, and 100 mg per body weight kg doses. SPR206 at 50 mg per body weight kg improved survival by approximately 50% at the 48-hour mark, therefore this treatment dosage was chosen for the *in vivo* resistance experiment.

### Zebrafish bacterial infection model

Experiments were conducted in accordance with the Hungarian Animal Welfare Law (XIV-I-001/2303–4/2012) and Directive 2010/63/EU. All procedures were completed prior to the free-feeding stage. Adult AB strain zebrafish were maintained under standard laboratory conditions in a recirculating system at 27.5 ± 0.5 °C under a 14 h light/10 h dark cycle. Embryos were obtained by natural spawning.

Microinjection was conducted as described previously^32^. Bacterial cultivation was performed in Luria-Bertani (LB) broth (10.0 g tryptone, 5.0 g yeast extract, and 9.0 g NaCl in 1000 mL distilled water). Overnight liquid cultures were diluted with fresh LB to prepare a stock solution with an optical density of OD_600_ = 0.580 - 0.600. In the optimization phase, the stock solution was serially diluted to reach 10%, 2%, and 1% concentrations for injection. Based on these preliminary results, the final bacterial concentrations for the testing phase were set to 2% of the stock solution. Colony-forming units of the stock solutions were determined with the pour plate technique. Microinjection was performed in all cases immediately after fertilization. The freshly prepared bacterial suspensions were injected directly into the zebrafish egg yolk through the chorion in a volume of 1.77 nl (corresponding to a droplet with a diameter of 150 µm). The injection parameters were set as described by Csenki et al. The size of the injected droplets was checked every 25 embryos. As a control, sterile LB medium was injected for each experiment under the same conditions as in the case of the bacterial strains (injected control).

Two hours after injection, coagulated and/or non-fertilized eggs were discarded, and developing embryos were transferred in groups of 15 into 6 cm diameter Petri dishes filled with sterilized E3 medium (5 mM NaCl, 0.17 mM KCl, 0.4 mM CaCl_2_, and 0.16 mM MgSO_4_). The treatments were performed in two independent experiments, with five replicates per experiment. Microinjected embryos were incubated at 28 ± 0.5 °C in a Memmert Thermostat to ensure the optimal temperature for both test organisms. Embryonic mortality was monitored daily for 72 hours post-infection, defined by the following criteria: embryo coagulation, absence of tail detachment, and/or absence of somite formation, and/or absence of heartbeat.

Survival data were visualized using Kaplan–Meier curves. Differences between strains were assessed using a Cox proportional-hazards model implemented in R (v4.2.2).

### RNA extraction and purification

Bacterial cultures were grown in Mueller–Hinton broth II to mid-exponential phase (OD_600_ 0.28– 0.35). Cells were harvested by centrifugation at 4 °C, flash-frozen in liquid nitrogen and stored at −80 °C until extraction.Pellets were resuspended in TE buffer containing lysozyme (1 mg mL⁻¹) followed by Proteinase K and SDS treatment. Total RNA was extracted using TRIsure reagent according to the manufacturer’s instructions, including an additional chloroform extraction step. Genomic DNA was removed by DNase I treatment, and RNA was purified using NucleoSpin RNA Plant columns. RNA concentration and purity were assessed spectrophotometrically, and integrity was verified by agarose gel electrophoresis. Samples were subsequently used for RNA sequencing.

### RNA-seq sequencing

Total RNA samples were quality checked with the Tape Station 4200 instrument using Agilent RNA ScreenTape (Agilent Technologies USA, Cat. No. 5067-5576). Next-generation sequencing (NGS) library preparation was carried out on 1000ng RNA for each sample, using the NEBNext rRNA Depletion Kit (Bacteria) (NEB #E7850). Briefly, total RNA is hybridized with DNA probes targeting unwanted abundant RNAs (e.g., rRNA), followed by an RNase H digestion where the enzyme recognizes the RNA:DNA hybrid and degrades the targeted RNA. Finally, the DNA probes are digested with DNase I, and the reaction is cleaned using magnetic beads. Following depletion, the rRNA-depleted material, which contains biologically relevant cellular RNA such as non-coding RNA or mRNA, was used for NGS library preparation with the NEBNext Ultra™ II Directional RNA Library Prep Kit for Illumina (NEB #E7760). Briefly, cDNA was generated after fragmentation of depleted RNA, and second-strand synthesis was done with the dUTP method to retain strand specificity. After this, double-stranded cDNA was end-prepped, and Illumina-specific adaptors were ligated to the cDNA fragments, followed by a final enrichment PCR. Sequencing-ready libraries were quality control checked by the Tape Station 4200 instrument using D5000 ScreenTape (Agilent Technologies USA, Cat. No. 5067-5588). Sequencing was carried out with NextSeq 2000 P3 XLEAP-SBS Reagent Kit (300 Cycles) chemistry (Illumina, Inc., USA, Cat. No. IL20100988) targeting 5M reads/sample.

### RNA-seq analysis

RNA-seq reads were quality-filtered and adapter-trimmed using fastp v1.0.1^93^, and subsequently aligned to the *Klebsiella pneumoniae* ATCC 10031 reference genome using BWA-MEM v0.7.18^94^, which is appropriate for prokaryotic genomes lacking spliced transcripts. Gene-level read counts were generated from BAM alignment files using featureCounts^95^ with the corresponding genome annotation. Across samples, 82–91% of reads were assigned to annotated genes, indicating stable and reliable gene-level quantification; samples with slightly lower assignment rates remained within an acceptable range and were retained for downstream analysis.

Differential expression analysis was performed in R using edgeR ^96^. Genes with low expression were filtered using the filterByExpression function with default parameters, and library sizes were normalized using the trimmed mean of M-values (TMM) method. The dataset comprised six biological replicates per condition (3 independent experiments, including 2 replicates). Raw count matrices were modeled using negative binomial generalized linear models with a design matrix containing one coefficient per strain, and dispersion parameters were estimated using the empirical Bayes framework implemented in edgeR to account for biological variability. Differential expression between each evolved line and the ancestor strain was tested using quasi-likelihood F-tests and resulting *P*-values were adjusted for multiple testing using the Benjamini–Hochberg false discovery rate procedure. Genes were considered differentially expressed at FDR < 0.05 and |log₂ fold change| > 1.

The exploratory outlier analysis shown in Fig. S9 was performed separately from the edgeR differential-expression tests. For each adapted line, gene-level log2-fold changes were taken from the corresponding adapted-line-versus-ancestor comparison. Genes were summarized by the mean log2-fold change across increased-virulence lines and by the mean log2-fold change across lines without increased virulence. We then fitted a linear regression with the mean log2-fold change in lines without increased virulence as the response and the mean log2-fold change in increased-virulence lines as the predictor. For each gene, residuals from this regression were standardized across genes to obtain a residual z-score, and the difference between the two group means was likewise standardized across genes to obtain a between-group difference z-score. Genes were highlighted as exploratory outliers when they satisfied both criteria: |residual z-score| > 1.5 and |between-group difference z-score| > 2. Thus, replicate-level uncertainty was accounted for in the upstream edgeR estimates for each adapted-line-versus-ancestor contrast, whereas the Fig. S9 outlier rule was based on line-level summaries and did not directly model replicate uncertainty.

To determine whether specific functional gene groups are systematically associated with increased virulence, we applied a competitive permutation-based gene set analysis. First, each gene was assigned a regression coefficient (β) describing the direction and strength of the association between its transcriptional change relative to the ancestor (log_2_-fold change) and virulence across evolved strains. For each functional category, we calculated the difference between the mean β of genes within the category and the mean β of all other genes in the genome. Genes were grouped into predefined functional categories, including broad functional annotations (e.g., COG functional categories), virulence-related functional classes predicted by VirulentHunter, or manually defined gene sets such as lipid A/LPS remodeling genes and the arn operon (table S3). To assess statistical significance, category membership labels were randomly permuted across genes while keeping the regression coefficients fixed, and the difference in mean β values was recalculated for each permutation (10,000 iterations). This generated a null distribution representing the expectation if the functional category had no specific association with increased virulence. Empirical P-values were calculated as the fraction of permutations producing differences at least as extreme as the observed value and were corrected for multiple testing using the Benjamini–Hochberg false discovery rate (FDR) procedure. This approach identifies functional categories whose genes tend to undergo coordinated transcriptional shifts during adaptation that are associated with increased virulence while preserving the full quantitative information contained in the regression coefficients.

### Lipid A extraction

Lipid A was extracted from bacterial cultures using an established isobutyric acid-ammonium-hydroxide based extraction method^97,98^. Bacterial strains were inoculated from frozen stocks and into 3 mL of Mueller–Hinton broth II (MHB2) in three biological replicates. Cultures were incubated overnight at 37 °C with shaking at 250 rpm. A cell suspension volume corresponding to an OD₆₀₀ of 0.8 was centrifuged at 13000 rpm for 2 min in microcentrifuge tubes (Eppendorf Safe Lock Tube 1.5 mL). To remove most of the membrane phospholipids, bacterial cells were washed with 500 μl of a fresh, single-phase mixture of chloroform-methanol (1:2, v/v) and sonicated (Cole-Parmer 8851) for 10 minutes. Then, samples were centrifuged at 13000 rpm for 10 min. After discarding the supernatant, the pellets were dried under a flow chamber. Dried pellets were transferred to a screw-cap test tube (Sarstedt, 2mL SC Micro Tube PCR-PT) and suspended in 500 μl of concentrated isobutyric acid - 1M ammonium hydroxide (5:3, v/v) (isobutyric acid >99.5%, Sigma-Aldrich 58360-100ML; ammonia solution 32%, HiPerSolv CHROMANORM® for HPLC) and kept for 2 h at 100°C. Samples were centrifuged (13000 rpm for 10 min) and the supernatant was lyophilized in microcentrifuge tubes (Eppendorf Safe Lock Tube 1.5 mL).

Lipid A was recovered by two-phase liquid-liquid extraction. Lyophilized residues were resuspended in 1,190 μL of a one-phase mixture of chloroform:methanol:water (1:2:0.8, v/v/v) and vortexed thoroughly. Phase separation was induced by adding 940 μL chloroform and 220 μL 0.2 M KCl. The lower organic phase was collected, evaporated to dryness, and the lipid A residue was redissolved in 100 μL of chloroform:methanol:iso-propanol (1:2:1, v/v/v). Samples were stored at −20 °C until analysis.

### Mass spectrometry measurement of lipid A samples

Lipid A species were analysed by electrospray ionization mass spectrometry (ESI-MS) using an Orbitrap Fusion Lumos instrument (Thermo Fisher Scientific) equipped with a TriVersa NanoMate robotic nanoflow ion source (Advion BioSciences) and chips with spraying nozzles having a diameter of 5.5 μm. The ion source was controlled by Chipsoft 8.3.1 software. Measurements were performed in negative ion mode with an ionization voltage of −1.8 kV and a back-pressure of 1 psi. The temperature of the ion transfer capillary was 260 °C. Spectra were acquired over the m/z range 700– 2400 at the mass resolution of Rm/z 200 = 240,000 in full scan mode. All samples were analyzed in a single batch (within 2 days) following full calibration of the instrument, including the high mass range. For analysis, 6 μL of the lipid A extract was diluted with 30 μL chloroform:methanol:iso-propanol (1:2:1, v/v/v/); 10 μL of the diluted sample was directly infused, and data were acquired for 0.6 min. Distinct lipid A types – differing in the number of acyl chains in the hydrophobic region or having different modifications in the hydrophilic region, such as removal of phosphates or addition of phosphoethanolamine or amino-arabinose groups – were detected in negative ion mode.

### Data processing and annotation of lipid A samples

Based on published database searches ^99^, Lipid A identification was performed using in-house queries written for the LipidXplorer software^100^ by matching the monoisotopic m/z values to predefined elemental composition constraints with a mass tolerance of 3 ppm. The resulting data files were further processed by custom Excel macros.

Lipid A species were classified by phosphate-headgroup modification (A - single 4-amino-4-deoxy-L-arabinose; AA - double 4-amino-4-deoxy-L-arabinose; PEtn - phosphoethanolamine; APEtn - 4-amino-4-deoxy-L-arabinose and phosphoethanolamine; UM - unmodified). Species were annotated using sum formulas, e.g., AA(6:84:0:4:2), where the headgroup abbreviation is followed by parentheseis indicating the number of acyl chains, total chain length, total number of double bonds in the acyl chains, total number of oxygen atoms in the acyl chains, and the total number of phosphate groups. Lipid A intensities were obtained by averaging 50 MS1 scans and integrating signal after built-in C13 isotopic correction (both types I and II). Relative abundances were expressed as percentages of the total intensity from all identified lipid A species.

### Surface charge measurement

To investigate changes in bacterial cell surface charge, we performed a fluorescein isothiocyanate-labelled poly-L-lysine (FITC-PLL) (Sigma) binding assay^101^. In brief, cells were grown overnight in 8 biological replicates in Mueller–Hinton broth II (MHB2) medium and then washed twice with 1× phosphate-buffered saline (PBS) buffer. The cells were suspended in the PBS buffer to a final OD600 of 0.1. The suspension was incubated with 6.5 µg/mL FITC-PLL for 10 min and centrifuged at 5500 rpm for 5 min. Fluorescence of the supernatant (excitation 500 nm, emission 530 nm) was measured to quantify unbound FITC–PLL. Surface-bound FITC–PLL was calculated as the difference between total fluorescence in control samples (no bacteria) and fluorescence in bacteria-exposed samples.

### Membrane fluidity measurements

Bacterial bulk membrane fluidity was assessed by diphenylhexatriene (DPH) fluorescence anisotropy^102,103^. *K. pneumoniae* ATCC 10031 (ancestor) and the BasR G53V mutant strain were grown to early exponential phase (OD_600_ = 0.1) in four biological replicates. Cells were harvested, washed in PBS, and adjusted to OD_360_ = 0.1 in quartz cuvettes (Hellma 101-10-40 Suprasil® quartz UV/Vis cuvette (4 polished windows), 45 × 12.5 × 12.5 mm with 10 × 10 mm pathlength and 3500 µL volume). Samples were labelled with a final concentration of 0.2 µM DPH in 1.5 mL volume. After 10 minutes of incubation in the prewarmed sample holder, steady-state fluorescence anisotropy measurements were carried out at 37/42 °C using a Horiba Quantamaster T-format spectrofluorimeter. The excitation wavelength was set to 360 nm and the emission wavelength to 435 nm, with slit widths of 5 nm for both excitation and emission, and an integration time of 0.2 s. Fluorescence anisotropy (R) was calculated according to the following equation:

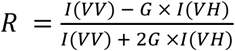

G is the correction factor accounting for the instrument sensitivity to vertically and horizontally polarized emission, VV denotes vertical excitation and emission, and VH denotes vertical excitation and horizontal emission. Membrane fluidity is reported as the inverse of fluorescence anisotropy (1/R).

### Large unilamellar vesicle preparation and temperature gradient membrane fluidity

Large unilamellar vesicles (LUVs) prepared from membrane lipid extracts of the *K. pneumoniae* ancestor (ATCC 10031) were used to assess membrane fluidity by DPH fluorescence anisotropy across a temperature gradient. This approach enabled estimation of the temperature shift corresponding to the fluidity differences observed between the ancestor and the BasR G53V mutant^104,105^.

300 mL culture of the ancestor *K. pneumoniae* was grown to early exponential phase (OD₆₀₀ = 0.1), harvested, and pelleted by centrifugation at 13,000 rpm for 2 min. Lipids were extracted with 9.5 mL of a single-phase water–methanol–chloroform mixture (1 : 2.5 : 1.25, v/v/v). Phase separation was induced by the addition of 3,750 μL chloroform and 880 μL of 0.2 M KCl. The lower organic phase was collected and evaporated to dryness, and lipid extracts (0.4mg) were resuspended by vortexing in 300 μL PBS containing DPH at a final concentration of 0.1 mM, followed by five freeze–thaw cycles (20 s in liquid nitrogen followed by 30 s at 40 °C) to improve lipid hydration. LUVs were formed by extrusion using a LiposoFast extruder (Avestin) equipped with two stacked 200 nm polycarbonate membranes (Avestin).

LUV suspension (15 μL, 1.8 mM) was diluted in 1.5 mL PBS in quartz cuvettes, and steady-state fluorescence anisotropy measurements were performed across a temperature range of 15–45 °C with 0.06 °C resolution using a Horiba Quantamaster T-format spectrofluorimeter, as described above.

### Macrophage assay

C57BL/6 mice were bred under specific-pathogen-free conditions. All animal procedures were performed in accordance with institutional and national ethical guidelines. Bone marrow-derived macrophages were differentiated as described previously^106^. Bone marrow cells from the femur, tibia, and humerus of 8–12-week-old male mice were differentiated for 6 days in DMEM supplemented with 15% FBS and 20% L929 conditioned medium. BMDMs were seeded at 5×10⁵ cells per well in 24-well plates 16 h prior to infection.

*K. pneumoniae* ATCC 10031 and the BasR G53V mutant were grown overnight in Mueller–Hinton broth II at 37 °C with shaking, washed, and resuspended in PBS. BMDMs were infected at a 1:50 macrophage-to-bacteria ratio in antibiotic-free medium for 6 h. Supernatants were collected, cleared by centrifugation, and stored at −20 °C until analysis.

Immortalized bone marrow-derived macrophages were generated as described previously^107^. TLR4 knockout cells were generated by CRISPR–Cas9 ribonucleoprotein electroporation using guide RNA targeting the Tlr4-coding genomic region. A non-targeting guide RNA served as a control. Edited cells were expanded and maintained in macrophage differentiation medium.

Immortalized bone marrow-derived macrophages (TLR4KO and control) were infected with *K. pneumoniae* ATCC 10031 or the BasR G53V mutant at a 1:50 macrophage-to-bacteria ratio for 6 h.

Secreted protein levels of TNF (DY410), IL-10 (DY417), and IL-6 (DY406) were determined using Mouse DuoSet ELISA Kits (DY008B; R&D Systems), in accordance with the instructions provided by the manufacturer. The plates were read using the Multiskan RC Thermo Microplate Reader.

### Epithelial cell lines and cell culture

Human colorectal adenocarcinoma Caco-2 cells (ATCC, #HTB-37) and human lung adenocarcinoma A549 cells (Sigma Aldrich, #86012804-1VL) were maintained under standard cell culture conditions. Caco-2 cells were cultured in DMEM High Glucose (4.5 g/l) with L-Glutamine (Capricorn Scientific, #DMEM-HA) supplemented with 10% (v/v) Fetal Bovine Serum (Capricorn Scientific, #FBS-HI-12A), 1% (v/v) MEM Non-Essential Amino Acids (Capricorn Scientific, #NEAA-B), and 1% (v/v) Penicillin-Streptomycin solution (Capricorn Scientific, #PS-B). A549 cells were cultured in a 1:1 mixture of DMEM/F-12 medium (Sigma Aldrich, # D8062-500ML), supplemented with 10% (v/v) Fetal Bovine Serum (Capricorn Scientific, #FBS-HI-12A) and 1% (v/v) Penicillin-Streptomycin solution (Capricorn Scientific, #PS-B). Cells were grown in 10-cm tissue culture dishes (SPL Life Sciences, #20101) at 37 °C in a humidified incubator with 5% CO₂. Mycoplasma contamination was routinely monitored using the MycoAlert™ Mycoplasma Detection Kit (Lonza, #LT07-518) according to the manufacturer’s instructions. Cells were passaged upon reaching approximately 90% confluence and were used for experiments for up to four weeks post-thaw.

### Cell plating prior to infection assays

Human cancer cell lines Caco-2 and A549 were used at passages 2–3 post-thaw and cultured as adherent monolayers to 70–85% confluence in 10-cm tissue culture dishes. Prior to seeding, cells were washed twice with 1× phosphate-buffered saline (PBS; prepared from a 10× stock; Capricorn Scientific, #PBS-10XA) and detached using trypsin–EDTA solution (Capricorn Scientific, #TRY-1B) for 5 min at 37 °C. Enzymatic dissociation was quenched by the addition of complete culture medium, and cells were collected by gentle resuspension.

Cell viability was assessed by mixing the cell suspension 1:1 with Trypan Blue solution (Gibco, #15250-061), and viable cells were quantified using a LUNA-II™ automated cell counter (Logos Biosystems). Based on viable cell counts, defined numbers of cells (as specified for each assay) were seeded into tissue culture–treated plates and incubated for 24 h at 37 °C in a humidified atmosphere containing 5% CO₂ prior to infection experiments.

### Bacterial infection of human cells

Bacterial strains were stored at −80 °C in dimethyl sulfoxide (DMSO)–containing stocks until use. For infection experiments, fluorescently labelled (yellow fluorescent protein, YFP-expressing) or unlabelled *K. pneumoniae* strains were cultured in Mueller–Hinton Broth II (MHB-2) for 16 h at 37 °C with shaking at 200 rpm in sterile polypropylene tubes. Following overnight growth, bacterial cultures were equilibrated to room temperature, and 100 µl aliquots were transferred to transparent 96-well plates for optical density measurements. Optical density was measured using a Synergy HTX multimode microplate reader (BioTek). Based on the measured values, bacterial cultures were diluted to achieve a final OD₆₀₀ of 0.25 or 0.5, as specified for each experiment. The required volume of bacterial suspension was transferred to 15-ml conical tubes containing 1× phosphate-buffered saline (PBS) to remove residual growth medium and centrifuged at 4,500 rpm for 5 min. The supernatant was discarded, and bacterial pellets were resuspended in antibiotic-free human cell culture medium supplemented with 10% (v/v) FBS and 10 mM HEPES (Capricorn Scientific, #HEP-B).

For infection assays, culture medium was aspirated from human cells that had been seeded 24 h earlier, and infection medium containing bacteria was added. Non-treated control cells received infection medium without bacteria. Cells were incubated for 1–3 h at 37 °C in a humidified atmosphere containing 5% CO₂. Following co-incubation, cells were washed with 1× PBS, and downstream assays were performed as described in the corresponding sections.

### Immunofluorescence staining and visualization of ZO-1

To visualize zonula occludens-1 (ZO-1), Caco-2 cells were seeded at a density of 6.0 × 10⁴ cells per well onto glass coverslips (5 mm diameter) placed in 24-well tissue culture plates. Cells were allowed to adhere overnight and subsequently infected with YFP-labeled ancestor or YFP-labeled BasR G53V mutant strains (final OD₆₀₀ = 0.5) for 2 h at 37 °C in a humidified incubator with 5% CO₂ for 1h or 3h. After infection, cells were treated with 100 μg/ml gentamicin-containing cell culture medium to remove the extracellular bacterial cells. Samples were washed with 1× phosphate-buffered saline (PBS) to remove non-adherent bacteria and treated with gentamicin (100 µg ml⁻¹) for 1 h at 37 °C to eliminate extracellular bacteria. After antibiotic treatment, cells were washed with 1× PBS and fixed with 4% (w/v) paraformaldehyde (PFA) for 10 min at room temperature. Cells were then permeabilized with 0.5% (v/v) Triton X-100 (Sigma Aldrich, #T8787-100ML) in PBS for 10 min and blocked for 1 h at room temperature in blocking buffer consisting of 3% bovine serum albumin (BSA) and 0.1% (v/v) Triton X-100 in PBS. Samples were incubated overnight at 4 °C with primary antibodies diluted in blocking buffer (rabbit anti-ZO-1; Thermo Fischer Scientific, #61-7300). After three washes with 0.1% (v/v) Triton X-100 in PBS, cells were incubated for 1 h at room temperature in the dark with fluorescently conjugated secondary antibodies (anti-rabbit Alexa Fluor 555; Thermo Fischer Scientific, # A-21428) diluted in blocking buffer. Cells were subsequently washed twice with 0.1% (v/v) Triton X-100 in PBS and counterstained with Hoechst (1 µg ml⁻¹ in PBS) for 15 min.

Z-stack images were acquired by confocal fluorescence imaging using an Opera Phenix™ Plus High-Content Screening System equipped with a 40× water-immersion objective (NA 1.1; Part no. HH14000422). Images were collected with a camera region of interest of 2,160 × 2,160 pixels using 2× binning. Fluorophores were excited with diode lasers at 405, 488, and 561 nm. Maximum-intensity projections were generated from z-stacks, and images were processed using Fiji (ImageJ)^108^.

### Cytotoxicity assay

Caco-2 and A549 cells were seeded into 24-well tissue culture plates at densities of 7.0 × 10⁴ and 6.0 × 10⁴ cells per well, respectively. Cells were allowed to adhere overnight prior to infection. Bacterial strains were prepared as described above and diluted in antibiotic-free human cell culture medium supplemented with 10 mM HEPES to achieve a final OD₆₀₀ of 0.5 for Caco-2 cells and 0.25 for A549 cells. Cells were infected for 3 h at 37 °C in a humidified incubator with 5% CO₂. Following infection, the inoculum was aspirated and replaced with complete culture medium containing primocin (100 µg ml⁻¹) (InvivoGen, #ant-pm-1) to eliminate extracellular and intracellular bacteria. Cells were incubated for a further 48 h at 37 °C with 5% CO₂, with a complete medium exchange performed after 24 h.

Cell viability was assessed by Trypan Blue exclusion. Cells were harvested, stained with Trypan Blue at a 1:1 ratio, and viable cells were quantified using a LUNA-II™ automated cell counter with the following settings: dilution factor = 2, noise reduction = 5, live cell sensitivity = 7, roundness = 85%, minimum cell size = 3 µm, maximum cell size = 60 µm, and declustering level = low. Viable cell numbers were determined for each condition and normalized to uninfected control samples, which were defined as 100% survival. Survival fractions of infected samples were calculated relative to this control.

Statistical analyses were performed by comparing each bacterial strain to its corresponding ancestor using generalized linear models with a Gamma distribution and log link, fitted separately for each cell line. Two-sided Wald test p-values were adjusted for multiple testing using the Benjamini– Hochberg false discovery rate (FDR) correction. FDR-adjusted significance levels are indicated in the figures as follows: **** p ≤ 0.0001, *** p ≤ 0.001, ** p ≤ 0.01, * p ≤ 0.05.

### Antibiotic susceptibility assays

Antibiotic susceptibility to primocin and gentamicin was assessed under host cell culture–mimicking conditions. Bacterial strains were cultured overnight in Mueller–Hinton Broth II (MHB II). Cultures were pelleted by centrifugation, washed once with 1×PBS, and adjusted to an OD₆₀₀ of 0.5 in either DMEM/Ham’s F-12 or DMEM high-glucose cell culture medium. Suspensions were transferred to 96-well plates and incubated for 3 h at 37 °C in a humidified atmosphere containing 5% CO₂. For primocin susceptibility testing, following the initial incubation, media were aspirated, and bacterial samples were divided into antibiotic-treated and untreated conditions by adding medium supplemented with primocin (100 µg ml⁻¹) or antibiotic-free control medium. For gentamicin susceptibility assays, *K. pneumoniae* ATCC 10031 ancestor and BasR G53V mutant were processed in parallel using medium containing gentamicin (100 µg ml⁻¹) or antibiotic-free medium. Following a 1 h incubation, the antibiotic-containing medium was aspirated and replaced with fresh, antibiotic-free medium.

After 24 h of incubation, bacterial suspensions were diluted 1:10,000, and 10 µl aliquots were spotted onto MHB II phyto agar plates using a Platemaster spotting device (Greiner). Spots were allowed to air-dry prior to incubation, and colony-forming units within each spot were enumerated to assess bacterial survival. To validate cytotoxicity assay conditions, all KP-SEN strains were tested for primocin susceptibility. Complete sensitivity to primocin confirmed efficient elimination of bacteria following infection, allowing epithelial cell recovery and accurate quantification of surviving host cells 48 h post-infection.

### Adhesion and invasion assay

YFP-labelled *K. pneumoniae* ancestor or BasR G53V mutant strains were cultured as described above. The assay was performed using established protocols^109–111^. A549 lung epithelial cells were seeded into 24-well tissue culture plates at a density of 6.0 × 10⁴ cells per well and allowed to adhere overnight. On the following day, cells were labelled with Hoechst 33342 (Invitrogen, #H3570) (1:10,000 dilution in complete human cell culture medium) for 1 h at 37 °C in a humidified incubator with 5% CO₂. Cells were subsequently infected with YFP-labelled *K. pneumoniae* strains at a final OD₆₀₀ of 0.5 for 1, 2, or 3 h. For each condition, parallel samples were processed to quantify total cell-associated bacteria (adhesion + invasion) or intracellular bacteria (invasion only). To assess total cell-associated bacteria, infected cells were washed with 1xPBS to remove non-adherent bacteria and directly analyzed. To quantify invaded (intracellular) bacteria, extracellular bacteria were eliminated by incubating cells with gentamicin (100 µg ml⁻¹) for 1 h at 37 °C. Following antibiotic treatment, cells were washed with 1× PBS prior to analysis.

Samples were analyzed by flow cytometry using an Attune™ NxT flow cytometer (Thermo Fischer Scientific), and data were processed with Invitrogen Attune Cytometric Software. At least 10,000 viable epithelial cell events per sample were acquired and gated based on Hoechst fluorescence. Bacterial association was quantified by YFP fluorescence. Invasion was defined as the fraction of Hoechst-positive epithelial cells containing gentamicin-resistant (intracellular) YFP-positive bacteria. Adhesion was calculated as the difference between total cell-associated bacteria (adhesion + invasion) and the invaded fraction, as described previously ^112–114^. Adhesion and invasion fractions were plotted for each condition. All experiments were performed in at least three independent biological replicates. Statistical comparisons between the ancestor and the BasR G53V mutant strain were conducted separately for each time point and readout using two-sided Welch’s t-tests.

### Necrosis/Apoptosis assay

Bacterial strains were cultured as described above. A549 human lung epithelial cells were seeded into 6-well tissue culture plates at a density of 2.0 × 10⁵ cells per well and allowed to adhere overnight. Prior to infection, epithelial nuclei were pulse-labelled with Hoechst 33342 to enable identification of host cells during flow-cytometric analysis. Cells were subsequently infected with either the Ancestor strain or the BasR G53V mutant at a final OD₆₀₀ of 0.5 in antibiotic-free DMEM supplemented with 10% fetal bovine serum (FBS) and 100 µM HEPES. Infections were carried out for 1 h at 37 °C in a humidified incubator with 5% CO₂. Following infection, host-cell death pathways were assessed by flow cytometry using the CellEvent™ Caspase-3/7 Green Flow Cytometry Assay Kit (Invitrogen, #C10740), according to the manufacturer’s instructions. Cells were incubated with CellEvent™ Caspase-3/7 Green Detection Reagent for 45 min to detect activated caspase-3/7, followed by incubation with SYTOX™ AADvanced™ Dead Cell Stain for 5 min to assess membrane integrity. Hoechst 33342 labelling was used to gate epithelial cells and exclude non-cellular events^115^. Samples were analyzed by flow cytometry using an Attune™ NxT flow cytometer (Thermo Fischer Scientific), and data were processed with Invitrogen Attune Cytometric Software. The analysis resolved three epithelial cell populations: viable cells (caspase⁻/SYTOX⁻), apoptotic cells (caspase⁺/SYTOX⁻), and necrotic or late-apoptotic cells (caspase⁺/SYTOX⁺)^116^. At least 30,000 Hoechst-positive epithelial cell events were acquired per sample. The proportions of cells within each population were quantified and compared between infections with the ancestor and the BasR G53V mutant strains. Statistical significance for differences in each cell population was assessed using two-sided Welch’s t-tests.

### Cytoskeleton rearrangement and F-actin staining

Bacterial strains were cultured as described above. The assay was performed using a protocol adapted from a previously described method^117^. A549 human lung epithelial cells were seeded at a density of 6.0 × 10³ viable cells per well into tissue culture–treated 96-well plates (Greiner Bio-One, #655090) and allowed to adhere overnight prior to infection. Cells were infected with bacterial cultures diluted to a final OD₆₀₀ of 0.25 and incubated for 1 h at 37 °C in a humidified atmosphere containing 5% CO₂. Following infection, cells were fixed by the addition of 4% paraformaldehyde (PFA; Thermo Scientific Chemicals, #J19943.K2) in 1×PBS and incubated for 10 min at room temperature to preserve cellular architecture. Cells were then permeabilized with 0.5% (v/v) Triton X-100 in 1× PBS for 10 min at room temperature, followed by a wash with 0.1% (v/v) Triton X-100 in PBS to reduce residual detergent. F-actin structures were stained using CellMask™ Deep Red Actin Tracking Stain (Invitrogen, #A57245) diluted 1:1,000 in 0.1% (v/v) Triton X-100 in PBS and incubated for 1 h at room temperature. Cells were subsequently washed with 0.1% (v/v) Triton X-100 in PBS and counterstained with Hoechst 33342 (Invitrogen, #H3570) diluted 1:10,000 in PBS for 1 h to visualize nuclei. After a final wash with 1× PBS, 100 µl of PBS was added to each well, and plates were stored at 4 °C until imaging.

Z-stack images were acquired by confocal fluorescence imaging using an Opera Phenix™ Plus High-Content Screening System equipped with a 40× water-immersion objective (NA 1.1; Part no. HH14000422). Images were collected with a camera region of interest of 2,160 × 2,160 pixels using 2× binning. Fluorophores were excited with diode lasers at 405 and 561 nm.

### Quantitative analysis of cytoskeletal rearrangement

Cytoskeletal rearrangement experiments were performed in three independent biological replicates, with a minimum of 2,000 cells quantified per sample. Image analysis was carried out using a custom Python-based workflow developed based on the framework described in^118^.

Nuclear and whole-cell segmentation were performed using a customized Cellpose deep-learning model, enabling robust boundary detection across heterogeneous epithelial cell morphologies^119^. Actin channel images were processed following established cytoskeletal analysis pipelines. Images were first deconvolved and subjected to Gaussian filtering to suppress imaging artefacts and diffuse cytoplasmic signal. Polymerized actin bundles were enhanced using a multiscale Hessian-based Sato tubeness filter^120^, which selectively emphasizes curvilinear filament structures. The enhanced signal was reduced to a one-pixel-wide topological skeleton using a ridge-detection algorithm, preserving filament continuity and network connectivity.

Shape-independent spatial analysis was performed using a nuclear-anchored radial zoning strategy adapted from previously described methods^121^. For each cell, the cytoplasm was partitioned into ten concentric radial bins normalized from the nuclear boundary (r = 0) to the cell periphery (r = 1). The peripheral region was defined as the outer one-third of bins, and the perinuclear region as the inner one-third. Radial mean actin bundle density was computed as the mean filament skeleton signal within each radial bin, normalized to bin area, providing a spatially resolved measure of structurally prominent filament organization along the radial axis. Peripheral actin organization was summarized as the ratio of peripheral to perinuclear (closest to the nuclei) radial mean actin bundle density, quantifying the degree of radial redistribution toward the cell edge.

Image-derived cell-level measurements were collected across three independent biological experiments, with experiment identifiers retained throughout processing to account for batch structure. To respect the nested data structure (cells measured within wells), preprocessing was performed at the cell level, whereas statistical inference was conducted at the well level. Cells were filtered using a quality threshold based on skeleton fidelity (≥ 0.25). To limit the influence of extreme values without distorting treatment-specific effects, outliers were removed at the cell level within Treatment × Well groups using an interquartile range–based method (Tukey fences; ±1.5× IQR). Across all experiments, a minimum of 2,500 BasR G53V mutant cells and 2,800 ancestor cells per experiment were retained after filtering. Filtered cell-level feature values were aggregated to well-level medians, yielding one value per feature per well per treatment. Treatment effects were assessed using linear mixed-effects models comparing conditions while accounting for experiment-to-experiment variability. Multiple testing across features was controlled using the Benjamini– Hochberg false discovery rate (FDR) correction. Adjusted p values were used for statistical inference and graphical annotation (*p < 0.05, **p < 0.01, ***p < 0.001, ****p < 0.0001; ns, not significant). Well-level distributions were visualized using violin plots of per-well median values. For reference, the median value of the uninfected control condition was included as a horizontal line to indicate baseline cortical actin organization.

### Preparation of apical out colon organoids

Human colon organoids were generated using a previously described apical out organoid preparation protocol^122^, with adaptations as outlined below. Briefly, freshly isolated human colon tissue was collected and dissected into approximately 1 mm³ fragments from distinct regions. Tissue fragments were subjected to enzymatic digestion at 37 °C to dissociate the extracellular matrix, followed by multiple washes with cold wash medium to remove debris and enrich for viable epithelial cells.

For three-dimensional expansion, isolated cells were resuspended in ice-cold Matrigel (Corning, #354234) and plated as domes into prewarmed 24-well tissue culture plates. After polymerization, organoid culture medium was added, and cultures were maintained under standard conditions to allow organoid formation and expansion. Matrigel provided extracellular matrix support essential for organoid growth and structural organization.

For passaging, organoids were mechanically collected by pipetting and enzymatically dissociated using 1× TrypLE™ Express Enzyme (phenol red–free; Gibco, #12605028) to generate smaller clusters or single cells. Following additional washing steps, cells were re-embedded in fresh Matrigel and replated for continued growth. Organoids were maintained by replacing the culture medium every 2 days to ensure adequate nutrient supply and support sustained growth.

To induce polarity inversion and generate apical-out organoids, mature apical-in organoids were collected. After supernatant removal, Matrigel was removed with 5 min 37°C incubation with digestion solution according to the protocol described by Kiss et al^122^. After the careful removal of matrigel the organoids were transferred to anti-adherence–coated culture surfaces (Anti-Adherence Rinsing Solution; StemCell Technologies, #07010) and cultured for 48 h. This suspension culture condition promoted reorganization of epithelial polarity, resulting in inversion of the apical surface from an inward-facing to an outward-facing orientation, thereby enabling direct apical accessibility for downstream physiological and infection studies.

### Infection of apical-out colon organoids with bacteria

Bacterial strains were cultured as described above. Apical-out colon organoids derived from healthy human biopsy samples were infected with YFP-labelled *Klebsiella pneumoniae* ancestor or BasR G53V mutant strains. Infections were performed in 2-ml microcentrifuge tubes. Briefly, apical-out organoids were transferred to tubes containing 750 µl of human colon organoid culture medium, and bacterial cultures were diluted in the same medium supplemented with 10 mM HEPES to a final OD₆₀₀ of 0.5. Organoids were co-incubated with bacteria for 1 h at 37 °C in a humidified atmosphere containing 5% CO₂. Organoid cell death was assessed immediately following infection using a LIVE/DEAD™ viability assay (Invitrogen, #L34955). Organoids were incubated with LIVE/DEAD™ staining solution diluted 1:1,000 in 1× PBS for 30 min at room temperature. Samples were subsequently fixed with 4% paraformaldehyde (PFA; Thermo Scientific Chemicals, #J19943.K2) prepared in 1× PBS for 12 min at room temperature. For cytoskeletal and nuclear visualization, organoids were permeabilized and stained with CellMask™ Deep Red Actin Tracking Stain (Invitrogen, #A57245) diluted 1:500 in 0.1% Triton X-100 and incubated overnight at 4 °C. Samples were stored in 1× PBS at 4 °C until imaging. Confocal z-stack images were acquired using a Leica Stellaris 8/STED confocal microscope equipped with a 63x/1.40 HC PL APO Oil CS2 objective and controlled by Microscope Imaging Software. Fluorophores were excited using diode lasers at 405, 488, and 561 nm. Maximum intensity projections were generated from z-stacks. Cytotoxicity in apical-out human colon organoids was quantified using LIVE/DEAD™ fluorescence staining followed by image-based analysis. Following infection with either the ancestor or the BasR G53V mutant, confocal images were processed in ImageJ (Fiji)^108^. For each organoid, the total projected organoid area (OrganoidArea) was defined from the segmentation of the organoid outline, and cell death was defined as the projected area positive for LIVE/DEAD™ staining (LiveDeadArea). Cytotoxicity was calculated as a size-normalized image-based metric, referred to as the cell death fraction, adapted from established organoid live/dead image-analysis workflows in which fluorescent viability or cell-death signals are normalized to the corresponding organoid area or mask^123–125^. The cell death fraction was defined as the ratio of LIVE/DEAD™-positive area to total organoid area (Cell death fraction = LiveDeadArea / OrganoidArea)^55,126–128^. This metric represents the fraction of each organoid undergoing cell death and enables comparison across organoids of varying sizes. Cell death fraction values were calculated for individual organoids and plotted as violin plots for each condition. Uninfected control organoids were included as a reference, and the mean cell death fraction of control samples was displayed as a horizontal reference line. Experiments were performed using apical-out colon organoids generated from a single healthy human colon biopsy sample. This model exposes the apical epithelial surface to bacteria, as previously described for apical-out intestinal organoid infection models^55^. Statistical comparisons between organoids infected with the ancestor strain and those infected with the BasR G53V mutant were performed using two-sided Mann– Whitney U tests. Statistical significance was defined at *p < 0.05, **p < 0.01, ***p < 0.001, ****p < 0.0001; ns, not significant.

### Field-emission scanning electron microscopy (FE-SEM)

Bacterial cells were cultured as described above. Human epithelial cell lines A549 and Caco-2 were used for infection experiments. Cells were seeded in 24-well plates at a density of 50,000 cells/well for A549 and 70,000 cells/well for Caco-2. Each well contained three sterile 5-mm glass coverslips (Epredia, Braunschweig, Germany), onto which the cells were seeded. Cells were infected with either the ancestor strain or the BasR G53V mutant strain for 1 h or 3 h, as indicated. Infection was carried out at an OD600 of 0.25 for both cell lines. Following infection, cells were washed with 1× PBS to remove non-adherent bacteria and subsequently fixed with ice-cold Karnovsky solution containing 2% paraformaldehyde (Sigma-Aldrich, St. Louis, MO, USA) and 2.5% glutaraldehyde (Polysciences, Warrington, PA, USA) in PBS overnight at 4 °C. After fixation, samples were rinsed in PBS (pH 7.4) for 10 min, then dehydrated in a graded ethanol series (from 30% to 100%) for 60 min at each concentration. Subsequently, the samples were incubated in mixtures of absolute ethanol and hexamethyldisilazane (HMDS; Sigma-Aldrich, St. Louis, MO, USA) at ratios of 3:1, 1:1, and 1:3 (v/v) for 60 min each, followed by a final incubation in pure HMDS for 60 min. The chemically dried samples were coated with a 12 nm layer of gold using a Quorum Q150T sputter coater and observed under a Zeiss Sigma 300 field emission scanning electron microscope (FE-SEM; Carl Zeiss Microscopy GmbH, Jena, Germany). Imaging was performed using a secondary electron (SE) detector at an acceleration voltage of 2 kV. For large-area scanning and high-throughput measurements, the Atlas 5 system was utilized to generate large-scale mosaics at a resolution of 10 nm/pixel, enabling the visualization of extensive surface areas at high magnification (Video S1).

### Systemic mouse infection model and tissue analysis

All animal experiments were approved by the relevant institutional animal care and use committee and animal care and handling procedures adhered to the national and the European Federation for Laboratory Animal Science Association (FELASA) guidelines (permit Reference Nr. XX./2365/2023.)

Immunocompetent female BALB/c mice (6–8 weeks old) were used for systemic infection experiments. Mice were intravenously infected via tail vein injection with mScarlet-expressing *Klebsiella pneumoniae* strains (ancestor and BasR G53V mutant, respectively) at a dose of 5 × 10⁶/animal in a total volume of 100 µL sterile Hanks’ Balanced Salt Solution (HBSS). At defined time points post-infection (0, 6, 24, and 72 h), groups of 6 mice per condition were euthanized and perfused via the vasculature with 5 mL sterile PBS to remove circulating blood with bacterial cells. Organs, including liver, lungs, intestines, and stomach, were then aseptically collected for downstream analyses.

For bacterial quantification (CFU), organs were weighed and homogenized in sterile PBS using a mechanical tissue homogenizer. Serial tenfold dilutions of homogenates were prepared and plated on CHROMagar™ Orientation agar plates on which *K. pneumoniae* can be distinguished from potential contaminants. Plates were incubated at 37 °C overnight, and CFUs were enumerated to determine bacterial load per organ. Bacterial load subsequently was normalized to tissue weight. Bacterial clearance was defined as the change in bacterial load during the duration of the experiment, in line with previous works in the field^61^, by the following formula: Clearance = log_10_(CFU_0_)-log_10_(CFU_72_).

For histological and imaging analyses, liver tissues were fixed in 4% paraformaldehyde (PFA) for 48– 72 h at 4 °C, followed by cryoprotection in 15% (w/v) sucrose in phosphate-buffered saline (PBS) for 6–12 h at 4 °C and subsequently in 30% (w/v) sucrose in PBS overnight at 4 °C. Tissues were then embedded in Cryogel (Epredia™ Cryomatrix™ #67-690-06) and optimal cutting temperature (OCT) compound, snap-frozen, and sectioned at a thickness of 7 µm using a cryostat (Thermo Fisher Scientific). Cryosections were processed for immunofluorescence staining using CellMask™ Green Actin Tracking Stain (1 µg/mL, 1:1,000 dilution; Invitrogen, #A57243) to visualize the actin cytoskeleton and Hoechst 33342 (1 µg/mL, 1:10,000 dilution; Invitrogen, #H3570) to label nuclei.”

Mouse liver sections were imaged using a Leica Stellaris 5 confocal microscope (63×/1.40 NA oil objective) in sequential line-scanning mode. Excitation and emission parameters were as follows: Hoechst 33342 (Invitrogen, #H3570; 405 nm laser, 420–495 nm), CellMask™ Green Actin Tracking Stain (Invitrogen, #A57243; White Light Laser [WLL] 498 nm; 503–572 nm), and mScarlet-expressing bacteria (WLL 570 nm; 575–814 nm). High-resolution z-stacks (∼8 µm total depth, 0.6 µm steps) were acquired at 1,024×1,024 pixel resolution, 400 Hz scan speed, 1 Airy Unit, and a line average of 2. While these baseline parameters remained constant, detector gain was optimized per field of view to maximize dynamic range and prevent pixel saturation. Maximum intensity projections and orthogonal cross-sections were generated using Imaris 11 (Oxford Instruments) and Fiji/ImageJ^108^. Three-dimensional (3D) video animations were rendered using the Imaris 11 blend mode.

### Pulmonary mouse infection model and tissue analysis

Mouse pulmonary infections and all subsequent experiments were performed by Link-U Biotech Inc. on behalf of Eurofins Discovery Ltd. in Taiwan. Animal care and handling procedures adhered to the national guidelines and international recommendations. Immunocompetent female BALB/c mice (6– 8 weeks old) were used for pulmonary infection experiments. Mice were infected via intratracheal injection with mScarlet-expressing *Klebsiella pneumoniae* strains (ancestor and BasR G53V mutant, respectively) at a dose of 3 × 10^8^/animal. At 48 hours post-infection groups of 4 mice per condition were euthanized.

Bronchoalveolar Lavage Fluid (BALF) Collection was performed as follows: 0.5 mL PBS was administered once through a tracheal cannula, after which approximately 0.2 to 0.3 mL of BALF was recovered. The BALF was centrifuged at 2,500 × g for 15 minutes at 4°C. The supernatant was then collected and stored at -80°C for subsequent cytokine analysis.

Three Abcam ELISA kits were used to measure cytokine levels: TNF-α (Cat#208348), IL-6 (Cat# 222503), and IL-10 (Cat# 255729). Assays were performed according to the manufacturer’s instructions. Briefly, BALF samples were thawed and diluted using the provided sample diluent. Standards were reconstituted, vortexed, and serially diluted to generate a calibration curve. Samples and standards (50 µL) were incubated with 50 µL of an antibody cocktail (comprising capture and detector antibodies) at room temperature on a shaker (400 rpm) for 1 hour. After being washed three times, the wells were incubated with 100 µL of tetramethylbenzidine (TMB) development solution in the dark on a shaker (400 rpm) for 5–20 minutes. Subsequently, 100 µL of stop solution was added and mixed gently. The assay plate was read at OD450 nm within 30 min of adding the stop solution using a microplate reader (Tecan). The calibration curve was generated by plotting the absorbance values against the standard concentrations and fitting the data to a four-parameter logistic (4PL) regression model.

Lung samples were harvested and then preserved in 10% neutral buffered formalin (NBF) after necropsy. The lung tissues (comprising five lobes) were prepared for histopathological examination. They were sectioned at 4-6 μm and stained with hematoxylin and eosin (H&E). Finally, samples were examined through an optical microscope (Nexcope®, NE700). Histopathological scoring was performed based on an established semi-quantitative scoring system^129^.

## Acknowledgments

General

We are grateful for the possibility to use the GenAI4Science service of the HUN-REN Cloud (Héder et al., 2022; https://science-cloud.hu/)^130^ to standardize the host and isolation source metadata of the analyzed genomes.

## Author contributions

Conceptualization: PS, MSC, ZF, LD, SzJ, CP

Methodology: PS, MSC, ZF, LD, BT, IL, AE, NK, ZCB, EK, GB, MP, ZsT, ZV, DA, CGP, ZS, EV, TM, JM, FA, AF, RT, BC

Investigation: PS, MSC, ZF, LD, BT, EszK, AZV, MC, RK, EM, GI, IL, LM, AS, DN, NZsF, GG, BP, ZCB, EK, GB, MP, IG, AP, ZV, DA, CGP, ZS, EV, TM, FA, AF, RT, BK, VV, BC

Visualization: PS, MSC, ZF, LD, IL, GI, AS, GG, FA, AF,

Funding acquisition: BKr, ISz, ZsT, AG, SzJ, CP

Project administration: AD, ISz, SzJ, CP

Supervision: BP, BK, LH, SzJ, CP

Writing – original draft: PS, MSC, ZF, LD, BP, SzJ, CP

Writing – review & editing: PS, MSC, ZF, LD, AD, IL, BP, ZCB, EK, ISz, ZsT, AG, JM, BK, SZJ, CP

## Data, code, and materials availability

The data supporting the findings of this study are available in the article and its Supplementary Information. Source data for the main figures are provided with this paper, while the supplementary figures can be reproduced using the accompanying Supplementary Datasets. Whole genome sequencing data of the adapted lines used in this study have been previously deposited in the European Nucleotide Archive (ENA) at EMBL-EBI under accession number PRJEB63210 (https://www.ebi.ac.uk/ena/browser/view/PRJEB63210). RNA-seq reads for this study have been deposited in the European Nucleotide Archive (ENA) at EMBL-EBI under accession number ERP203279.

The clinical isolates featured in this study in this study are deposited in the bacterial collection of HUN-REN Biological Research Centre (https://hun-ren.hu/en/research-network/research-centers/hun-ren-biological-research-centre-szeged). As specified in Material Transfer Agreements, these isolates and their derivatives cannot be transferred to a third party as they can be used “only at the recipient organizations and only in the recipient scientists’ laboratories under the direction of the recipient scientists or others working under their direct supervisions.”

