## Supplementary Materials for "Laboratory resistance evolution discovers virulence risks of next-generation antibiotics"

**The PDF file includes:**

Supplementary Note 1-6
Supplementary Figs 1-31
References

**Other Supplementary Materials for this manuscript include the following:**

Supplementary Video 1-2
Supplementary Data 1-6

### Supplementary Notes

***Supplementary Note 1. Structurally diverse antimicrobial peptides and small molecules targeting the Gram-negative cell envelope applied in this study***

"Membrane targeting" antimicrobial compounds used in this study span distinct structural classes and modes of interaction with the envelope (Fig. 1). Polymyxin derivatives (colistin, polymyxin B, SPR-206, QPX9003) are cyclic lipopeptides composed of a cationic peptide ring and an N-terminal fatty acyl tail, enabling high-affinity binding to lipid A, leading to the disruption of outer membrane (OM) and later inner membrane (IM) integrity[1,2]. POL-7306 is a chimeric peptidomimetic antibiotic in which a β-hairpin macrocycle is linked to the peptide macrocycle of polymyxin, similarly targeting lipid A while also inhibiting the essential OM protein BamA[3]. Tridecaptin-M152-P3 is a linear non-ribosomal lipopeptide that binds lipid II and lipid A, creating proton-specific pores in the IM leading to the disruption of membrane potential, this blocking ATP synthesis[4]. SCH79797 is a small-molecule antibiotic with dual activity, inhibiting folate biosynthesis and the mechanosensitive channel MscL, thereby perturbing both intracellular metabolism and membrane homeostasis[5,6].

Antimicrobial peptides (AMPs) in this study exhibit diverse structural features and membrane activities. LL-37 is a linear, amphipathic α-helical host-defense peptide that disrupts membranes via a carpet-like mechanism dissimilar to polymyxin derivatives[7,8]. SCUB3_MLP22 and PPP1CB-derived peptides are short, cationic peptides with predicted amphipathic character, whereas pep19mod is a synthetic peptide enriched in hydrophobic and basic residues with anti-endotoxin properties; all induce membrane perturbation through unresolved mechanisms[9–11]. GKY25, a thrombin-derived linear peptide, binds lipid A and interferes with LPS-mediated membrane stability similar to polymyxins[12].

Finally, inhibitors of LPS biosynthesis target the committed step of the Raetz pathway. CHIR-090 is a hydroxamate-based LpxC inhibitor, whereas TP0586532 represents a structurally distinct, non-hydroxamate inhibitor of LpxC, both blocking lipid A synthesis and thereby compromising OM assembly[13,14].

***Supplementary Note 2. Prevalence of laboratory-observed virulence-associated substitutions in natural populations***

In our previous work, we demonstrated that mutations arising during laboratory evolution under antibiotic selection frequently overlap with variants present in environmental and clinical bacterial populations[15]. Given the enrichment of mutations in virulence-associated genes among antibiotic-adapted lines, we next examined whether these substitutions are detectable in natural isolates and whether they are preferentially associated with clinically relevant lineages.

**Genome datasets and mutation screening.** We analyzed 189 non-synonymous amino acid substitutions identified in virulence-associated genes from laboratory-evolved *E. coli* and *K. pneumoniae* lines[15]. These substitutions were screened against publicly available genome collections comprising approximately 394,900 *E. coli* and 72,400 *K. pneumoniae* natural isolates[16]. Using the full datasets, 70.9% (61 / 86) of the substitutions identified in *E. coli* and 43.7% (45 / 103) of those identified in *K. pneumoniae* were detected in at least one natural isolate. To account for unequal sampling depth between species, we performed random subsampling of 40,000 genomes per species (5,000 replicates, without replacement within replicates). After standardization, the expected detection rate was 38.4% for *E. coli* (95% CI: 30.2–45.3%) and 38.8% for *K. pneumoniae* (95% CI: 34.9–41.7%). Accordingly, the difference between species was no longer detectable (mean difference = 0.4 percentage points; 95% CI: −8.0 to 9.0%).

**Association with clinical infection-associated isolates.** To determine whether these substitutions were preferentially associated with clinical infection-associated strains, we classified *E. coli* isolates by isolation source, using metadata when available (N = 276,822, table S1, Methods). Isolates recovered from blood, nervous system, reproductive, respiratory, urinary, wound, or other infection sites were classified as clinical infection-associated (N = 40,721), whereas the remainder were considered carriage, environmental or other non-infection-associated (N = 236,101).

To minimize bias arising from gene absence, we restricted the analysis to virulence-associated genes present in ≥85% of genomes. Within this filtered set, substitutions were detected in 192 clinical infection-associated isolates and 335 carriage, environmental or other non-infection-associated isolates, whereas 35,536 clinical infection-associated and 195,729 carriage, environmental or other non-infection-associated isolates lacked such substitutions. This corresponds to a significant enrichment among clinical infection-associated isolates (Fisher's exact test, OR = 3.15, $P < 2.2 \times 10^{-16}$). Gene-level analysis indicated that this enrichment was not uniformly distributed but driven by recurrent substitutions in a limited set of loci, including *pmrB/basS*, *basR*, *acrB*, and *fimH*, each of which was individually enriched in clinical infection-associated isolates.

We performed the same analysis for *K. pneumoniae*, classifying isolates by isolation source where metadata were available (N = 43,302, table S1). Of these, 28,927 were categorized as clinical infection-associated and 14,375 as carriage, environmental or other non-infection-associated. Within the ≥85% gene-presence subset, substitutions were detected in 277 clinical infection-associated isolates and 77 carriage, environmental or other non-infection-associated isolates, while 21,814 clinical infection-associated and 11,356 carriage, environmental or other non-infection-associated isolates lacked such substitutions. This also revealed significant enrichment among clinical infection-associated isolates (Fisher's exact test, OR = 1.8, $P < 2.2 \times 10^{-8}$). At the gene level, enrichment was primarily driven by a substitution in *basS*, while additional substitutions were observed exclusively in clinical infection-associated isolates (without significant enrichment due to low counts), including those in *phoQ, sbmA* (transporter for microcin-type bacteriocins), *csrA* (mediates global changes in gene expression, shifting from rapid growth to stress survival by linking envelope stress, the stringent response, and the catabolite repression systems), *basR*, and *mprA* (regulator of capsule production).

**Distribution across high-risk clonal lineages.** To evaluate whether these substitutions occurred within clinically relevant *K. pneumoniae* lineages, we used Kleborate, a genome-based framework that infers lineage and assigns sequence type (ST) together with composite resistance and virulence scores (0–5) based on whole-genome sequencing data[17]. The substitutions that emerged in virulence-associated genes were distributed across multiple globally prevalent sequence types. Several of these STs correspond to recognized high-risk lineages characterized by elevated resistance and/or virulence scores, including ST2096[18,19], ST11[20], ST111, ST395[21], ST43, ST383[22,23], and ST2502 (fig. S3). Among these, ST2096 has emerged as a lineage combining high resistance and virulence, particularly in Saudi Arabia and India. ST395 is frequently linked with OXA-48 carbapenemase dissemination and has been implicated in clinical outbreaks in the United Kingdom. ST383 is known for the accumulation of multiple resistance determinants and involvement in clinical outbreaks in Southern Europe, including Greece and Italy.

**Conclusion.** The recurrence of identical non-synonymous substitutions across large collections of natural genomes, their enrichment among clinical infection-associated *E. coli* isolates, and their presence within globally disseminated high-risk *K. pneumoniae* lineages indicate that virulence-associated mutations selected during laboratory evolution also arise in clinical and environmental populations. These findings support the evolutionary relevance of laboratory-observed virulence-associated mutations beyond controlled experimental conditions.

### Supplementary Note 3. Estimating relative virulence from Galleria survival curves

Host-killing dynamics and relative virulence were quantified from Galleria survival experiments using Kaplan-Meier visualization and Cox proportional-hazards modelling. All analyses were performed in R using the survival and survminer packages[24,25].

The statistical framework is summarized in Fig. SN1A. To compare each adapted line directly with its corresponding ancestor, we used the Cox proportional-hazards model. This semiparametric survival model relates time to larval death to explanatory variables, here the infecting strain[26,27]. In this framework, the hazard is the instantaneous risk of larval death among larvae still alive at a given time. The model compares the strain-specific hazard with the ancestor hazard under the proportional-hazards assumption. This can be written as $h(t \mid strain) = h0(t) \times exp(beta)$, where $h(t \mid strain)$ is the hazard at time t for larvae infected with a given strain, h0(t) is the baseline hazard at time t, exp denotes exponentiation, and beta is the Cox model coefficient estimating the difference between the adapted line and its corresponding ancestor. Thus, a strain with increased host killing has a higher instantaneous death hazard relative to its ancestor, represented as a hazard ratio rather than as a raw killing rate or a slope over time.

The relationship between the Cox coefficient and the hazard ratio is shown in Fig. SN1B. The Cox coefficient is the log hazard ratio: coef = log(HR). Conversely, the hazard ratio is obtained by exponentiating the coefficient: HR = exp(coef). A coefficient of 0 corresponds to HR = 1, indicating no difference from the ancestor. Positive coefficients indicate higher instantaneous death hazard and shorter survival, whereas negative coefficients indicate lower instantaneous death hazard and longer survival. For example, a coefficient of 2 corresponds to HR = exp(2) = 7.39, not a twofold increase and not a two-order-of-magnitude increase.

The survival data themselves were visualized using Kaplan-Meier curves for each Galleria experiment (Fig. SN1C). A Kaplan-Meier curve is a non-parametric estimate of the survival function, in this case the probability that larvae remain alive over time[28,29]. The curve is a step function that drops each time a death event occurs. In the example shown, larvae infected with the BasR G53V mutant show reduced survival relative to larvae infected with the ancestor. The 95% confidence interval and the p-value from the log-rank test, also known as the Mantel-Cox test, are displayed together with the Kaplan-Meier curves.

For each adapted-line-versus-ancestor comparison, relative virulence was reported as the Cox model coefficient, coef (Fig. SN1D). The corresponding hazard ratio is exp(coef). Statistical significance was assessed separately for each adapted line by testing whether its survival curve differed from that of its corresponding ancestor in a Cox proportional-hazards model. The p-value for each adapted-line-versus-ancestor comparison was obtained by a likelihood-ratio test. The resulting p-values were corrected across tested strains using the Benjamini-Hochberg procedure, with a 10% false-discovery-rate threshold used to classify lines as increased virulence, decreased virulence, or no significant change.

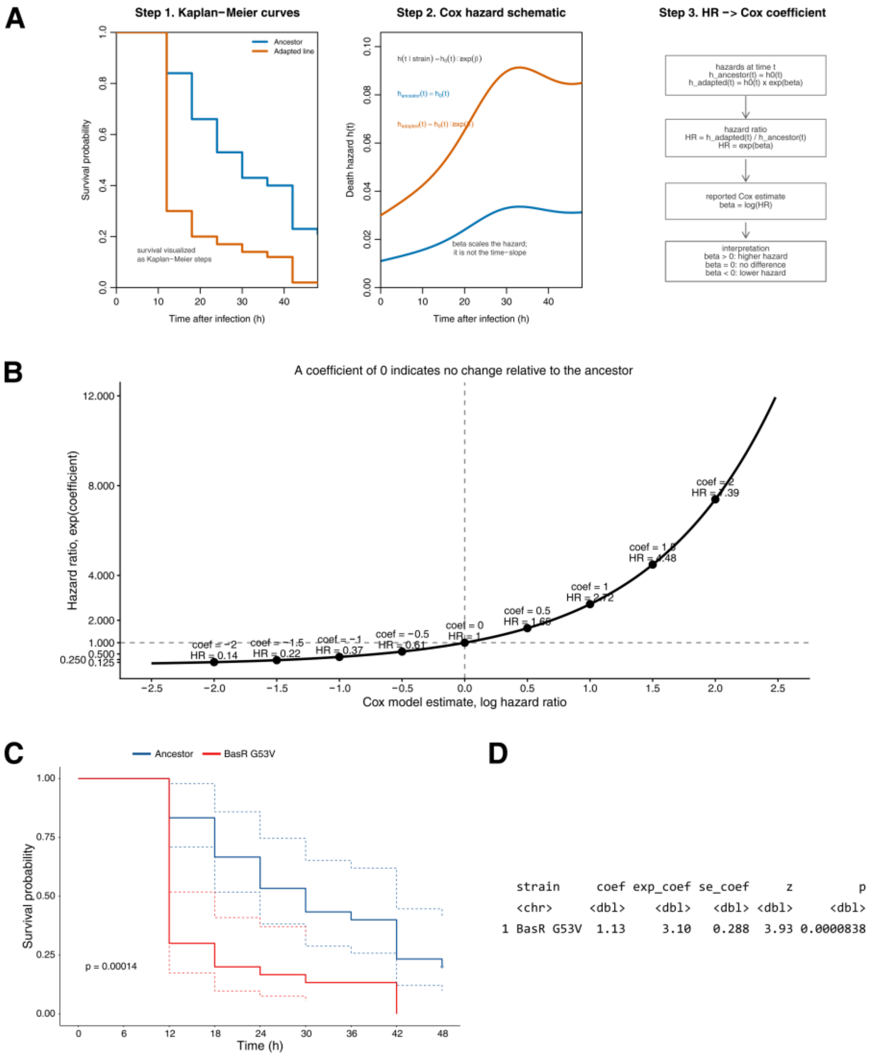


**Fig. SN1. Estimating relative virulence from *Galleria mellonella* survival data**. **(A) Conceptual illustration of the Cox proportional-hazards model used to compare each adapted line with its corresponding ancestor.** The model describes the strain-specific hazard as h(t | strain) = h0(t) × exp(beta), where h(t | strain) is the hazard at time t for larvae infected with a given strain, h0(t) is the baseline hazard at time t, exp denotes exponentiation, and beta is the Cox model coefficient estimating the difference between the adapted line and its corresponding ancestor. Under this model, the adapted-line hazard is represented as a multiplicative scaling of the ancestor hazard. **(B) Relationship between Cox model coefficients and hazard ratios.** The x-axis shows the Cox model coefficient (coef) on the log-hazard scale, and the y-axis shows the corresponding hazard ratio (HR) obtained by exponentiating the coefficient: HR = exp(coef). A coefficient of 0 corresponds to HR = 1, indicating no difference from the ancestor. Positive coefficients indicate higher instantaneous hazard of larval death, whereas negative coefficients indicate lower hazard. For example, a coefficient of 2 corresponds to HR = exp(2) = 7.39. Displaying Cox model effects on the log-hazard scale is standard because reciprocal hazard-ratio effects are symmetric around zero on this scale. **(C) Example**

**Kaplan-Meier survival curves comparing larvae infected with the ancestor and the BasR G53V mutant over 48 h.** Solid lines show Kaplan-Meier survival estimates, dotted lines show 95% confidence intervals, and the displayed p-value is from the log-rank test, also known as the Mantel-Cox test. This p-value is shown as part of the Kaplan-Meier visualization. **(D) Cox model summary for the BasR G53V-versus-ancestor comparison.** The regression coefficient, coef, is the reported relative virulence estimate and equals log(HR), where HR is the hazard ratio relative to the ancestor. The corresponding hazard ratio is shown as exp(coef). The p-value in this panel is calculated from the Cox proportional-hazards model and was the value used for statistical classification of adapted lines. Positive coefficients indicate higher instantaneous death hazard relative to the ancestor. Abbreviations: se(coef), standard error of the coefficient; z, Wald statistic; p, p-value.

***Supplementary Note 4. Contribution of lipid A remodeling features to virulence variation***

To formally assess the relative contribution of distinct Lipid A remodeling pathways to virulence evolution, we fitted ordinal regression models including antibiotic class, the presence of Lipid A–related mutations, and all measured lipid A modification features (see Methods). A reduced primary model, incorporating single- and double aminoarabinose, phosphoethanolamine-aminoarabinose modifications, together with increased hexa-acylation, explained the vast majority of virulence-associated variation (~83.7% of residual deviance, likelihood ratio test, $P = 1.3 \times 10^{-6}$). In contrast, phosphoethanolamine modifications alone (secondary model) explained ~7.75% of the residual deviance, while intermediate acylation states (tri-, tetra-, and penta-acylation, third model) explained ~22.9%. Of note, hepta-acylated lipid A showed no variance across strains and was therefore uninformative. Importantly, adding all remaining lipid A features did not significantly improve the primary model fit (ANOVA, $P = 0.09$), indicating that a restricted subset of lipid A remodeling features captures the majority of virulence-associated variation. The full model incorporating all modifications explained ~88.5% of the residual deviance in virulence outcomes. The robustness of the association between primary lipid A remodeling features and virulence was further supported by permutation testing of virulence labels (20,000 permutations; $P = 0.00159$), indicating that the observed linkage is unlikely to arise by chance.

### ***Supplementary Note 5. Convergent regulatory routes to lipid A remodeling in hypervirulent adapted lines***

Although the 13 adapted *K. pneumoniae* lines with increased virulence carried divergent resistance mutations (Fig. S8), they converged on a common outer-membrane remodeling phenotype characterized by increased LAra4N modification of lipid A (Fig. 3B, Fig.S9). The regulatory framework summarized in Fig. 6 suggests that this convergence can arise through an interconnected membrane-stress network centered on PhoPQ and BasSR, together with auxiliary regulators that modulate signal flow into this axis.

BasSR provides the most direct route to the observed phenotype. Activation of BasR promotes expression of lipid A modification functions, including ArnT, which catalyzes L-Ara4N transfer to lipid A, and can also enhance *eptA* expression, leading to phosphoetanolamine modification of lipid A[30]. In *K. pneumoniae*, signal integration between PhoPQ and BasSR is further reinforced by PmrD, which can protect phosphorylated BasR from dephosphorylation, thereby linking PhoP-dependent envelope sensing to the BasR-controlled lipid A remodeling program[31].

Our data is consistent with activation of this interconnected module from multiple upstream entry points. We detected mutations in *phoP, rstB, qseC, basR, plsC*, and *ackA* in adapted lines displaying increased virulence (Fig. S8), and increased expression of *phoP, basS, basR, arnT*, and *eptA* relative to lines without increased virulence (Fig. S9). This pattern supports a model in which genetically distinct resistance mutations converge on a shared transcriptional output rather than a single common mutation.

Several of the mutated loci plausibly tune membrane-stress signalling. PhoQ is a major sensor of envelope perturbation, whereas RstAB appears to functionally intersect with PhoPQ and may amplify PhoP-dependent transcriptional responses under stress conditions[30,32–34]. QseC provides another potential input: in related *Enterobacterales*, QseBC engages in non-cognate cross-talk with PmrAB, suggesting that perturbation of *qseC* could indirectly reshape BasR-dependent transcription[35,36].

Altered membrane composition may provide an additional layer of convergence. PlsC contributes to phospholipid biosynthesis, and changes in phospholipid composition could modify the physicochemical state of the inner membrane, thereby influencing the activity of membrane-embedded sensors such as PhoQ, RstB, QseC, and BasS. In this context, *plsC* mutations may act indirectly by biasing the envelope toward a persistent stress-like signalling state[37,38].

For *ackA*, a more plausible connection may be through acetyl-phosphate-dependent acetylation rather than phosphodonation. Prior work has shown that acetylation of PhoP can inhibit its phosphorylation, linking central metabolism to two-component signalling. Thus, mutations affecting the Pta-AckA axis may alter intracellular acetyl-phosphate homeostasis and thereby shift PhoP activity, with downstream consequences for the PhoP-PmrD-BasR regulatory cascade and lipid A remodeling output[39–41].

Together, these observations support a model in which distinct resistance mutations converge on a shared envelope-remodeling state through interconnected regulatory routes (Fig. 6). Despite mutational heterogeneity (Fig. S8), the hypervirulent lines consistently show transcriptional and biochemical hallmarks of increased BasR-linked lipid A modification, providing a mechanistic explanation for the recurrent elevation of L-Ara4N across independently evolved backgrounds (Fig. 3B, Fig.S9).

### ***Supplementary Note 6. Computational pipeline for quantitative mapping of radial actin organization.***

*Image-analysis pipeline for radial actin quantification*

F-actin organization was quantified using stepwise image-analysis workflow applied to individual infected A549 epithelial cells. Cell boundaries were first defined from raw fluorescence images using Cellpose-based segmentation, providing the reference mask for all downstream spatial analyses (Fig. SN2, panel 1)[42]. Curvilinear filament structures were enhanced using multiscale Hessian-based Sato tubeness filtering to selectively emphasize actin bundles while suppressing diffuse cytoplasmic background (panel 2)[43]. The filtered signal was subsequently thresholded to generate a binary filament mask (panel 3), followed by topological skeletonization to reduce filament networks to one-pixel-wide centerlines while preserving filament continuity and connectivity (panel 4)[43]. Overlay of the filtered signal and extracted skeleton enabled simultaneous visualization of filament distribution and structural prominence (panel 5).

To quantify intracellular spatial organization, the cytoplasm was partitioned using nuclear-anchored radial zoning into ten normalized concentric bins extending from the nuclear boundary ($r = 0$) to the cell periphery ($r = 1$) (panel 6)[44]. These bins were grouped into perinuclear ($r = 0–0.33$), intermediate ($r = 0.33–0.66$), and peripheral ($r = 0.66–1.0$) regions for biological interpretation. Mean actin bundle intensity was calculated across radial distance to generate a radial profile of filament enrichment (panel 7), where increasing signal toward the cell edge reflects peripheral actin accumulation. Final quantitative outputs included mean bundle density by radial zone, peripheral enrichment ratios (peripheral/perinuclear, peripheral/intermediate, intermediate/perinuclear), and a skeleton fidelity score used for quality filtering (panel 8)[45,46].

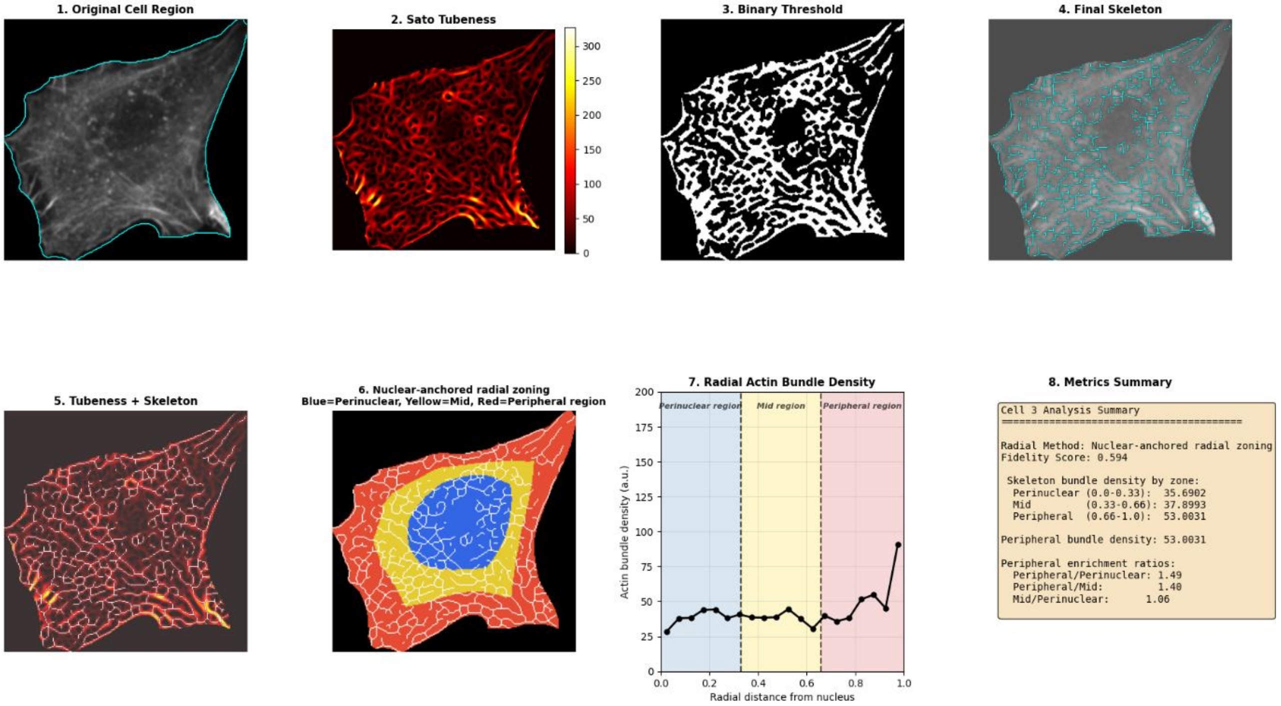


**Fig. SN2. Image-analysis workflow for quantifying radial F-actin organization in infected A549 cells.** Representative stepwise pipeline showing cell segmentation, filament enhancement, binary thresholding, skeletonization, radial zoning, intensity profiling, and extraction of quantitative metrics describing intracellular actin bundle distribution.

*Computational pipeline for peripheral actin quantification*

Quantification of peripheral actin organization was performed using a computational pipeline integrating deep-learning-based segmentation, filament extraction, and radial spatial analysis (Fig. SN3). Input confocal Z-stacks acquired on the Opera Phenix high-content imaging platform were separated into two parallel processing streams corresponding to nuclear segmentation and actin filament quantification.

The DAPI channel was processed using a customized Cellpose deep-learning model to generate nuclear and whole-cell masks. These masks were used to define nuclear-anchored radial zoning by partitioning each cell into ten normalized concentric bins extending from the nuclear boundary ($r = 0$) to the cell periphery ($r = 1$). This framework enabled the classification of cytoplasmic regions into perinuclear and peripheral compartments for downstream spatial quantification.

In parallel, the far-red F-actin channel was converted into a two-dimensional maximum-intensity projection, followed by noise reduction and background subtraction to improve filament detection. Filamentous actin structures were enhanced using Hessian-based Sato tubeness filtering, which selectively emphasizes curvilinear bundle-like structures while suppressing diffuse cytoplasmic background. Ridge-detection-based skeletonization was subsequently applied to generate a binary skeleton representation preserving filament continuity and network topology.

Quantitative analysis then proceeded through two complementary downstream paths. Path A quantified actin skeleton intensity as a measure of filament structural abundance. Path B quantified mean actin bundle density by measuring Sato filter signal along skeletonized filaments and normalizing this value to the area of each radial bin, providing a spatially resolved estimate of filament enrichment. Both outputs were subsequently used for regional ratio calculations comparing peripheral and perinuclear compartments, generating the final quantitative metric of peripheral actin enrichment.

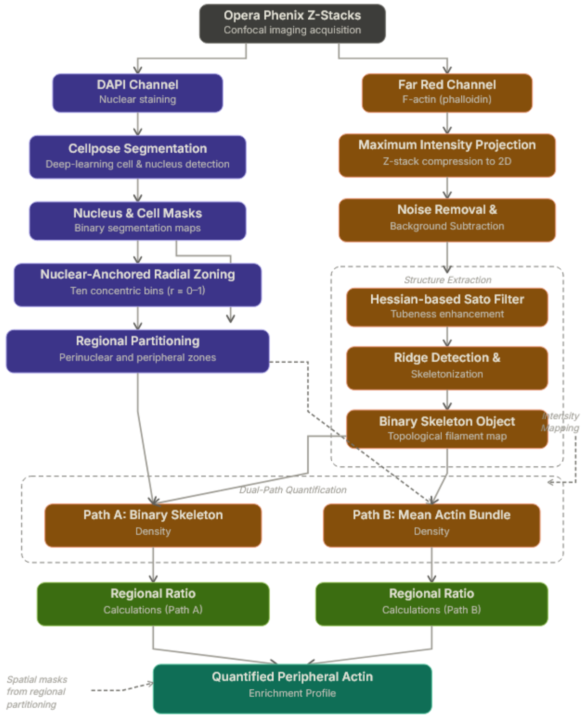


**Fig. SN3. Computational workflow for quantitative peripheral actin analysis.** Schematic overview of the full analysis pipeline used for quantifying peripheral actin enrichment from Opera Phenix Z-stack images, including segmentation-based radial zoning, filament extraction, skeletonization, and dual-path quantification of actin organization.

### Extended Data Figures

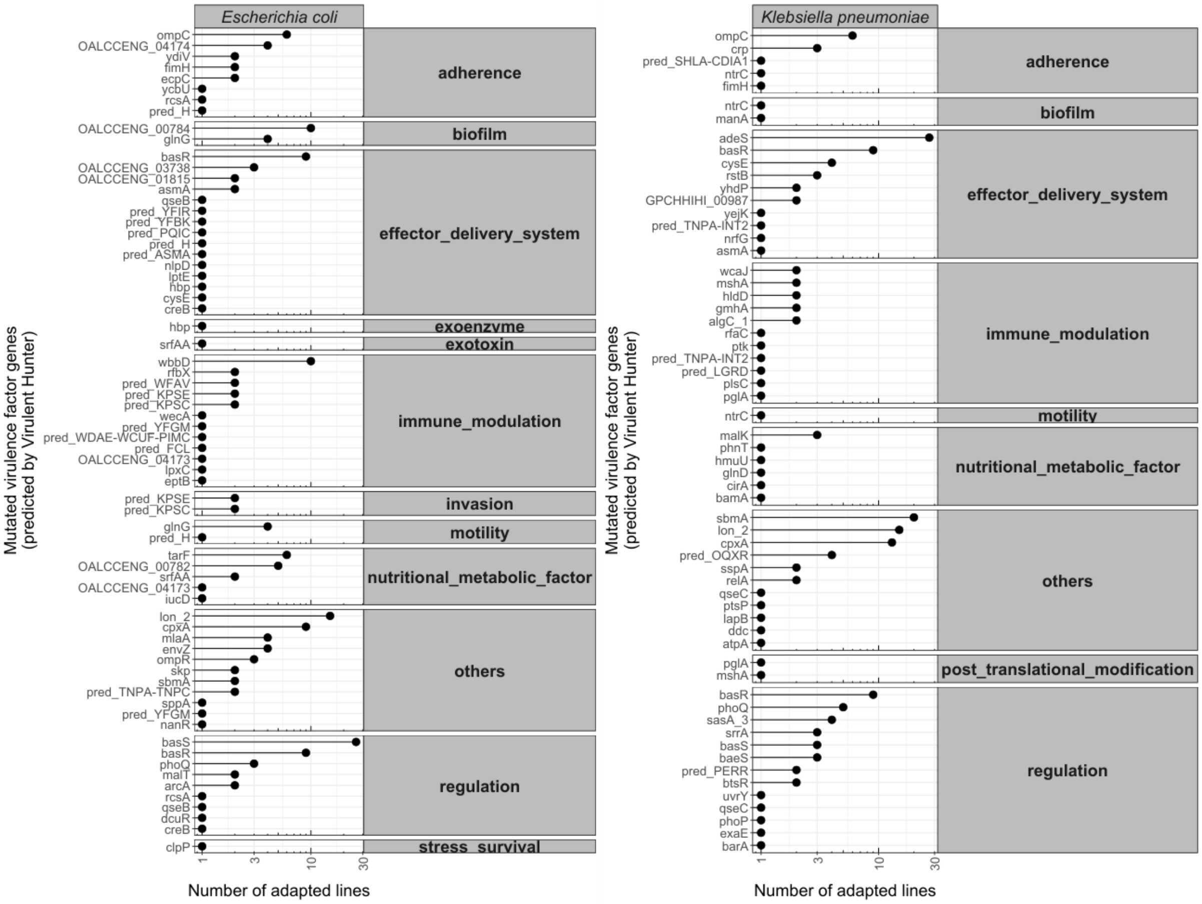


**Figure S1. Putative virulence-associated genes mutated in antibiotic-adapted *E. coli* and *K. pneumoniae* lines.** The figure shows virulence-associated genes that acquired mutations during laboratory adaptation to antibiotics in *E. coli* (left) and *K. pneumoniae* (right). Virulence-associated genes were identified using VirulentHunter, a deep-learning framework trained on curated virulence factor databases. Each row corresponds to a gene, grouped by functional virulence category (indicated on the right), and each dot represents the number of independently adapted lines in which mutations in that gene were detected. Functional virulence categories include adherence, biofilm formation, effector delivery systems, immune modulation, invasion, motility, nutritional and metabolic factors, regulation, stress survival, and unclassified functions (“others”).

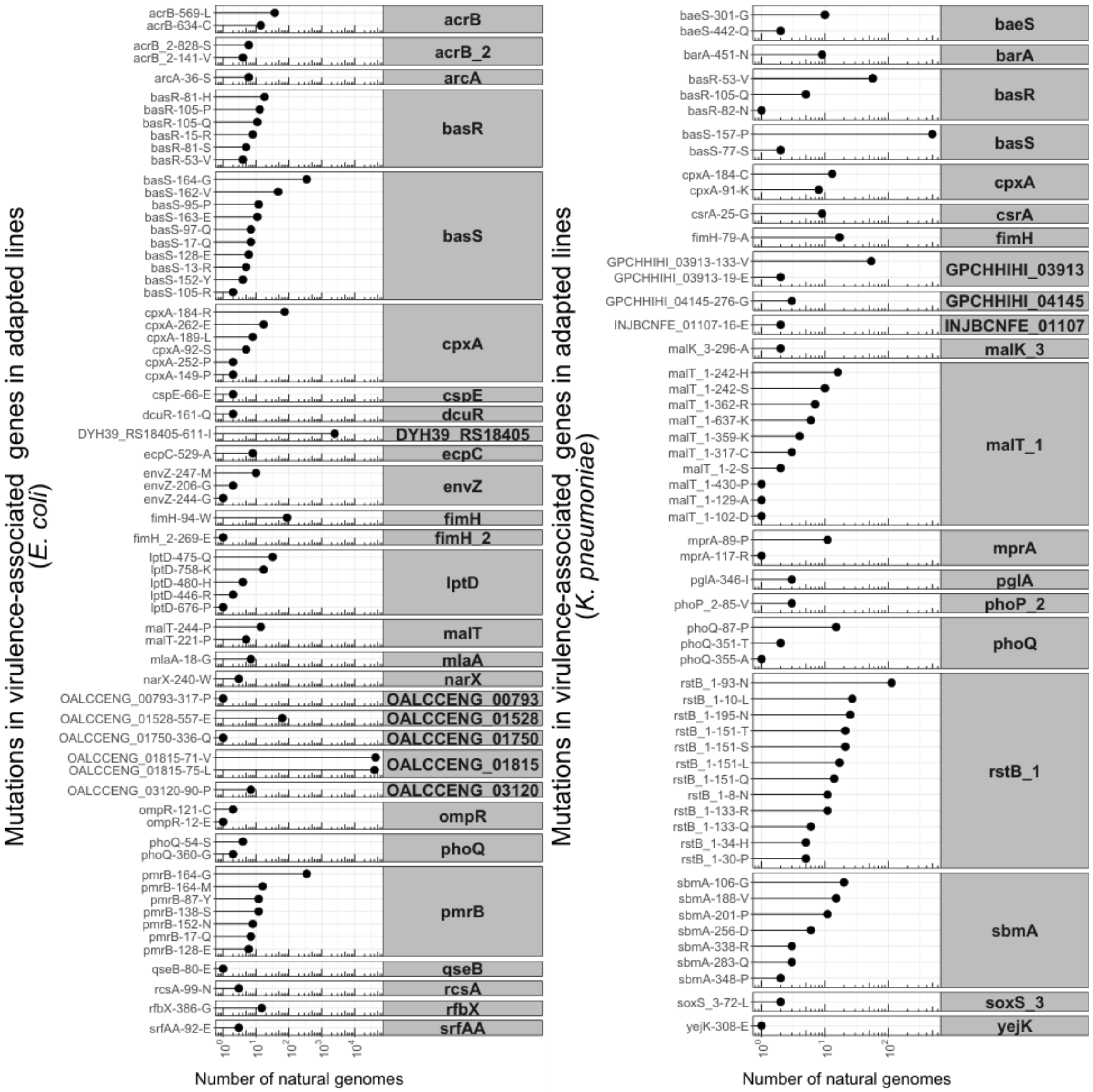


**Figure S2. Prevalence of laboratory-observed non-synonymous mutations in virulence-associated genes among natural populations.** The figure displays the frequency of laboratory-identified non-synonymous substitutions in virulence-associated genes across publicly available natural genomes of *E. coli* (left) and *K. pneumoniae* (right). Each row corresponds to a specific substitution, grouped by gene (gene names shown on the right). Black dots indicate the number of natural genomes in which the corresponding substitution was detected. Values are plotted on a logarithmic scale (x-axis).

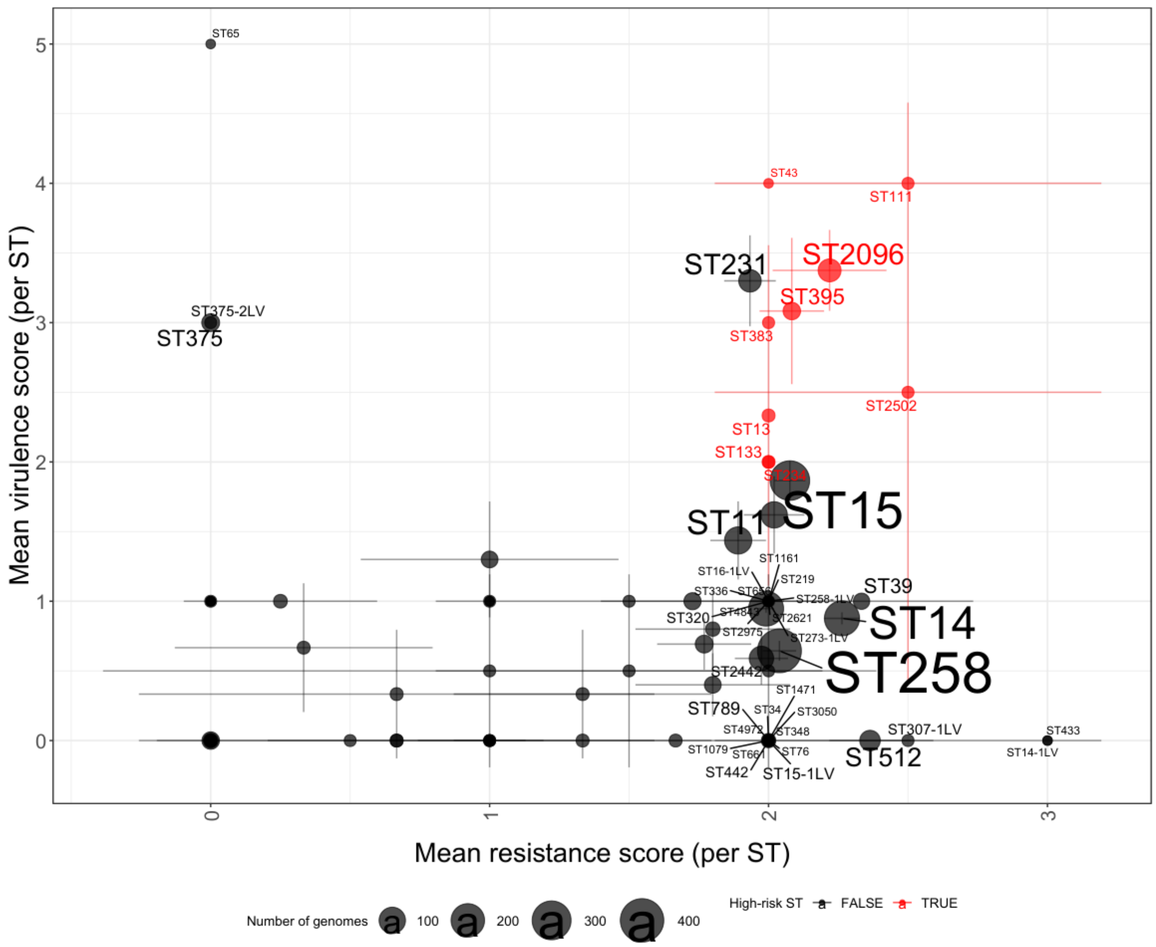


**Figure S3. Risk analysis of natural *K. pneumoniae* sequence types carrying laboratory-identified virulence-associated mutations.** The scatterplot shows the mean resistance score (x-axis) and mean virulence score (y-axis) per sequence type (ST), calculated across all genomes belonging to that ST that carry at least one virulence-associated mutation identified in laboratory-evolved lines. Scores are discrete composite indices calculated by a bioinformatics tool for genomic typing and risk profiling of *K. pneumoniae* strains (Kleborate )[17]. These indices summarize the presence of key virulence loci (e.g., siderophores, hypermucoidy-associated genes) and acquired resistance determinants, respectively, with higher values indicating greater virulence or resistance potential. Each point represents an individual ST. Horizontal and vertical error bars indicate 95% confidence intervals of the mean resistance and virulence scores, respectively. Point and label sizes are proportional to the number of genomes with laboratory-identified mutations assigned to each ST, reflecting lineage prevalence in the dataset. High-risk clonal lineages, defined as STs with mean resistance and/or mean virulence scores ≥ 2, are highlighted in red, while other STs are shown in grey. STs exceeding this threshold for either resistance or virulence score are labeled.

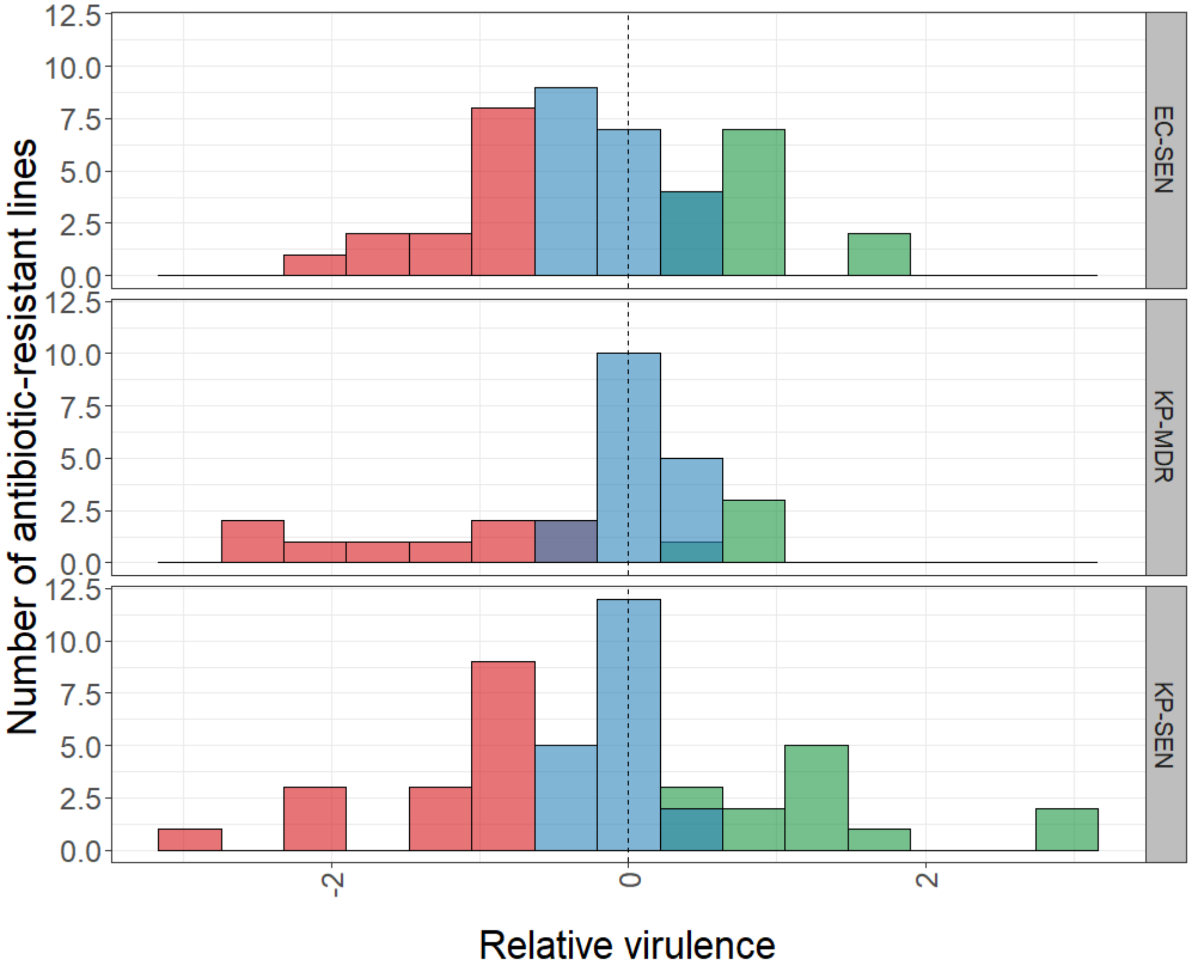


**Figure S4A. Distribution of host-killing rate of antibiotic-adapted lines in a *Galleria mellonella* infection model across genetic backgrounds.** The histogram shows the distribution of antibiotic-adapted lines derived from *E. coli* and *K. pneumoniae* (n = 124) across certain relative virulence values, where relative virulence was estimated as the host-killing rate compared to the corresponding ancestor, described by the Cox proportional hazards model (Supplementary Note 3). A relative virulence of 0 denotes a killing rate equivalent to the ancestor (vertical dashed line). Colors indicate virulence classification: increased (green) and decreased (red) virulence were assigned based on the direction of the Cox-model estimate together with statistical significance after 10% FDR correction (adjusted $P < 0.1$, likelihood-ratio test), whereas nonsignificant lines were classified as no change (blue). Strain backgrounds (panels): EC-SEN, *Escherichia coli* ATCC 25922; KP-SEN, *Klebsiella pneumoniae* ATCC 10031; KP-MDR, *Klebsiella pneumoniae* ATCC 700603. In total, 24.2% of the lines exhibited increased virulence, whereas 30.6% showed decreased virulence (Likelihood-ratio test, FDR-adjusted $P < 0.1$).

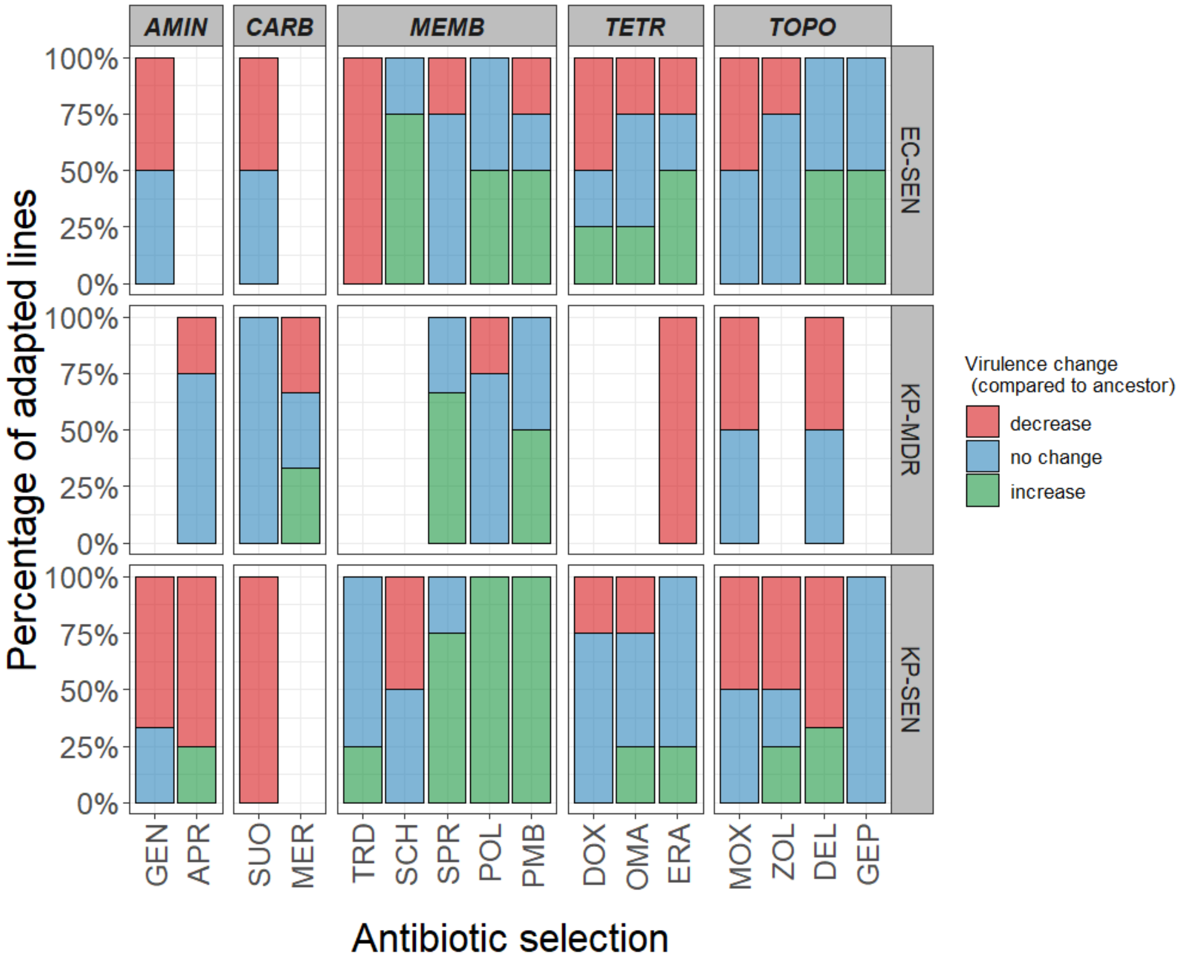


**Figure S4B. Distribution of virulence change categories across laboratory-evolved lines exposed to different antibiotics.** Relative virulence was defined as the host-killing rate in the invertebrate infection model, *G. mellonella,* relative to the corresponding ancestral strain, described by the Cox proportional hazards model (Supplementary Note 3). The stacked barplots show the percentage of adapted lines displaying decreased virulence, no change in virulence, or increased virulence compared with their ancestor. Panels are separated by antibiotic class (columns) and ancestral genetic background (rows). Missing bars indicate cases where no adapted lines were available, due to intrinsic resistance of the corresponding genetic background to that antibiotic. Strain backgrounds: EC-SEN: *Escherichia coli* ATCC 25922, KP-SEN: *Klebsiella pneumoniae* ATCC 10031, KP-MDR: *Klebsiella pneumoniae* ATCC 700603. Antibiotic classes: TOPO, topoisomerase inhibitors; TETR, tetracyclines; AMIN, aminoglycosides; CARB, carbapenems; MEMB, membrane-targeting antibiotics. For antibiotic abbreviations, see Table 1.

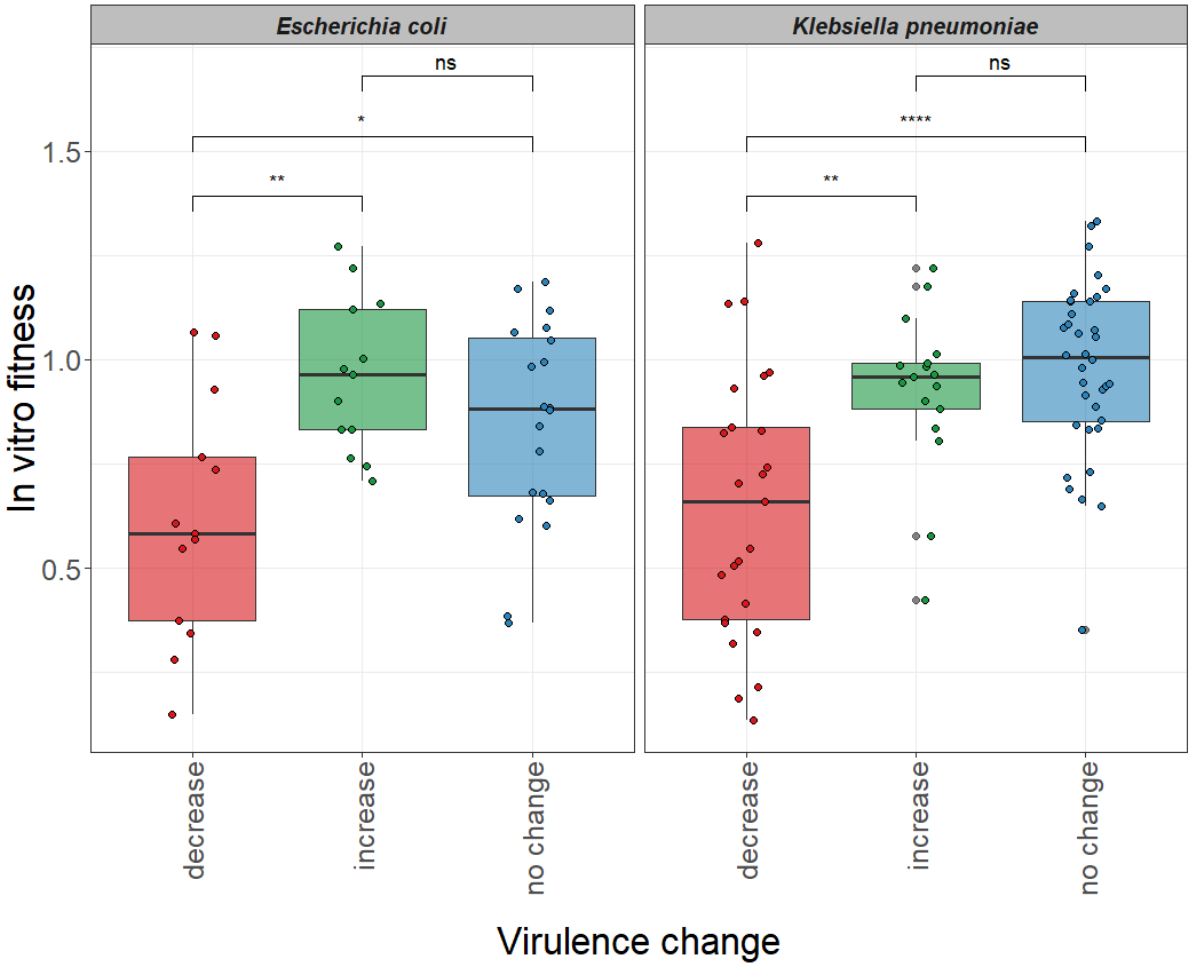


**Figure S5. *In vitro* relative fitness and virulence level.** Each boxplot represents the relative fitness values displayed within one virulence change category compared to the ancestor in the *G. mellonella* infection model (decreased virulence, increased virulence, no change in virulence). Individual points represent the mean relative fitness value of a given antibiotic-adapted line. Fitness was approximated as the area under the growth curve measured in the absence of antibiotics, normalized to the ancestor. While adapted lines with decreased virulence generally display lower relative fitness compared to the other two categories, there is no significant difference between lines with increased and unchanged virulence (Wilcoxon rank-sum test, **** $P \leq 0.0001$, ** $P \leq 0.01$, * $P \leq 0.05$).

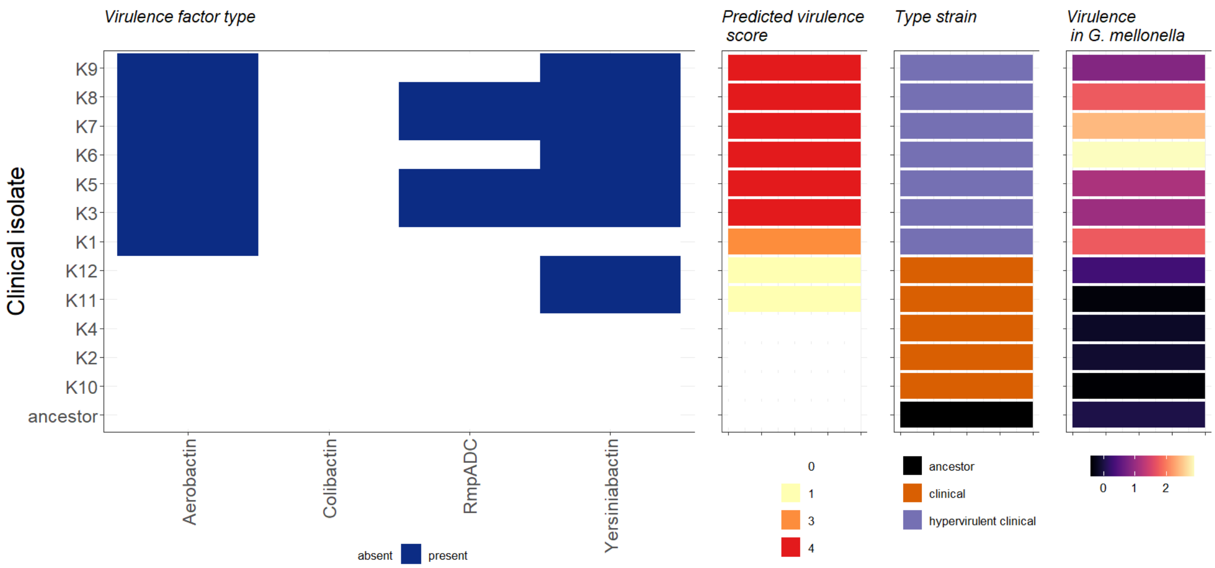


**Figure S6. Virulence factor genes characteristic of the clinical isolates of K. pneumoniae.** Hypervirulent clinical strains were identified based on the presence of virulence factor genes, utilizing the Kleborate scoring system[17]. The figure displays the presence of key virulence factor gene families included in this framework (aerobactin, colibactin, yersiniabactin), together with the RmpADC locus, alongside the corresponding Kleborate-derived virulence scores[47]. In this work, clinical isolates with a virulence score ≥2 were classified as hypervirulent. This classification is consistent with the virulence values measured in *G. mellonella*, as demonstrated in the two rightmost panels.

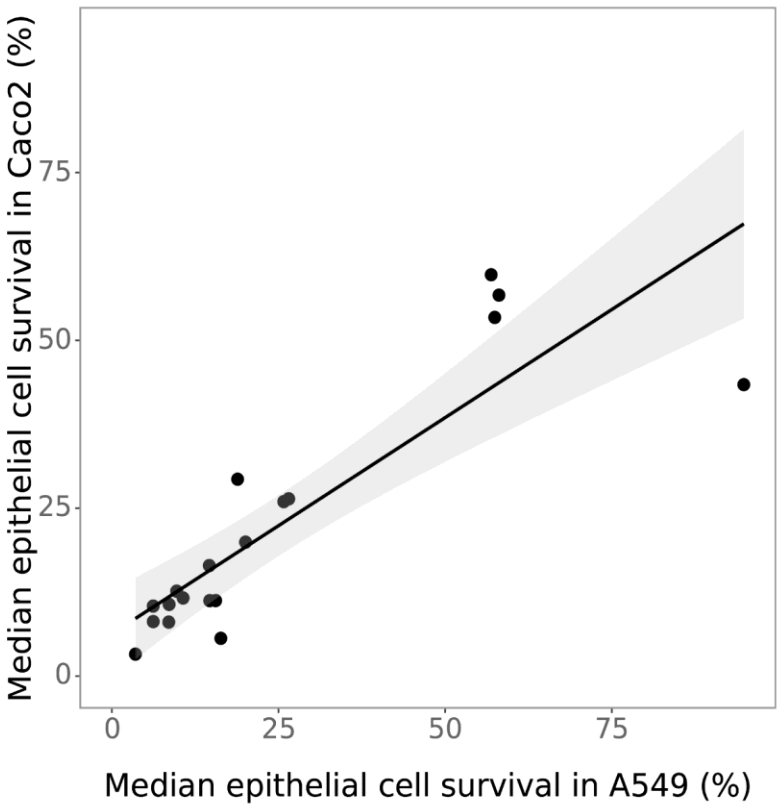


**Figure S7. Correlation of strain-dependent epithelial cell survival between A549 and Caco-2 cells**. Median epithelial cell survival (%) was calculated for each strain in A549 and Caco2 cells and plotted as paired medians (Caco2 vs A549). A least-squares linear regression (Caco2 ~ A549) was performed across strains. The fitted regression line with 95% confidence interval is shown. A strong positive correlation was observed between the two epithelial cell lines ($R^2$ = 0.75, $P = 1.81 \times 10^{-6}$, Pearson), indicating highly concordant strain-dependent effects on epithelial cell survival across cell lines.

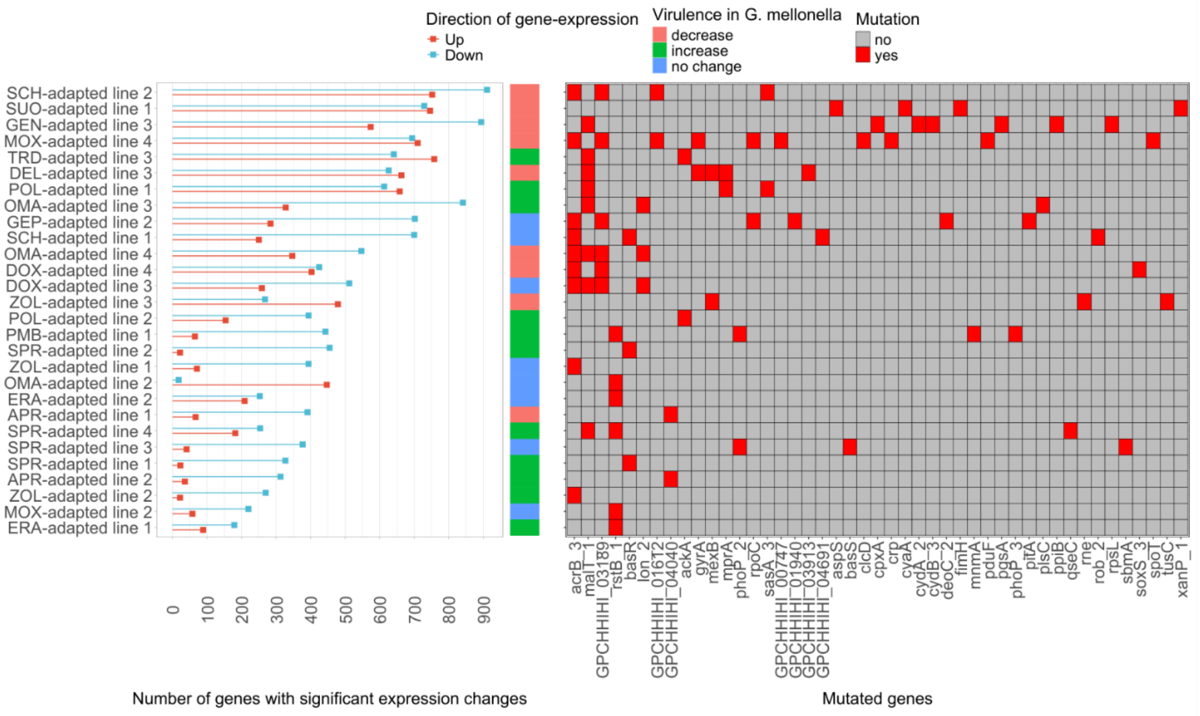


**Figure S8. Differential gene expressions across antibiotic-adapted lines and associated virulence phenotypes.** The figure shows the number of genes with significant expression changes (|log2-fold change| ≥ 1, FDR- adjusted p-value < 0.1) in independently evolved antibiotic-adapted lines. Each row corresponds to a single adapted line. Red squares indicate the number of significantly upregulated genes, while blue squares indicate the number of significantly downregulated genes relative to the ancestor. Annotation bar indicate the direction of change in virulence phenotype in *G. melonella* relative to the ancestor (increase, decrease, or no change). The heatmap on the right shows the presence of mutated genes (red) across the adapted lines, including non-synonymous mutations and indels.

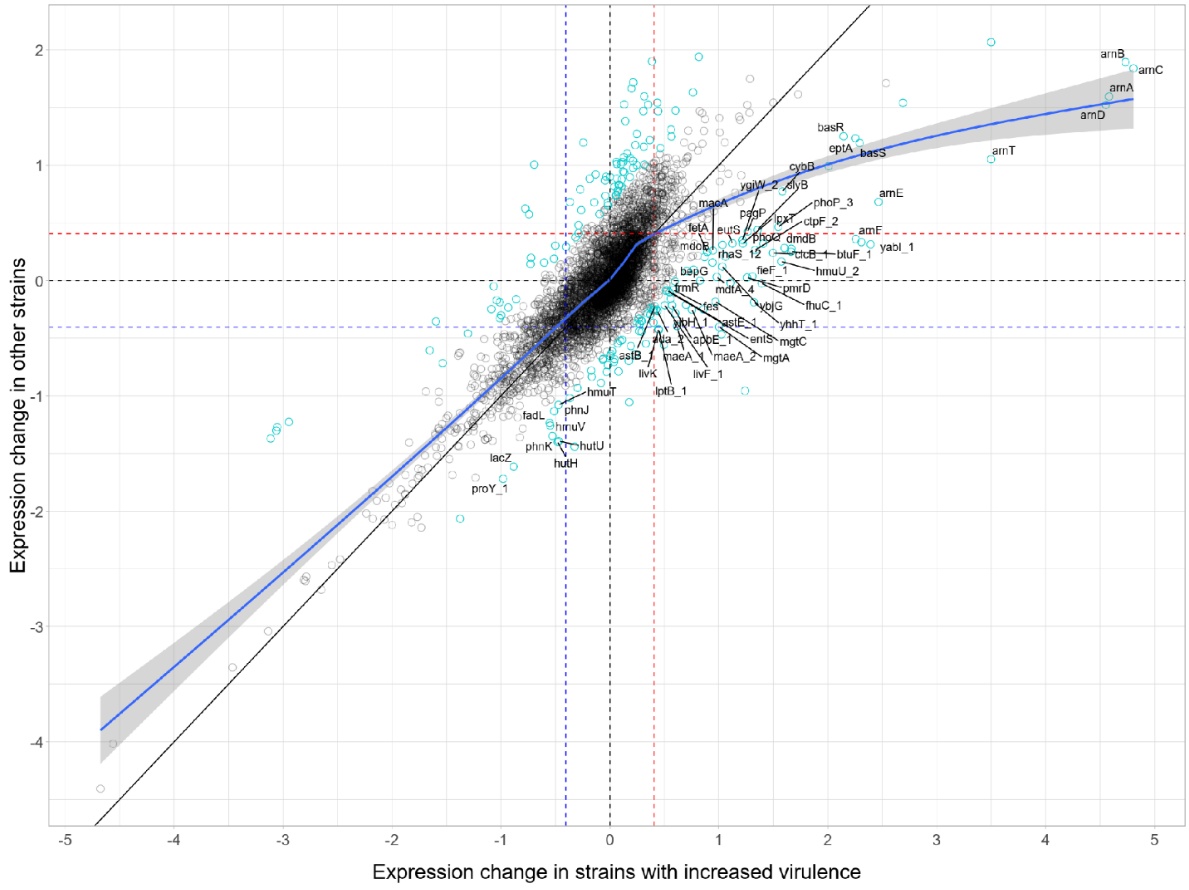


**Fig. S9. Exploratory expression outliers associated with increased virulence in antibiotic-adapted lines.** Scatter plot comparing, for each gene, the mean $\log_2$-fold expression change in antibiotic-adapted lines with increased virulence (x-axis, N = 11) and in lines without increased virulence (y-axis, N = 17), with all fold changes calculated relative to the corresponding ancestor. Each point represents one gene. Cyan points indicate exploratory outliers selected by a combined criterion of regression residual deviation (|residual z-score| > 1.5) and between-group expression divergence (|difference z-score| > 2); grey points represent non-outliers. The residual z-score was calculated after regression of the mean for lines without increased virulence on the increased-virulence mean, and the difference z-score was calculated from the difference between the two group means across genes. This outlier definition is distinct from edgeR differential-expression testing of individual adapted-line-versus-ancestor contrasts and was used to identify genes with large group-level deviations from the shared transcriptional trajectory (55 genes, Data S4). As a complementary line-level analysis, gene-wise Wilcoxon tests compared $\log_2$-fold changes between increased-virulence and non-increased-virulence adapted lines; 17 of the 55 outliers showed nominal evidence of group differences ($P < 0.05$, Data S4). The solid diagonal indicates equal expression change in both groups (y = x), and the blue smoothing curve shows the overall trend (loess regression). The strong positive correlation between the two axes (Spearman's $\rho = 0.79$, $P < 0.0001$) indicates a shared transcriptional response to antibiotic adaptation. Black dashed horizontal and vertical lines mark zero change, while colored dashed lines indicate $\pm\log_2(1.5)$ fold-change thresholds. Genes with the strongest group-specific deviations are labelled, focusing on those with elevated expression in increased-virulence lines. Prominent outliers include the *arnBCADTEF* operon, *eptA*, and the *BasRS/PhoPQ* regulatory systems, as well as genes involved in magnesium transport, ion transport and iron acquisition, including *mgtA, mgtE, hmuU, fhuC* and *fetA*. These genes show systematically higher expression in increased-virulence lines, consistent with enhanced activation of lipid A/LPS modification and nutrient/ion-acquisition pathways.

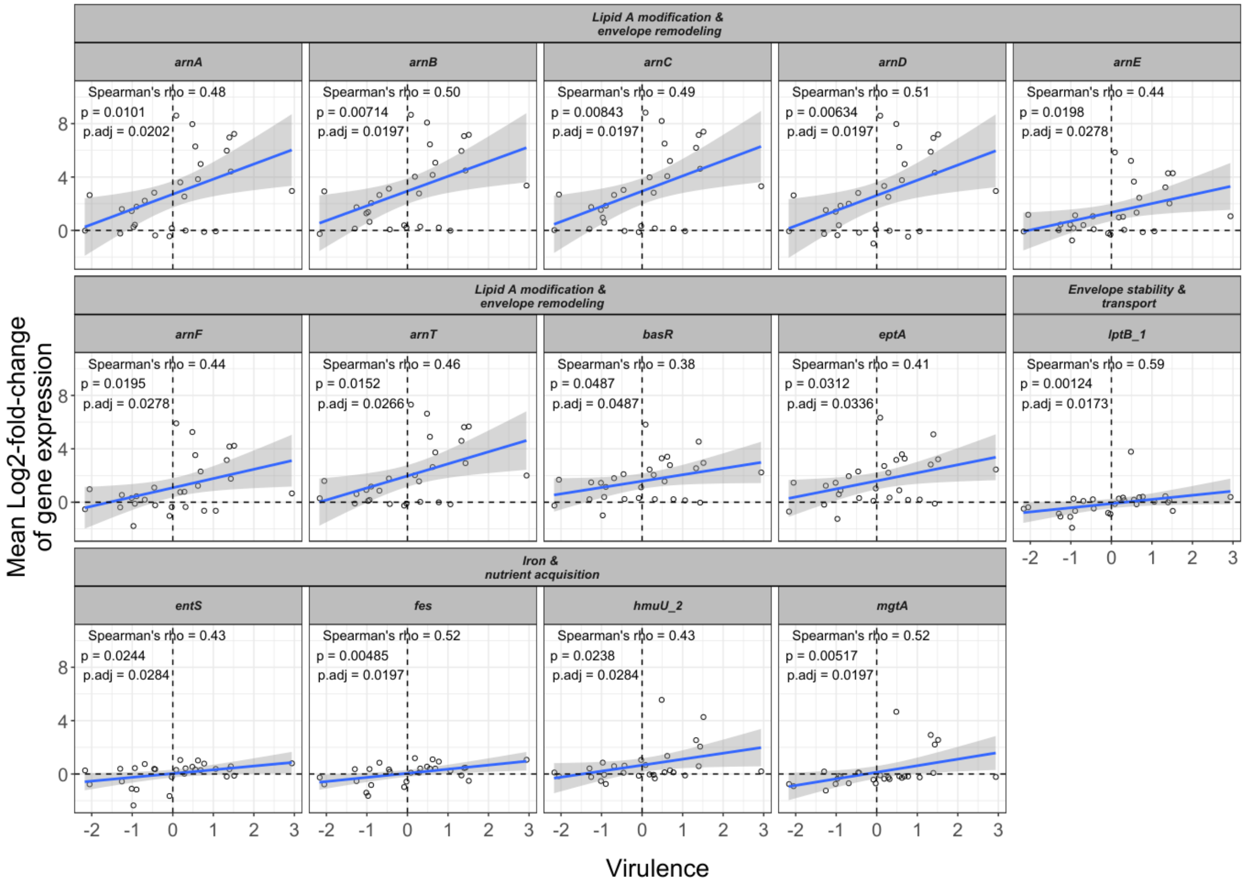


**Figure S10. Coordinated association of key functional modules with virulence across antibiotic-adapted lines.** Scatter plots showing, for selected outlier genes, the relationship between gene expression and virulence across evolved strains. The x-axis represents the continuous virulence measure in *G. mellonella* (relative to the ancestor), and the y-axis shows the corresponding $log_2$-fold change in gene expression. Each point represents one evolved strain. Blue lines indicate linear regression fits with shaded 95% confidence intervals. Dashed horizontal and vertical lines mark zero expression change and no change in virulence, respectively. Reported statistics correspond to Spearman's correlation coefficients (R) and associated nominal and FDR-adjusted *P*-values for each gene. Genes are grouped into non-overlapping functional modules based on curated annotation. The dominant module comprises lipid A modification and envelope remodeling genes, including the *arnBCADTEF* operon, *eptA*, and their regulators *basR*, which together mediate modification of lipid A structure and surface charge. A second module includes a gene involved in LPS transport (*lptB*), whereas the third module consists of genes associated with iron and nutrient acquisition (*hmuU*, *fes*, *entS*), and magnesium transport (*mgtA*), indicating adaptation to nutrient limitation.

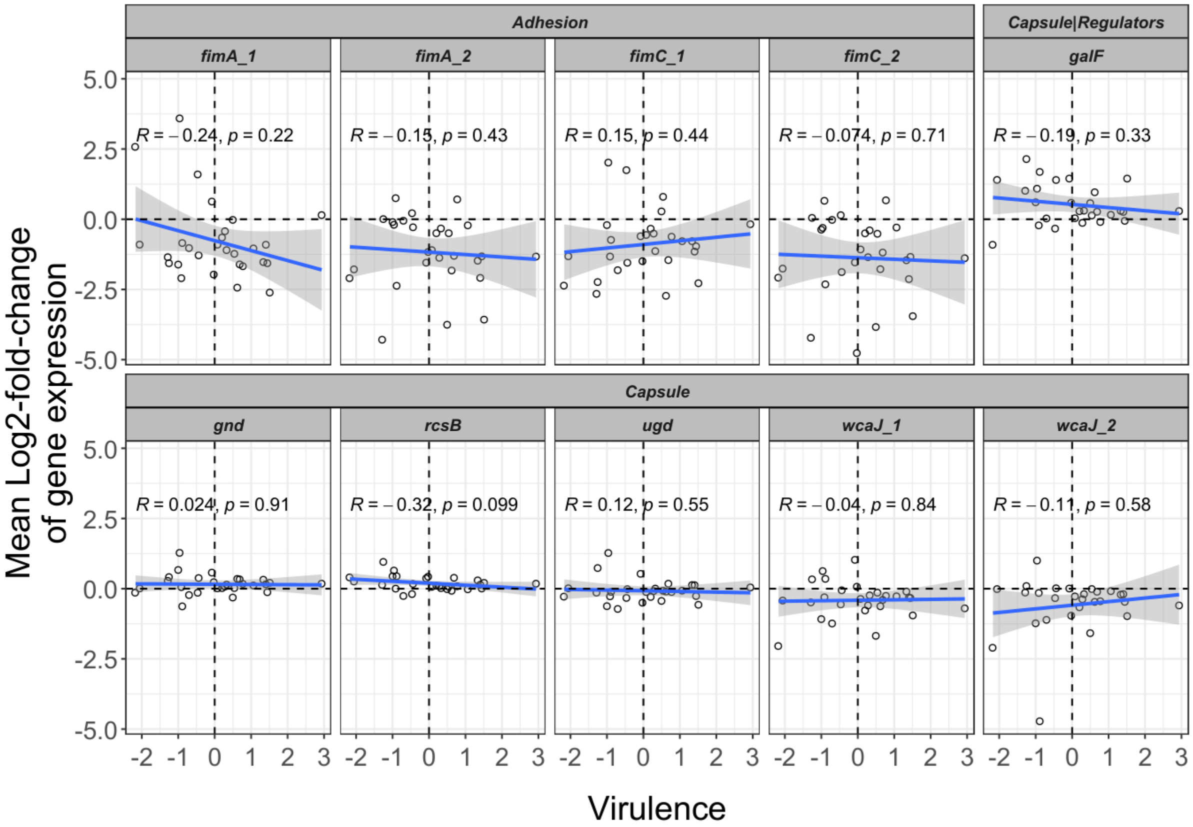


**Figure S11. Lack of coordinated association between capsule and fimbriae gene expression and virulence across antibiotic-adapted lines.** Scatter plots showing, for selected genes associated with adhesion (fimbriae) and capsule biosynthesis/regulation, the relationship between gene expression and virulence across evolved *K. pneumoniae* strains. The x-axis represents the continuous virulence measure in the *G. mellonella* infection model (relative to the ancestor), and the y-axis shows the corresponding $\log_2$-fold change in gene expression. Each point represents one evolved strain. Blue lines indicate linear regression fits with shaded 95% confidence intervals. Dashed horizontal and vertical lines mark zero expression change and no change in virulence, respectively. Reported statistics correspond to Spearman's correlation coefficients (R) and associated P-values for each gene. Genes are grouped into functional categories (adhesion, capsule, and capsule-associated regulators) based on curated annotation of a previous study[48]. Consistent with the aggregate analysis, these genes do not display a coherent or positive association between expression changes and virulence.

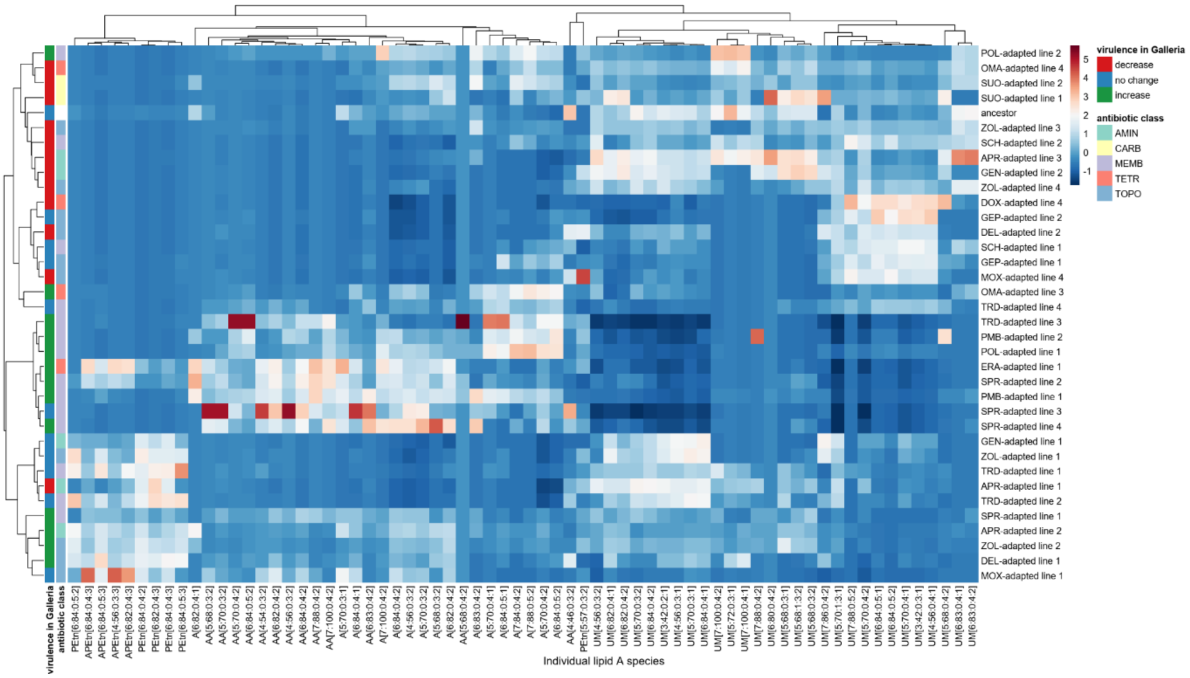


**Figure S12. Heatmap of lipid A composition across antibiotic-adapted *K. pneumoniae* lines.** Rows represent 35 adapted lines, and the corresponding ancestor strain, and columns represent individual lipid A species. Each cell shows the Z-score of the mean relative abundance of the corresponding lipid A species, expressed as a percentage of the total abundance across all identified lipid A species per line. Strains were hierarchically clustered using Spearman distance and average linkage, revealing distinct regions of Lipid A remodeling space. Side annotations indicate cluster assignment, virulence outcome in the *G. mellonella* infection model, and antibiotic class used during experimental evolution.

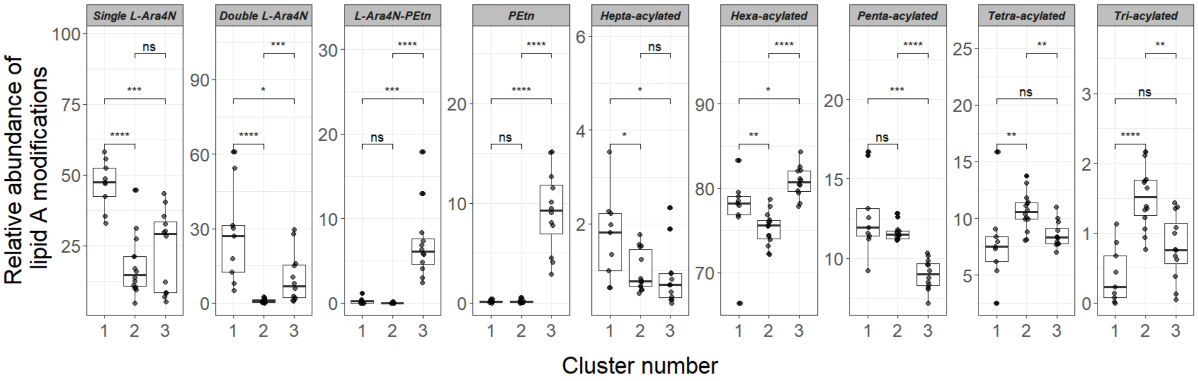


**Figure S13. Relative abundance of lipid A modifications across three clusters of antibiotic-adapted *K. pneumoniae* lines.** Box plots show the relative abundance of lipid A modifications across three clusters identified by hierarchical clustering of 35 antibiotic-adapted lines and the ancestor. Modifications include charge-modifying phosphate substitutions (single and double 4-amino-4-deoxy-L-arabinose (L-Ara4N), phosphoethanolamine (PEtn), and their combination (L-Ara4N–PEtn)) and variation in acyl-chain number (hepta-, hexa-, penta-, tetra-, and tri-acylated). The x-axis indicates cluster identity (1–3). The y-axis shows the relative abundance of each modification, expressed as the percentage of the total identified lipid A signal per line. Each point represents the mean of three biological replicates for an individual evolved line. Boxes indicate the interquartile range with centre lines denoting the median; whiskers extend to 1.5× the interquartile range. Horizontal brackets indicate comparisons between clusters (two-sided Mann–Whitney U test), with significance denoted as ns (not significant), * ($P$ < 0.05), ** ($P$ < 0.01), *** ($P$ < 0.001), and **** ($P$ < 0.0001).

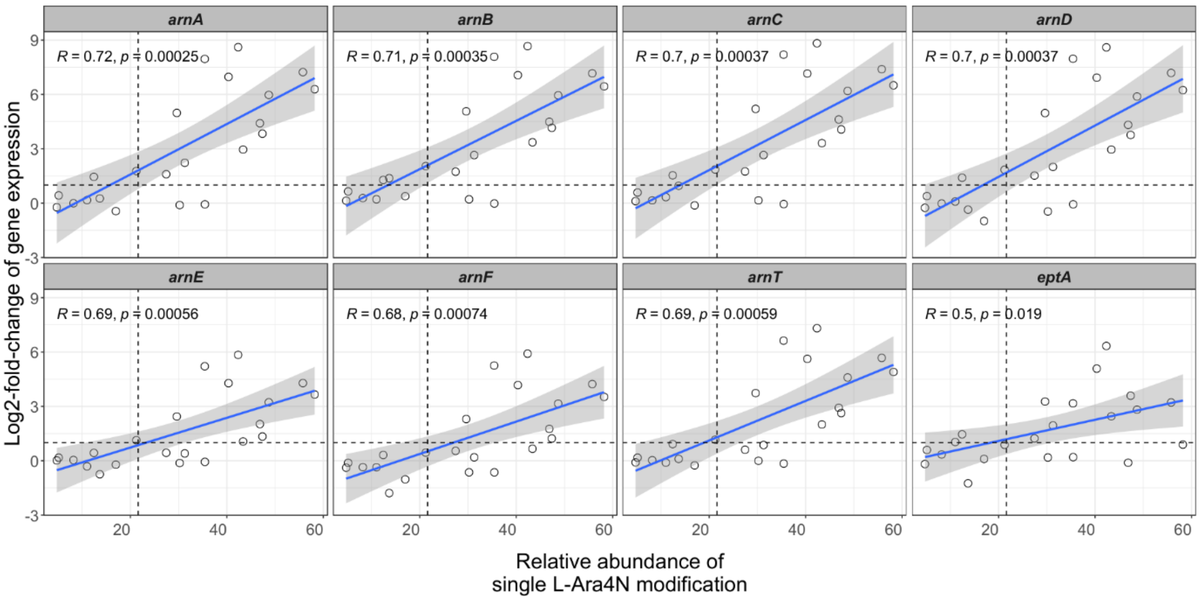


**Figure S14. Expression levels of genes involved in the L-Ara4N modification pathway correlate with L-Ara4N abundance in antibiotic-adapted lines.** Scatter plots show the relationship between the relative abundance of single aminoarabinose-modified lipid A species (L-Ara4N) and the log2 fold change in expression of genes (subpanels) involved in the L-Ara4N lipid A modification pathway (*arnA–F*, *arnT*) and the phosphoethanolamine transferase *eptA*. Each point represents an independently evolved strain grown under antibiotic-free conditions. Solid blue lines indicate linear regression fits with shaded 95% confidence intervals; Spearman's correlation coefficients (R) and associated *P*-values are shown within each panel. Dashed horizontal and vertical lines indicate zero expression change and the reference level of L-Ara4N in the ancestor, respectively. Strong positive correlations for *arnA–F* and *arnT* indicate that increased L-Ara4N modification is associated with upregulation of the *arn* operon, whereas *eptA* shows a weaker association.

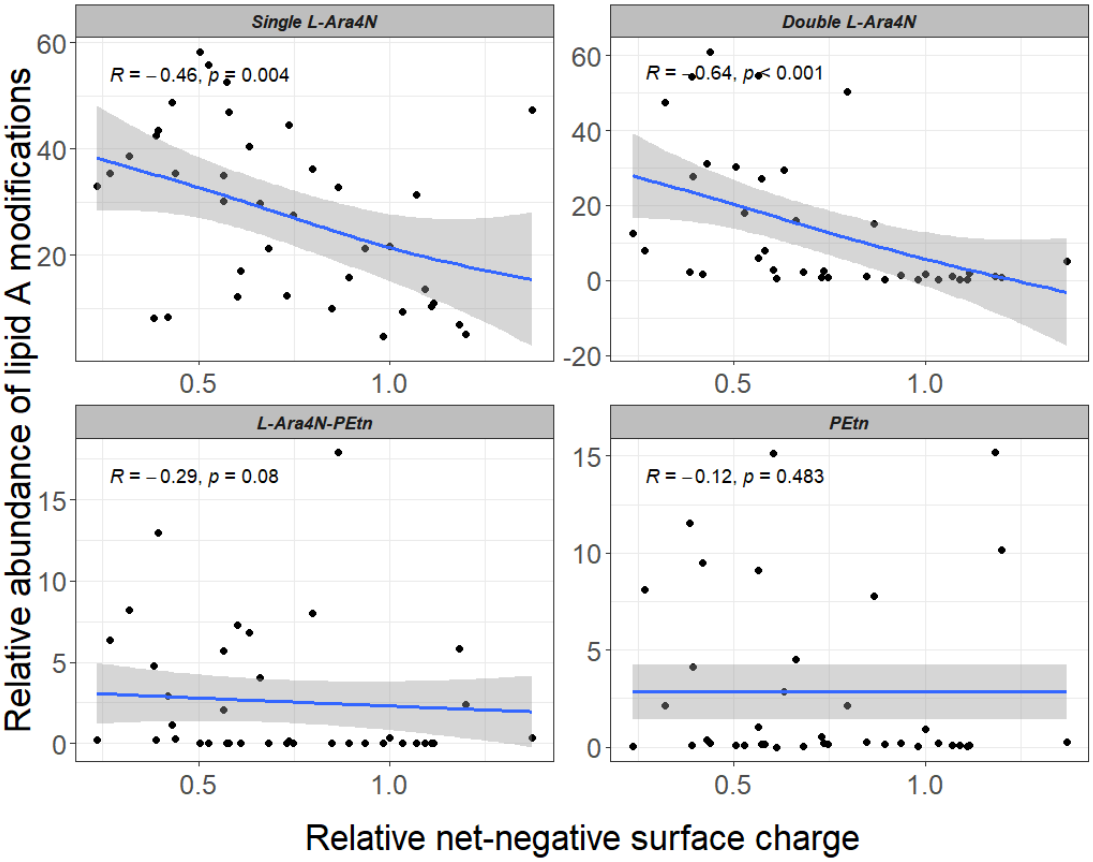


**Figure S15. Single and double 4-amino-4-deoxy-L-arabinose modifications of bacterial lipid A negatively correlate with net-negative surface charge.** The scatterplot shows the relative abundance of lipid A modifications of 35 antibiotic-adapted *K. pneumoniae* lines plotted against the relative net-negative surface charge. Lipid A modifications analysed here include single and double 4-amino-4-deoxy-L-arabinose (L-Ara4N) and phosphoethanolamine (PEtn) charge-modifying phosphate substitutions. Relative abundance of each modification is expressed as the percentage of the total identified lipid A signal per line. Net-negative surface charge was measured using a standard fluorescein isothiocyanate-labeled poly-L-lysine (FITC-PLL) binding assay in triplicate and normalised to the ancestor strain. Each point represents the mean of three biological replicates for an individual evolved line. Each point represents an evolved bacterial line, adapted to different antibiotics. Solid blue lines indicate generalized additive model (GAM) fits with shaded 95% confidence intervals. The association between variables was assessed using Spearman's rank correlation test.

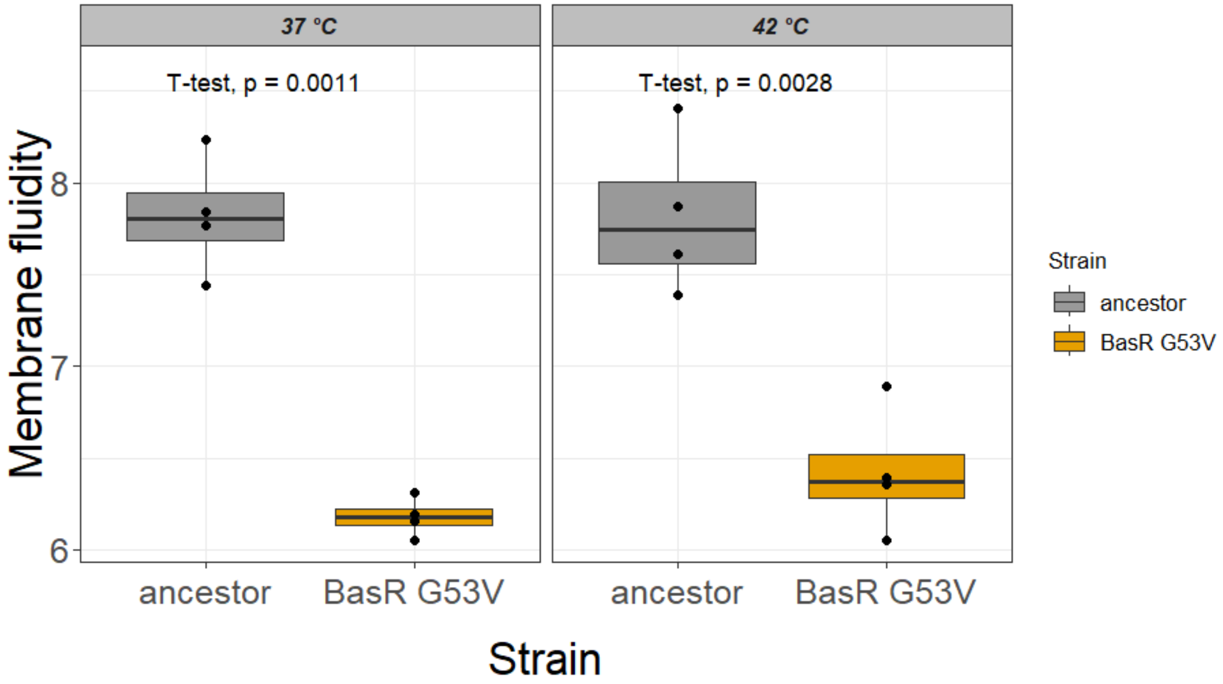


**Figure S16. Membrane fluidity of the BasR G53V mutant and the ancestor of *K. pneumoniae*.** Bulk membrane fluidity was assessed by DPH fluorescence anisotropy in BasR G53V mutant and the corresponding isogenic ancestor strain (*K. pneumoniae* ATCC 10031) at the indicated temperatures. The y-axis shows the inverse of DPH fluorescence anisotropy (1/anisotropy). Box plots indicate the median (centre line), interquartile range (box), and whiskers extending to 1.5× the interquartile range; dots denote independent biological replicates ($n = 4$). $P$ values were calculated using unpaired two-sided Student's $t$-tests.

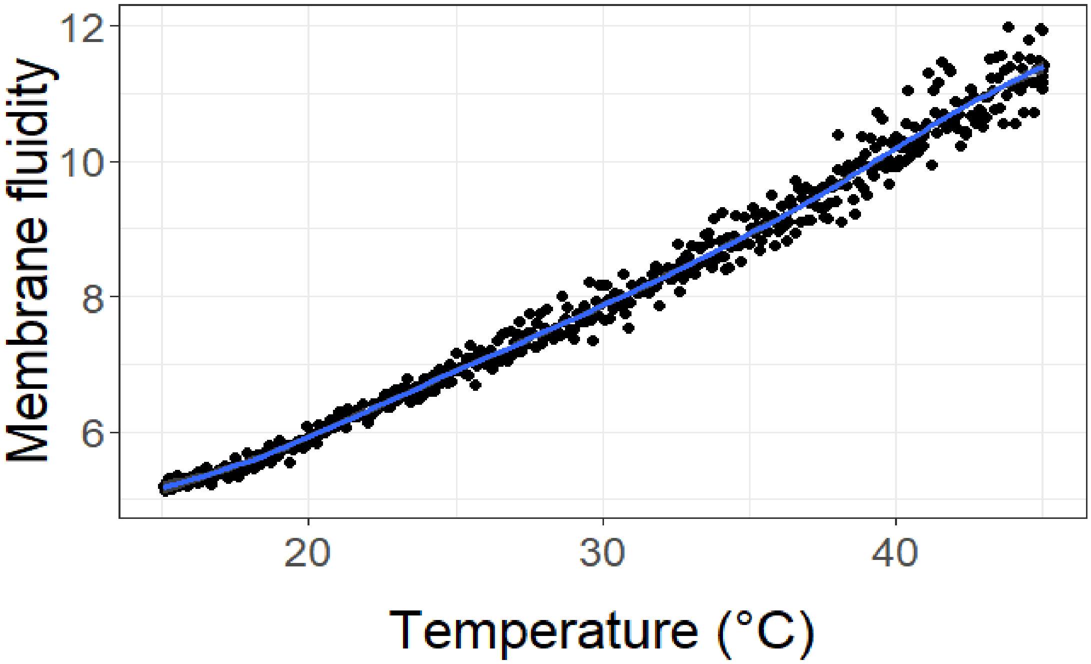


**Figure S17. Temperature dependence of membrane fluidity in lipid vesicles derived from the ancestor of *K. pneumoniae* membrane.** Large unilamellar vesicles (LUVs) prepared from membrane lipid extracts of the *K. pneumoniae* (ATCC 10031) ancestor were used to quantify membrane fluidity by DPH fluorescence anisotropy across a temperature gradient. This enabled estimation of the temperature shift corresponding to the fluidity difference observed between the ancestor and the BasR G53V mutant. The y-axis shows membrane fluidity as the inverse of DPH fluorescence anisotropy (1/anisotropy). Each point represents an individual measurement, and the blue curve indicates the fitted generalized additive model (GAM) describing the relationship between temperature and membrane fluidity.

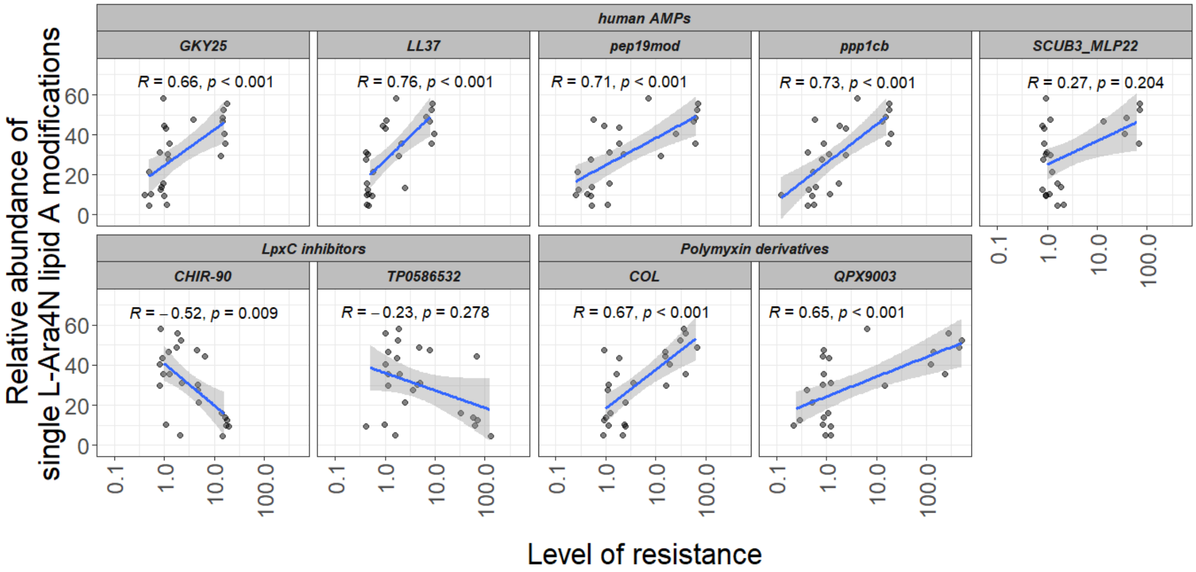


**Figure S18. Association between cross-resistance levels and single L-Ara4N lipid A modification.** Scatter plots show the relationship between the level of resistance (minimum inhibitory concentration fold change relative to the ancestor, log scale) and the relative abundance of single L-Ara4N lipid A modification across *Klebsiella pneumoniae* adapted lines, expressed as the percentage of the total identified lipid A signal per line. Antimicrobial compounds are grouped by class: human antimicrobial peptides (AMPs), LpxC inhibitors, and polymyxin derivatives. Each point represents an independent adapted line. Blue lines indicate linear regression fits with shaded areas representing 95% confidence intervals. Spearman correlation coefficients (R) and corresponding p-values are shown for each compound.

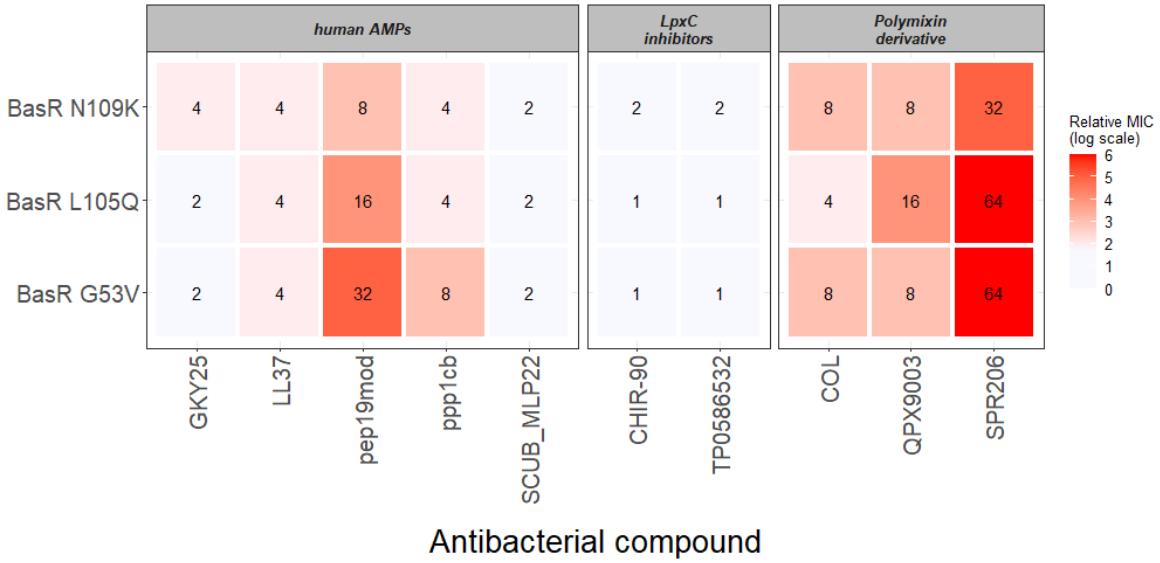


**Fig S19. Cross-resistance profiles of BasR single mutants of *K. pneumoniae*.** The heatmap shows relative minimum inhibitory concentrations (MICs) of 3 BasR single mutants. Susceptibility was determined against selected antimicrobial peptides (AMPs) encrypted in the human proteome and microbiome, polymyxin antibiotics, and LpxC inhibitors. Colors indicate relative MIC values on a $log_2$ scale, normalized to the wild type. The corresponding numerical relative MIC values are shown within each cell.

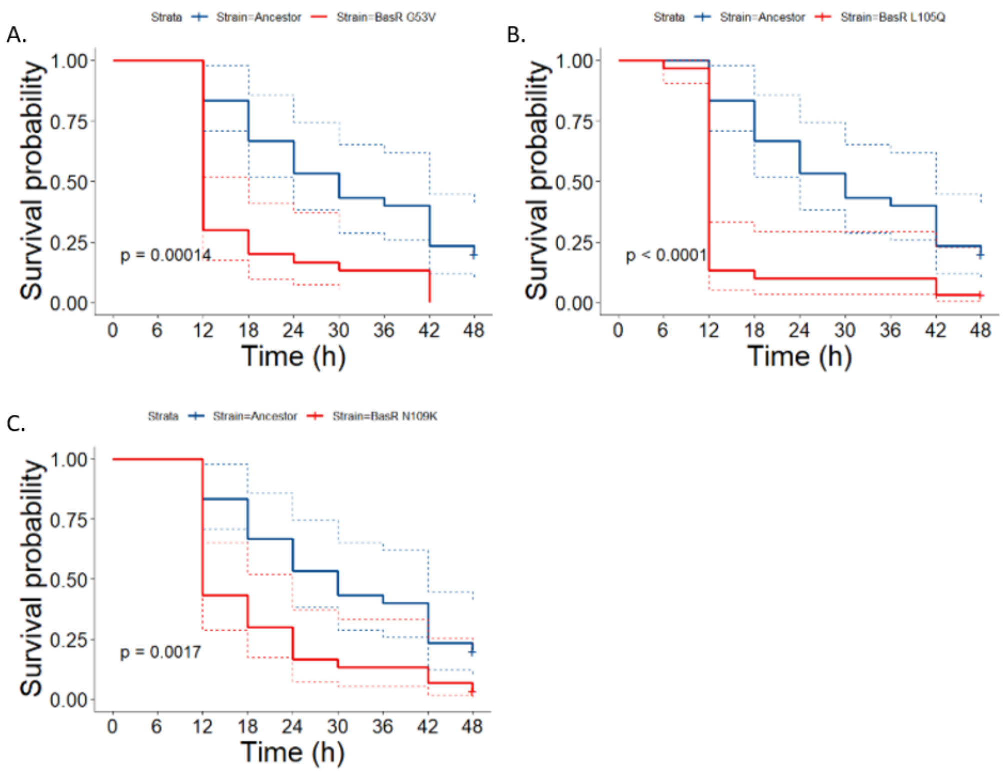


**Fig S20. BasR mutants of *K. pneumoniae* display an increased host-killing rate compared to the ancestor.** Kaplan-Meier survival curves depicting the survival probabilities for *G. mellonella* larvae infected with the ancestor (*K. pneumoniae* ATCC 10031) and three corresponding BasR residue mutants (A: G53V, B: L109Q, C: N105K). Virulence was defined as the host-killing rate in the invertebrate infection model, *G. mellonella,* relative to the corresponding ancestor. In this case, all three BasR residue mutants show increased virulence compared to the ancestor (Likelihood-ratio test). The phosphate-buffered saline (PBS) control group received one injection of sterile phosphate-buffered saline. Experiments were performed in three biological replicates, with 10 animals per treatment group, hence each curve represents 30 animals.

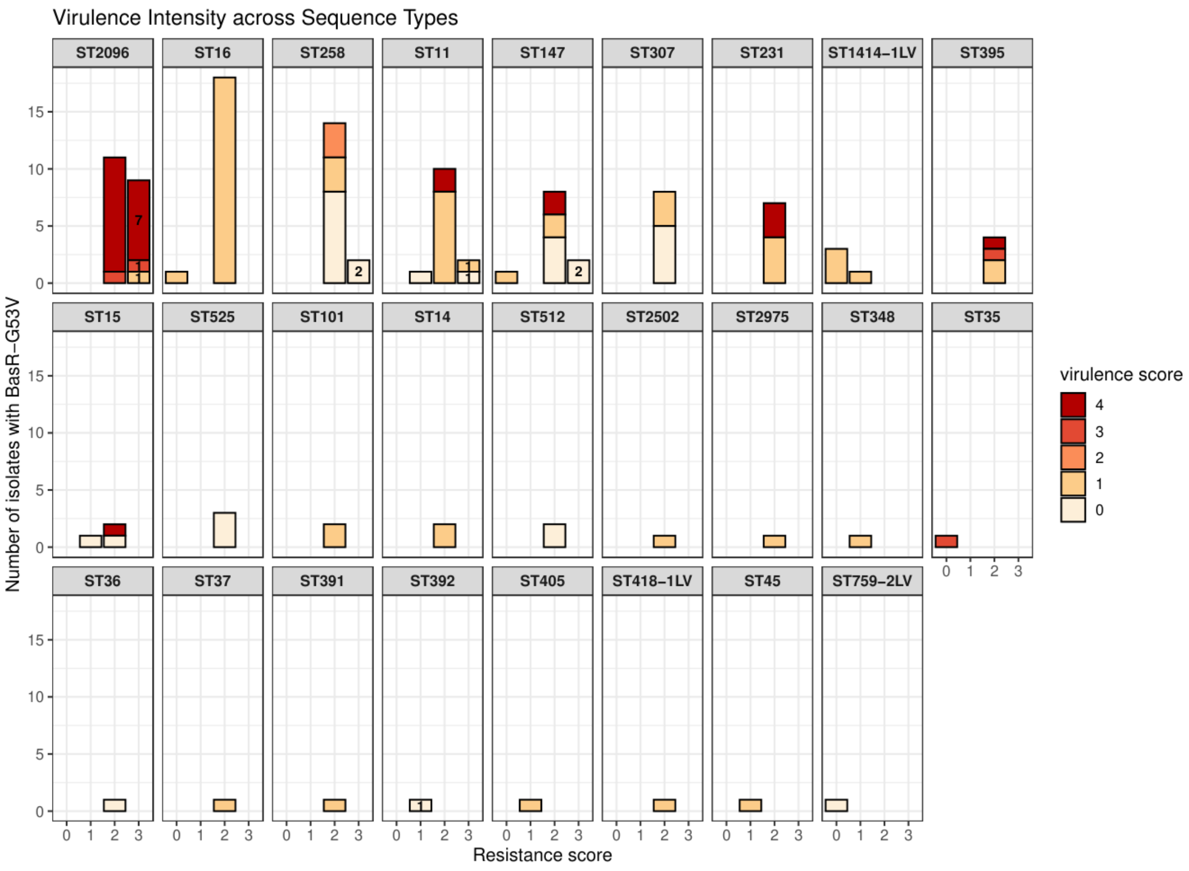


**Figure S21. Distribution of virulence and resistance profiles among natural *K. pneumoniae* isolates harboring the BasR G53V mutation.** Bar plots represent the number of isolates categorized by their resistance and virulence scores across various Sequence Types (STs). Virulence scores (indicated by the color scale) based on genomic data are defined by Kleborate. Resistance scores (X-axis) indicate the presence of critical resistance determinants: 0 (no ESBL or carbapenemase); 1 (ESBL-positive); 2 (carbapenemase-positive); and 3 (carbapenemase-positive with additional colistin resistance). Each facet represents a distinct ST, demonstrating the clonal distribution of these traits. Numerical labels within the colored segments specifically indicate the number of strains resistant to colistin (a last-resort antibiotic), identified within each resistance and virulence category.

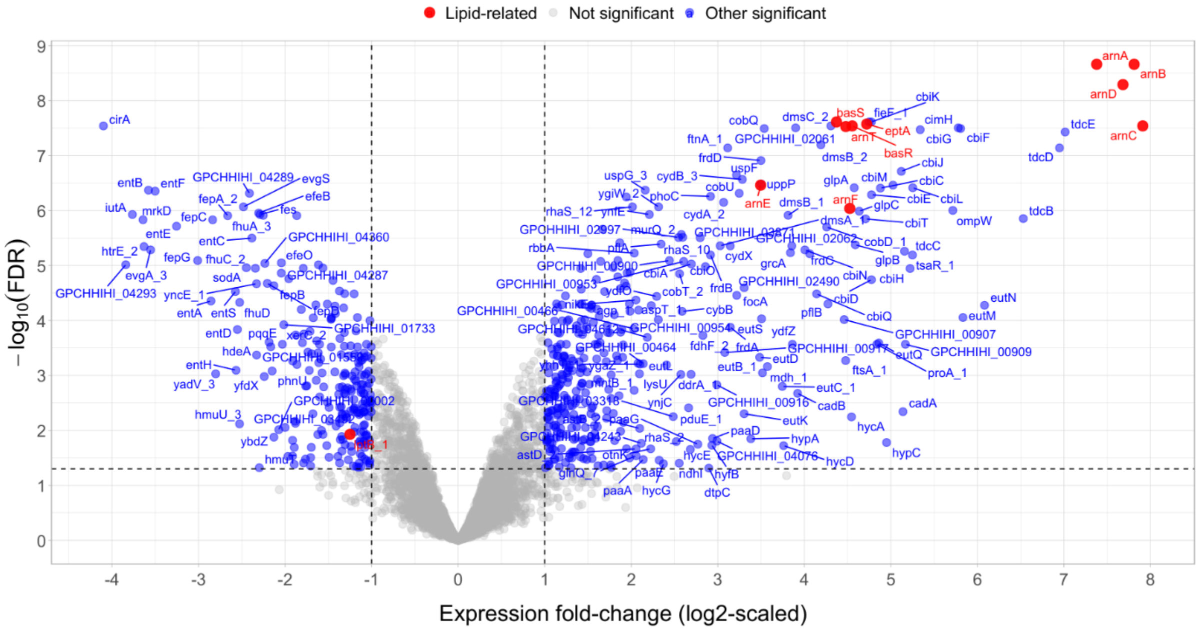


**Figure S22. Differential gene expression highlights LPS-related responses in the BasR G53V mutant *K. pneumoniae*.** The volcano plot shows gene-wise differential expression in the G53V BasR mutant of K. pneumoniae relative to the wild-type, with log2-fold change on the x-axis and $-\log_{10}$(FDR-adjusted P-value) on the y-axis. Each point represents a gene; LPS-related genes are highlighted in red, while non-LPS-related genes are shown in blue. Dashed vertical lines indicate $\log_2$ fold-change thresholds (±1), and the horizontal dashed line marks the nominal significance threshold (FDR-adjusted $P = 0.05$). In total, 364 genes were upregulated and 230 downregulated as a result of a single mutation. Selected genes with large effect sizes (at least 4-fold change in expression) and/or high statistical significance are labeled. Notably, LPS-related genes (e.g., *arn* operon, *basRS*, *eptA*, table S3) are strongly upregulated, consistent with membrane remodeling responses in the mutant strain.

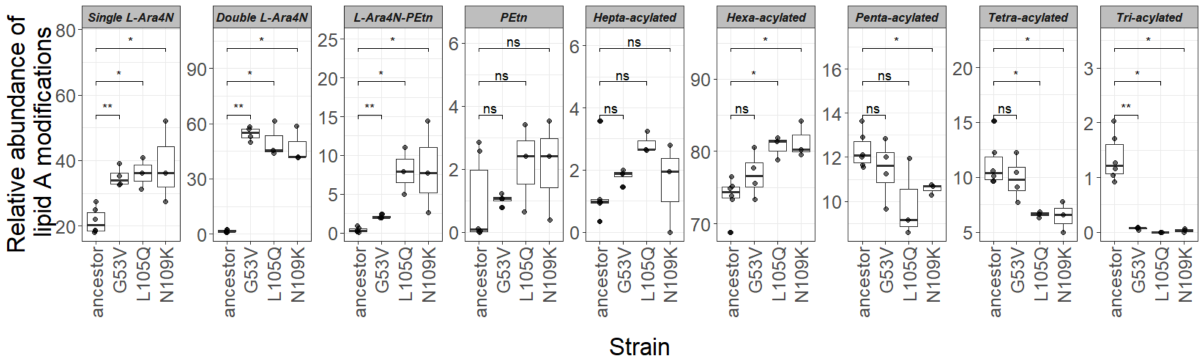


**Fig S23. Single BasR mutations reshape lipid A composition.** Box plots show the relative abundance of lipid A modifications across strains carrying the indicated BasR amino acid substitutions (G53V, L105Q, and N109K) and the isogenic ancestor strain *K. pneumoniae* ATCC 10031. Lipid A modifications include phosphate-group modifications (single and double 4-amino-4-deoxy-L-arabinose (L-Ara4N), phosphoethanolamine (PEtn), and their combination (L-Ara4N–PEtn)) and variation in acyl-chain number (hepta-, hexa-, penta-, tetra-, and tri-acylated). The y-axis shows the relative abundance of each modification, expressed as the percentage of the total identified lipid A signal per sample. Each point represents an independent replicate. Boxes denote the median and interquartile range (IQR), with whiskers extending to 1.5× IQR. Brackets denote Wilcoxon rank-sum tests between strains; significance is annotated as ns (not significant), * ($P < 0.05$), and ** ($P < 0.01$).

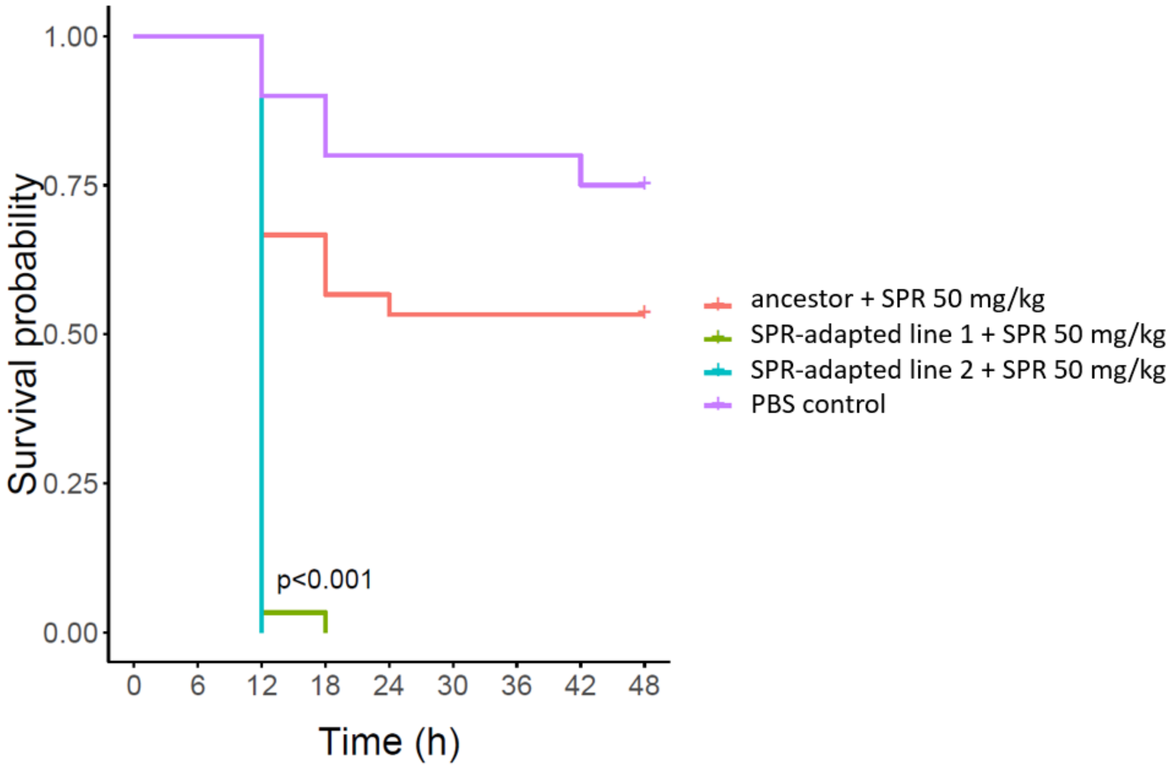


**Figure S24. Infection of *G. mellonella* with SPR206-resistant lines leads to antibiotic treatment failure.** Kaplan-Meier survival curve depicting the survival probabilities for animals infected with the ancestor (*K. pneumoniae* ATCC 1031) and two corresponding SPR206-adapted lines SPR-adapted lines 1 and 2). Each animal received a one-time dose of 50 mg/(body weight kg) SPR206 via injection to the hemocoel within 2 hours of bacterial injection. The phosphate-buffered saline (PBS) control group received two injections of sterile phosphate-buffered saline. While animals injected with the ancestor had approximately 60% survival at the termination of the experiment, 100% of the animals infected with the SPR206-adapted lines perished within 18 hours despite the antibiotic treatment ($P < 0.001$ for both lines 1 and 2; *P*-values were calculated using the standard log-rank test). Experiments were performed in three biological replicates, with 10 animals per treatment group, hence each curve represents 30 animals.

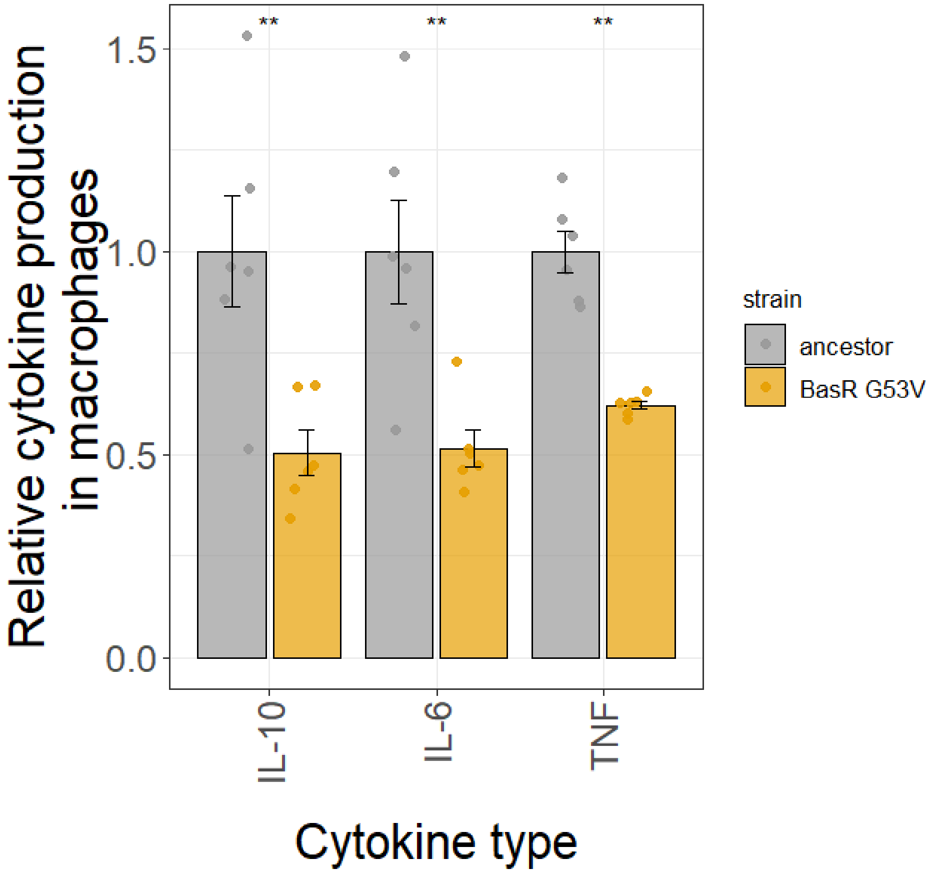


**Figure S25. Cytokine production in macrophages in response to the BasR G53V mutant *K. pneumoniae*.** Macrophages infected with the BasR G53V mutant released lower levels of IL-6, IL-10, and TNF-α compared with ancestor bacteria, as measured by ELISA. Bars show mean values from six biological replicates pooled from two independent experiments; error bars indicate standard error of mean. Cytokine levels are depicted as relative values normalized to the corresponding ancestor within each experiment. Statistical significance was determined using the Wilcoxon rank-sum test (** indicates $P < 0.01$).

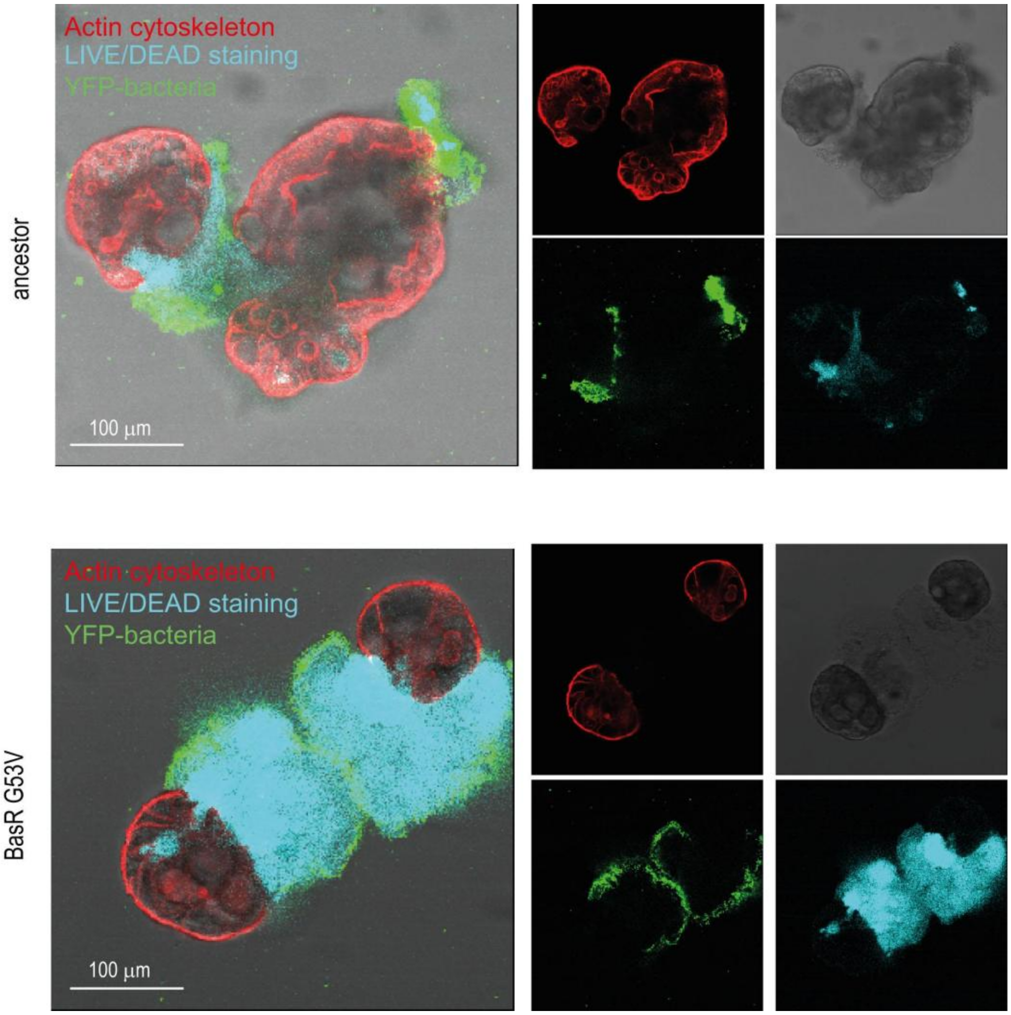


**Figure S26. The BasR G53V mutant *K. pneumoniae* displays enhanced cytotoxicity in apical-out colon organoids.** a, Representative confocal images of apical-out colon organoids derived from healthy human biopsy samples infected with yellow fluorescent protein (YFP) expressing ancestor or BasR G53V *K. pneumoniae* (1h, 37 °C). Organoid cell death was assessed immediately after infection using LIVE/Dead staining. YFP-labeled bacteria (green), dead cells (blue), and the actin cytoskeleton are labeled with CellMask (red), respectively. Scale bar=100 µm.

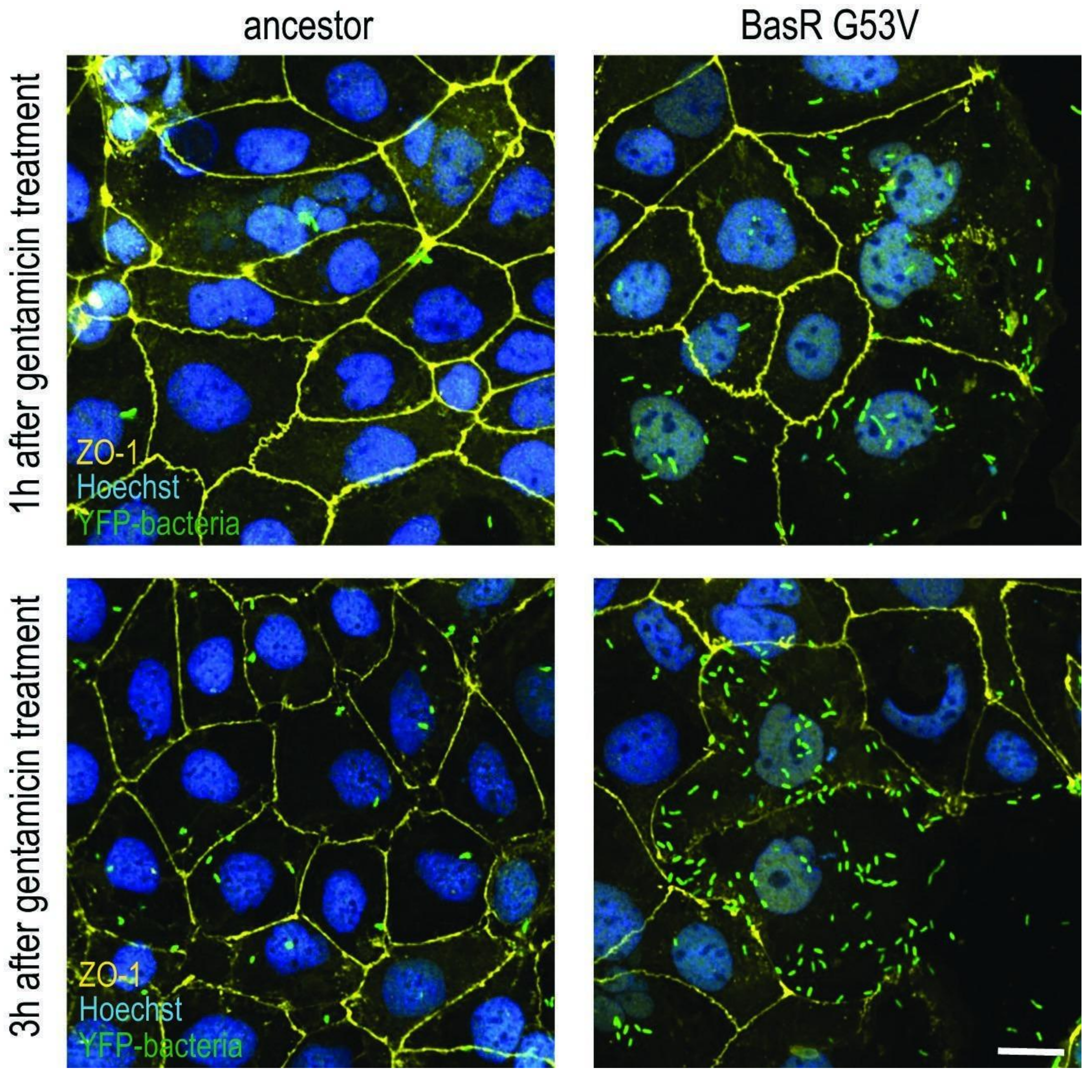


**Fig S27. The BasR G53V mutant *K. pneumoniae* exhibits increased epithelial invasion.** Representative fluorescence images of Caco-2 epithelial cells infected with yellow fluorescent protein (YFP) expressing ancestor or BasR G53V *K. pneumoniae* (green) and subjected to gentamicin treatment to eliminate extracellular bacteria. Tight junctions are visualized by ZO-1 staining (orange), and nuclei are counterstained with Hoechst (blue). Following antibiotic treatment, cells infected with the BasR G53V mutant display more intracellular bacteria compared with the ancestor strain, consistent with increased invasion efficiency. ZO-1 staining delineates epithelial cell boundaries, enabling visualization of bacterial localization within host cells. Images are representative of independent experiments; scale bars = 10 μm, as indicated.

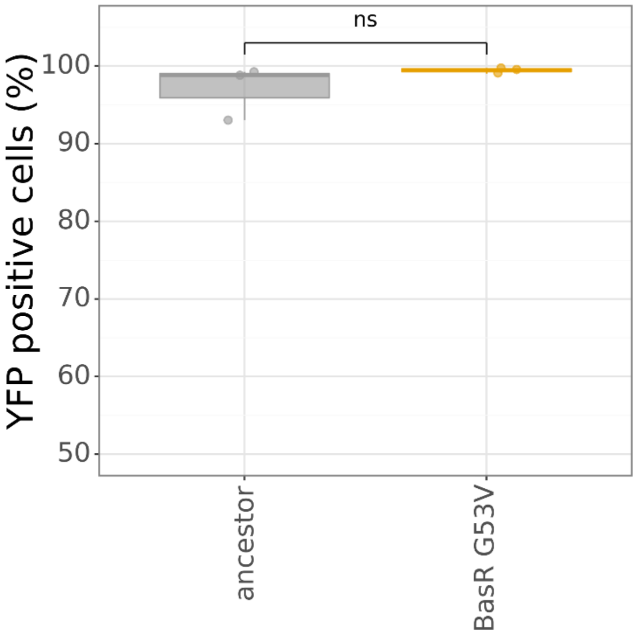


**Figure S28. Validation of fluorescent reporter retention in the ancestor and the BasR G53V mutant *K. pneumoniae*.** The percentage of YFP-expressing cells is comparable between the ancestor and the BasR G53V mutant strains. YFP-expressing bacteria were analysed by flow cytometry, with at least 3,000 bacterial events acquired per sample. The percentage of YFP-positive cells is shown for each strain, demonstrating that the reporter construct is maintained at a similar percentage in both populations. Statistical comparison was performed using Welch's two-sample *t*-test (ns, not significant).

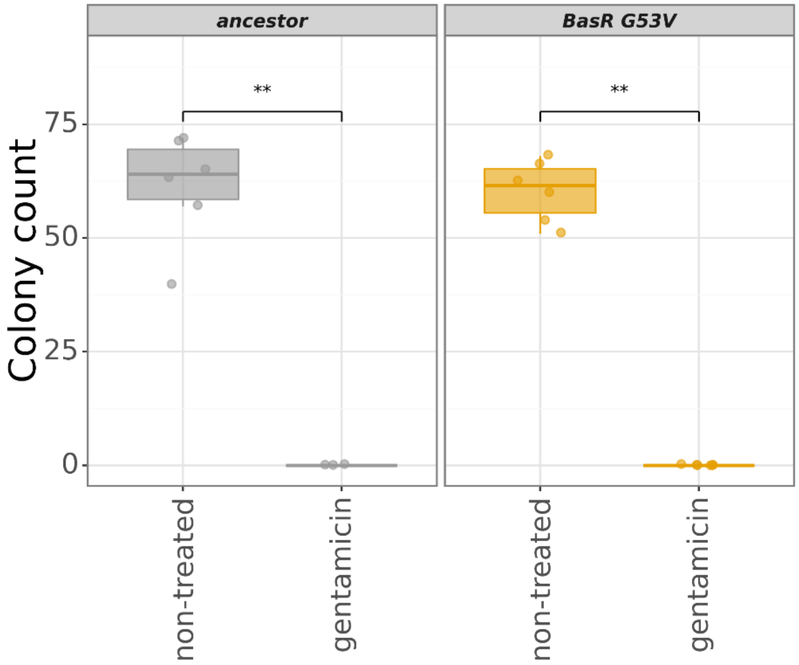


**Figure S29. Validation of antibiotic susceptibility in the ancestor and the BasR G53V mutant *K. pneumoniae*.** The boxplot shows gentamicin susceptibility of the ancestor and BasR G53V strains. Susceptibility was assessed by comparing colony-forming unit (CFU) counts between samples incubated for 1 h in gentamicin-containing or antibiotic-free medium. Within each strain, treated and non-treated samples were compared using a two-sided Mann–Whitney U-test (** indicates $P \leq 0.01$). Both strains showed comparable reductions in CFU following treatment, indicating similar susceptibility and supporting the use of gentamicin protection assays to quantify intracellular bacteria.

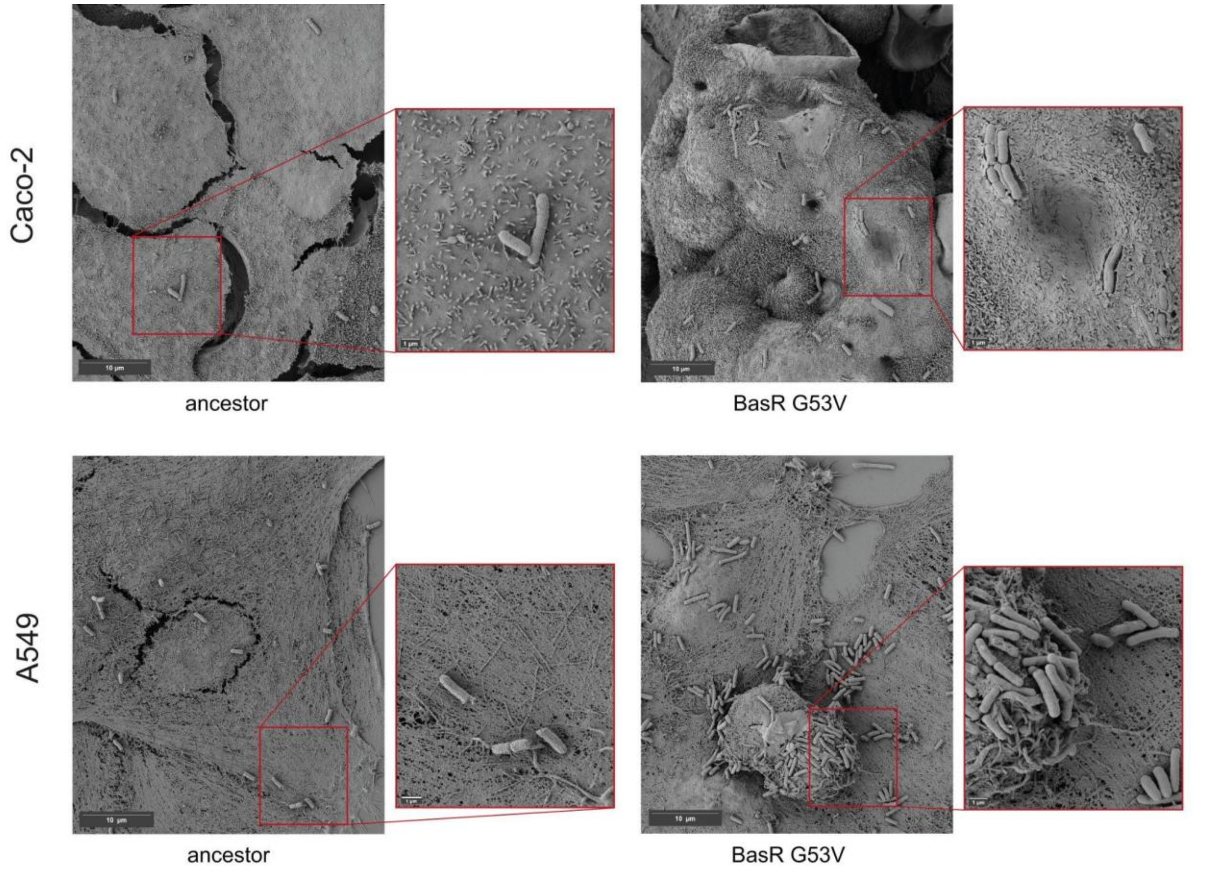


**Figure S30. The BasR G53V mutant disrupts epithelial microvilli and induces membrane damage.** A549 and Caco-2 epithelial cells were infected for 3 h with either the ancestor or the BasR G53V mutant strain and examined by field-emission scanning electron microscopy (FE-SEM). Representative micrographs show extensive membrane damage, loss of apical microvilli, and pronounced surface disruption in epithelial cells infected with the BasR G53V mutant compared with the ancestor strain. FE-SEM imaging of bacteria grown on glass discs further illustrates strain-specific differences in surface morphology and cell–substrate interactions. All samples were fixed, dehydrated through a graded ethanol series, chemically dried with hexamethyldisilazane, sputter-coated with a 12-nm gold layer, and imaged using a Zeiss Sigma 300 FE-SEM. Scale bar = 10 µm. Video S1A and S1B, demonstrating the difference between the ancestor and Bas G53V mutant, respectively, can be found at the following Zenodo link: https://tinyurl.com/5xs5pywc.

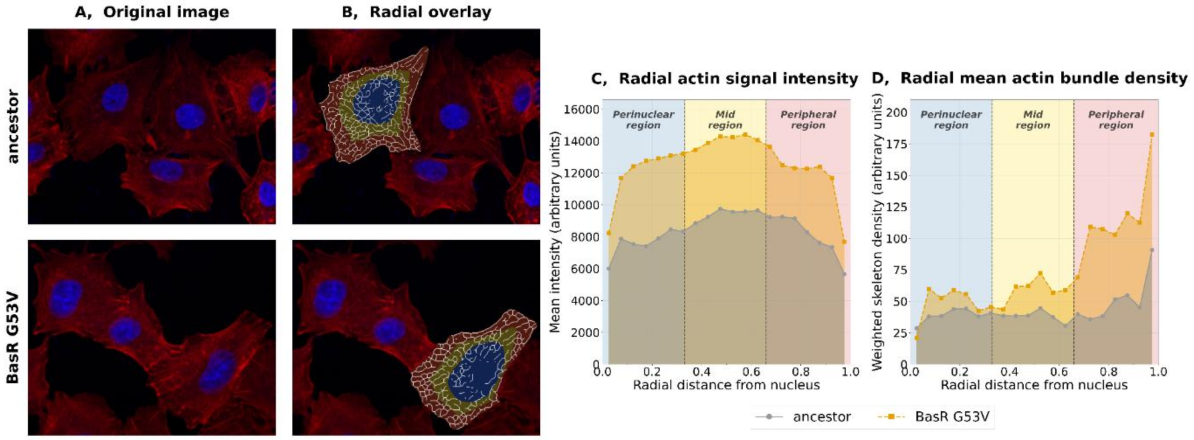


**Figure S31. The BasR G53V mutant *K. pneumoniae* drives peripheral actin remodeling in epithelial cells. A,** Representative images of F-actin (CellMask Deep Red, red) and nuclei (Hoechst, blue) in A549 cells infected with the ancestor or the BasR G53V mutant. **B,** Actin network extraction via Sato tubeness filtering and topological skeletonization. A normalized radial distance r is defined for each pixel, anchored at the nucleus (r = 0) and extending to the cell membrane (r = 1.0), where r corresponds to "Radial distance from nucleus" in panels c and d. Concentric radial zones spanning this axis enable shape-independent spatial comparison across cells of varying size and morphology. **C,** Radial profiles of mean actin signal intensity (arbitrary units, mean raw 16-bit pixel brightness per concentric annulus). The peripheral (red) region (r > 0.66) corresponds to the zone used for quantification. **D,** Radial profiles of mean actin bundle density (arbitrary units, sum of Sato tubeness filter responses along the skeletonized actin network per radial bin, normalized by bin area in pixels). The peripheral (red) region (r > 0.66) corresponds to the zone used for quantification. Compared to the ancestor, the BasR G53V mutant shows higher peripheral actin signal intensity and bundle density, reflecting enhanced membrane-proximal actin polymerization. (For detailed description, see Supplementary Note 6., SN-Fig2-3).

**List of supplementary materials for this manuscript:**

**Video S1A.** Video demonstration of field-emission scanning electron microscopy (FE-SEM) of A549 and Caco-2 epithelial cells after infection with the ancestor strain. All movies are all available at the following Zenodo link: https://tinyurl.com/5xs5pywc.

**Video S1B.** Video demonstration of field-emission scanning electron microscopy (FE-SEM) of A549 and Caco-2 epithelial cells after infection with the BasR G53V mutant strain. All movies are all available at the following Zenodo link: https://tinyurl.com/5xs5pywc.

**Video S2A.** Video demonstration of representative confocal images of murine liver sections at 6 h (A, B) and 24 h (C, D) post-infection with mScarlet-expressing ancestor *K. pneumoniae*. All movies are all available at the following Zenodo link: https://tinyurl.com/5xs5pywc.

**Video S2B.** Video demonstration of representative confocal images of murine liver sections at 6 h (A, B) and 24 h (C, D) post-infection with mScarlet-expressing BasR G53V *K. pneumoniae*. All movies are all available at the following Zenodo link: https://tinyurl.com/5xs5pywc.

**Data S1. (separate file):** Genome identifiers and metadata for ~470,000 genomes of *E. coli* and *K. pneumoniae*.

**Data S2. (separate file):** Various characteristics of antibiotic-adapted lines (resistance, fitness, virulence, surface charge, lipidomics, mutations, epithelial cell survival) or ancestor / BasR G53V mutant strains (cytotoxicity by flow cytometry, organoid cytotoxicity, invasion & adhesion).

**Data S3. (separate file)**: Clinical isolate metadata and associated characteristics.

**Data S4. (separate file)**: Results of RNA-sequencing of *K. pneumoniae* adapted lines and BasR G53V mutant strain.

**Data S5. (separate file):** Literature-derived list of LPS-related genes.

**Data S6. (separate file):** Peripheral actin remodeling raw data (data of 3 experiments).
